# Adaptive Immune Landscape of T-Cell Mediated Rejection of Human Kidney Allografts

**DOI:** 10.1101/2022.05.15.492021

**Authors:** Franco B. Mueller, Hua Yang, Carol Li, Catherine Snopkowski, Darshana M. Dadhania, Jenny Z. Xiang, Steven Salvatore, Surya V. Seshan, Vijay K. Sharma, Manikkam Suthanthiran, Thangamani Muthukumar

**Author notes:** **Corresponding author:** Thangamani Muthukumar, MD, Division of Nephrology and Hypertension, Department of Transplantation Medicine, 525 East 68th Street, Box 3, New York, NY 10065.

## Abstract

The frequently occurring T cell mediated rejection (TCMR) is a risk factor for allograft failure. Immunosuppressive therapy fails to reverse almost 40% of TCMRs occurring in human kidney allografts. A better understanding of the molecular mechanisms of TCMR and precision therapeutics may improve allograft longevity. We investigated adaptive immune landscape of TCMR by genome wide RNA sequencing of 34 prototypic kidney allograft biopsies from 34 adult recipients of human kidney allografts. Sixteen of the 34 biopsies were categorized as Banff TCMR and the remaining 18 as Banff Normal biopsies. Computational analysis identified higher intragraft abundance of the gene sets for key players of adaptive immune system in TCMR. TCMR allografts were characterized by, i) increased antigen processing and presentation and T cell receptor signaling, ii) increased memory T cells, Tregs, Th1, Th2 and Th17 subsets, iii) increased aerobic glycolysis of lymphocytes and reduced metabolic activity of graft parenchymal cells, iv) increased T cell inhibitory receptors and exhaustion markers, v) increased apoptosis and necroptosis, and vi) increased extracellular matrix remodeling, all in comparison to Normal biopsies. Our genome-wide transcriptomics provides an atlas of adaptive immune landscape of TCMR in human kidney allografts, help deduce molecular mechanisms and prioritization of therapeutic targets.

## Introduction

Kidney transplantation is the best treatment option for individuals with irreversible kidney failure. However, immune injury to the transplanted organ—allograft rejection—limits the survival of the transplanted organ and remains a major risk factor for graft failure (1–3). Among the types of allograft rejection, acute T cell mediated rejection (TCMR) is the most frequent type in solid organ transplants. Almost 40% TCMR are not reversed by anti-rejection therapy (4) and recalcitrant TCMR is associated with allograft failure (5).

The anti-allograft immune response is primarily a manifestation of adaptive immunity wherein the processing and presentation of donor antigens by antigen presenting cells (APC) to recipient’s T cells and B cells initiate the cascade immune events culminating in allograft damage and failure unless checked by immunosuppressive drug therapy (6). Because T cells play a non-redundant role in allograft rejection, and B cells contribute to the rejection process by production of antibodies directed at the allograft, T cells and B cells are the primary targets of immunosuppressive drug regimens currently used in organ transplantation. An earlier study from our laboratory (7) suggests that innate immunity may also contribute to allograft rejection.

A better understanding of the molecular mechanisms mediating TCMR may help develop mechanism-based precision therapy. Hybridization-based microarray platforms have been used to interrogate allograft rejection in pre-clinical models and humans and TCMR models have been proposed (8–10). In view of the well-recognized advantages of RNA sequencing over microarray-based profiling, we performed RNA sequencing and bioinformatics analysis to identify genome-wide transcriptional changes in human kidney allograft biopsies classified as TCMR or normal.

## Results

### Study groups, kidney allograft biopsies, RNA sequencing, and genome-wide transcriptional analysis

To characterize genome-wide transcriptional changes during an episode of TCMR in human kidney allografts, we chose 34 kidney allograft biopsies from our biorepository and performed global RNA sequencing and bioinformatics. These 34 biopsies were from 34 unique adult recipients of human kidney allografts and among the 34 biopsies, 16 were prototypic for Banff TCMR (TCMR) and the remaining 18 biopsies were prototypic for Banff Normal/ Nonspecific biopsies (Normal). Demographics of the patients chosen for this investigation, stratified by biopsy diagnosis, are summarized in Supplemental Table S1.

Figure 1A is a heatmap of the Banff kidney allograft biopsy scores. Tubulitis (t) and interstitial infiltration (i) scores were scored on a scale of 0, 1, 2, or 3, and the t and i score of each TCMR biopsy were 4 or greater and fulfilled the Banff histological score criteria for categorization as a TCMR biopsy. Not only t and i scores were zero in each Normal biopsy selected for sequencing but also the scores for peritubular capillaritis and glomerulitis were zero and the biopsies met the Banff histological score criteria for categorization as a Normal biopsy.

**Figure 1.**
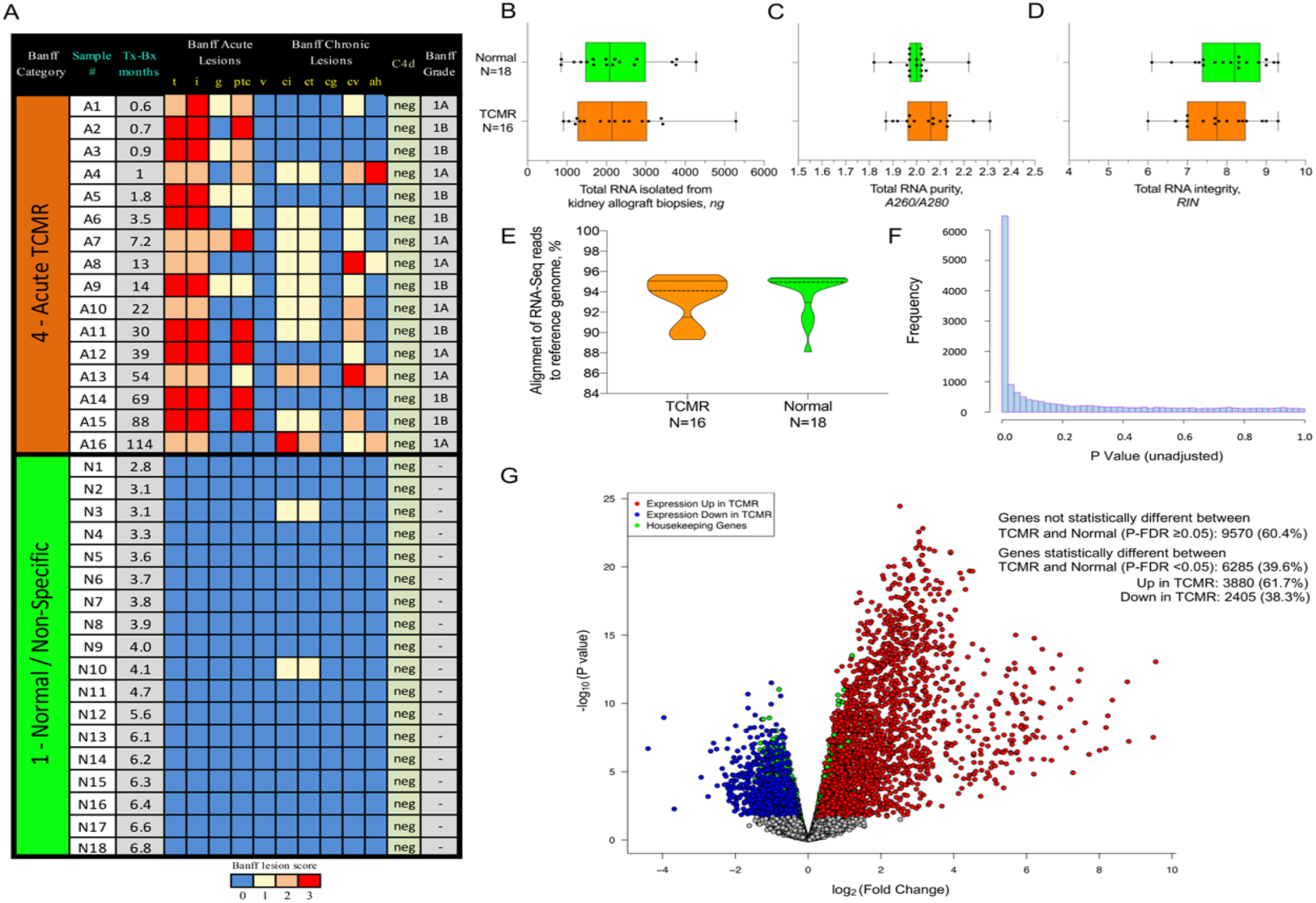
Banff Histopathology Scores, RNA Characteristics, and Volcano Plots of Differential Gene Expression in Human Kidney Allografts. **(A)** Heatmap based on Banff semiquantitative histopathological sores (0, 1, 2, or 3) of the 34 kidney allograft biopsies from 34 kidney allograft recipients (acute T cell mediated rejection [TCMR], samples A1 through A16 and normal/non-specific, samples N1 through N18). The median number of glomeruli were 13 (inter quartile range 9-18) in TCMR and 15 (10–18) in normal/non-specific. All biopsies were negative for peritubular capillary staining for complement factor 4 fragment d (C4d) and for BK virus (SV40 large T antigen). Four of the 16 patients with TCMR had circulating IgG antibodies directed against donor HLA class I (samples A3 and A11) or class II (samples A7 and A10) at the time of biopsy. Donor anti-HLA antibodies were detected using by Luminex single antigen bead assay. Tx-Bx: time from transplantation to biopsy. **(B-D)** Box plots depict the minimum, maximum, and percentile (25th, 50th, and 75^th^) values of the RNA quantity (B), RNA purity (C), and RNA integrity (D) of total RNA isolated from kidney allograft biopsy tissues. The differences in RNA quantity (median 2143ng in TCMR and 2085ng in Normal), purity (2.06 and 2.00), and integrity (7.75 and 8.20) were not statistically significant (P>0.05, Mann Whitney test). **(E)** Violin plot depicts the minimum, maximum, percentile values, and the frequency distribution of the sequence read alignments to the reference human genome (GRCh37/hg19 assembly). The differences in sequence read alignment between TCMR (median 94.10) and Normal (94.95) were not statistically significant (P>0.05, Mann Whitney test). **(F)** Histogram of 16,381 P-values derived by comparing the expression of 16,381 genes between TCMR biopsies and Normal. biopsies Each P-value tests the null hypothesis of no difference in gene expression between TCMR and Normal. If all 16,381 null hypotheses were true, the P-values will follow a uniform distribution between 0 to 1 and the histogram will be flat. If the null hypothesis is not true, there will be greater occurrence of low P-values close to 0. **(G)** Volcano plot depicts the relation between the fold change in intragraft mRNA expression (TCMR vs. Normal) and P-value. Each dot represents an mRNA. Log_2_(Fold Change) value of 0 on the X-axis is equivalent to no difference in gene expression between TCMR and Normal (TCMR gene count/Normal gene count=1). P-FDR is the P value adjusted for false discovery rate.

The box plots show that the quantity of total RNA isolated from the biopsy tissue (Figure 1B), RNA purity (Figure 1C), and RNA integrity (Figure 1D) were not different between TCMR biopsies and Normal biopsies (P>0.05). Figure 1 E, violin plots depict the percentages of alignment of the RNA sequencing reads from each biopsy group to the reference genome GRCh37/hg19 (P>0.05). Figure 1F is a histogram of 16,381 P values, computed by comparing of intragraft abundance of mRNAs in TCMR biopsies vs. Normal biopsies, indicate that the null hypothesis that gene expression is similar between the groups could be rejected. Figure 1G, volcano plot demonstrates differential gene expression between TCMR biopsies and Normal biopsies. Our genome-wide transcriptome analysis of kidney allograft biopsies identified that 6285 (39.6%) of the 16,381 protein-coding genes, including mitochondrial genes, are differentially expressed in TCMR biopsies compared to Normal biopsies at P-adjusted false discovery rate (P-FDR) <0.05. Among the differentially expressed genes, the abundance of 61.7% mRNAs was higher in TCMR biopsies compared to Normal biopsies.

### Intragraft expression of antigen processing and presentation pathway genes

To investigate the expression pattern of antigen processing and presentation pathway gene sets within the allograft during an episode of TCMR, we analyzed intragraft abundance of mRNA encoding: (i) MHC class I and II antigens, (ii) immunoproteasomes—essential for the processing of class I MHC peptides, (iii) heat shock proteins—chaperones involved in signal transduction, (iv) protein folding and degradation, (v) transporters—proteins that pump degraded cytosolic peptides across the endoplasmic reticulum into the membrane-bound compartment where class I molecules are assembled, and (vi) additional proteins such as beta-2 microglobulin (B2M) involved in the MHC class I assembly.

Of the 48 key MHC class I genes, 34 were expressed in the kidney allograft and of the 29 key MHC class II genes, 26 were expressed. Supplemental Figure 1 shows the heatmaps of MHC class I and class II KEGG pathway genes. Supplemental Table 2A lists the median and the IQR of the abundance of mRNAs for class I MHC, stratified by biopsy diagnosis. Among the 26 listed mRNAs, the abundance of 14 were significantly higher and 2 were significantly lower in TCMR biopsies vs. Normal biopsies. The median abundance of all six MHC class I genes (HLA-A/ B/ C/ E/ F/ G) and B2M mRNA were higher in TCMR biopsies. Transporter associated with antigen processing (TAP) is an ATP-binding-cassette transporter family protein that delivers cytosolic peptides into the endoplasmic reticulum where they bind to MHC class I molecules. We identified that the abundance of mRNAs for TAP1 (Fold Change [FC] 4.2, P-FDR 7.3E-12) and TAP2 (FC 3.1, P-FDR 3.1E-12) were higher in TCMR biopsies compared to Normal biopsies.

Supplemental Table 2B lists the abundance of mRNAs for class II MHC pathway genes, stratified by biopsy diagnosis. Of the 26 key class II protein coding mRNAs expressed in the allograft, 20 were significantly different between TCMR biopsies and Normal biopsies and the abundance of all but one, mRNA for NFYC, was higher in TCMR biopsies. Expression levels for IFI30 (GILT) of the endocytosis machinery, and the key MHC class II assembly genes CD74 and CTSS (cathepsin) were higher in TCMR biopsies and all 11 MHC class II genes (HLA-DMA/ DMB/ DOA/ DPA1/ DPB1/ DQA1/ DQA2/ DQB1/ DRA/ DRB1/ DRB5 were also significantly higher. HLA-DO alpha chain mRNA was detected in 8 of the 18 normal allografts and HLA-DO beta chain was not detected in any of the normal allograft samples.

Autosomal minor histocompatibility antigens (mHAgs) are polymorphic cell-derived self-peptides coded by diallelic autosomal genes. mHAgs are displayed on the cell surface by HLA and recognized by HLA class I-restricted CD8+ T cells or HLA class II restricted CD4+ T cells. We identified the 51 genes known to have mHAgs polymorphic sites and measured their intragraft abundance. Of the 51 genes, 46 were expressed in the kidney allografts. Heatmap of expressed mRNAs in each biopsy is shown in Supplemental Figure 2. The median and IQR of the abundance of mRNAs for mHAgs, stratified by biopsy diagnosis, is listed in Supplemental Table 2C. Among the differentially expressed mRNAs, the abundance of 21 mRNAs were significantly higher and 4 lower in TCMR biopsies vs. Normal biopsies. mRNA for HMHA1, a GTPase activator for the Rho-type GTPases, was not detected in the Normal biopsies but was present in 13 of the 18 TCMR biopsies.

### Top enriched adaptative immune system pathways in TCMR biopsies

T cells play a nonredundant role in TCMR and a rejecting allograft is infiltrated by multiple cell types including B cells and macrophages. Fast Gene Set Enrichment Analysis (FGSEA) identified enrichment of T cell receptor signaling pathway genes (Figure 2A) and B cell receptor signaling pathway genes in TCMR biopsies compared to Normal biopsies (Figure 2B). The top eight enriched adaptive immune system pathways (*P*-FDR<0.0001, for each pathway) in TCMR biopsies are shown in Figure 2C. The enriched biological pathways consisted of antigen processing-cross presentation, co-stimulation by CD28 family, downstream TCR signaling, downstream signaling of BCR, endoplasmic reticulum-phagosome pathway, generation of second messenger molecules, PD-1 signaling, and phosphorylation machinery for CD3 and TCR zeta chains.

**Figure 2.**
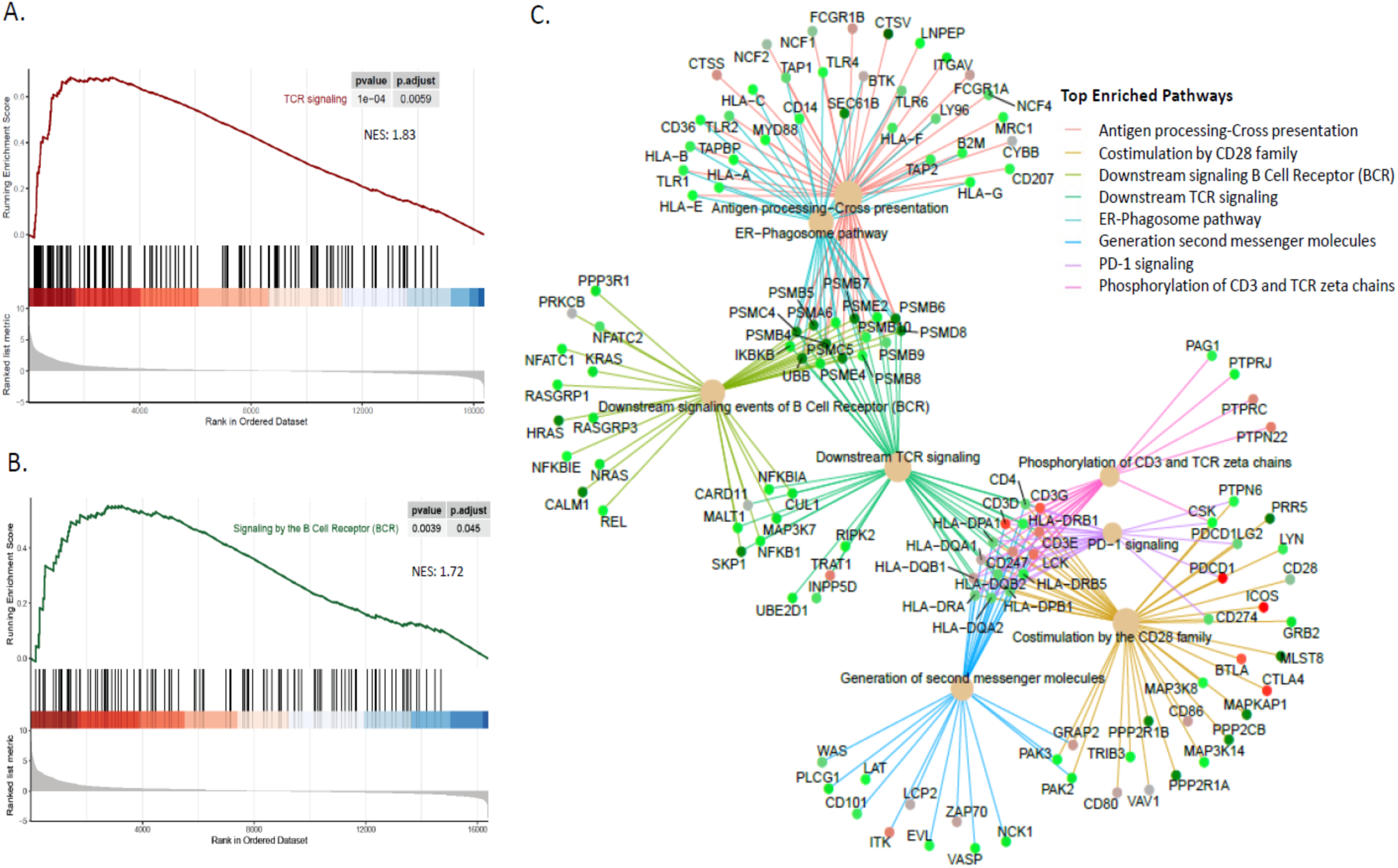
Top Enriched Adaptive Immune System Pathways Based on Intragraft mRNA Abundance: TCMR vs. Normal Biopsies. **(A)** T-cell receptor (TCR) signaling gene set enrichment assessed with FGSEA shows statistically significant, concordant differences between TCMR and Normal. FGSEA is a computational method to determine whether a priori defined set of genes are significantly differentially expressed (each TCMR vs. all Normal) relative to the background whole gene set of 16,381 genes (each TCMR vs. all Normal) that we identified, using a rank-based two-sample T test (equivalent to the non-parametric Mann-Whitney test). The ranked order of differentially expressed genes is shown on the X-axis. The enrichment score reflects the degree to which the TCR signaling gene set is overrepresented at the top of the ranked order of genes. The red line is the enrichment profile of the TCR signaling gene set. The peak of the line is the enrichment score. A positive score indicates that the top-ranked differentially expressed genes are enriched in the TCR signaling gene set. Normalized enrichment score (NES) is shown. The P-FDR is the estimated probability that the gene set with a given NES represents a false-positive finding. The bars in the middle portion of the graph shows where the members of the TCR signaling gene set appear in the ranked list of differentially expressed genes. The bottom portion of the graph shows the correlation of the differentially expressed genes with TCMR (positive) or Normal (negative). **(B)** B-cell receptor signaling gene set enrichment assessed with FGSEA. **(C)** Gene-concept network graph depicts a linkage of genes as a network to extract the complex association in which a gene may belong to multiple annotation categories. Genes shown as colored dots—color representing the fold change value—are the ones that are differentially expressed (P-FDR<0.05) between TCMR and Normal. Top eight of 315 enriched Reactome (database of biological pathways) adaptive immune system pathways are shown. A biological pathway is an ordered series of molecular events that leads to the creation of a new molecular product or a change in a cellular state or process. The statistical significance of the pathways is an over-representation (enrichment) analysis that tests whether a specific pathway is over-represented among the differentially expressed genes—more genes of a particular pathway than would be expected by chance. All eight depicted pathways were statistically significant (P-FDR<0.05).

### Expression of T cell receptor signaling pathway genes

We found that 104 of 112 genes reported to participate in TCR activation and co-stimulation were present in kidney allografts and 64 were differentially expressed between TCMR and Normal biopsies. Of the 73 signal 1 (antigenic signal) genes, 45 were differentially expressed with increased abundance of 42 in TCMR biopsies vs. Normal biopsies (Supplemental Table 3A). Figure 3, box plots, show the higher abundance of the mRNAs for CD3D, CD3E, CD3G, CD247 of the T cell receptor complex (Figure 3A) and for the mRNAs for CD4 and CD8 (Figure 3B). Figure 3C is a heatmap of differentially expressed signal 1 T cell activation genes. The abundance of the three of the four key PTKs—LCK, ITK and ZAP70, and the co-receptor PTPRC (CD45) were also significantly higher in TCMR biopsies.

**Figure 3.**
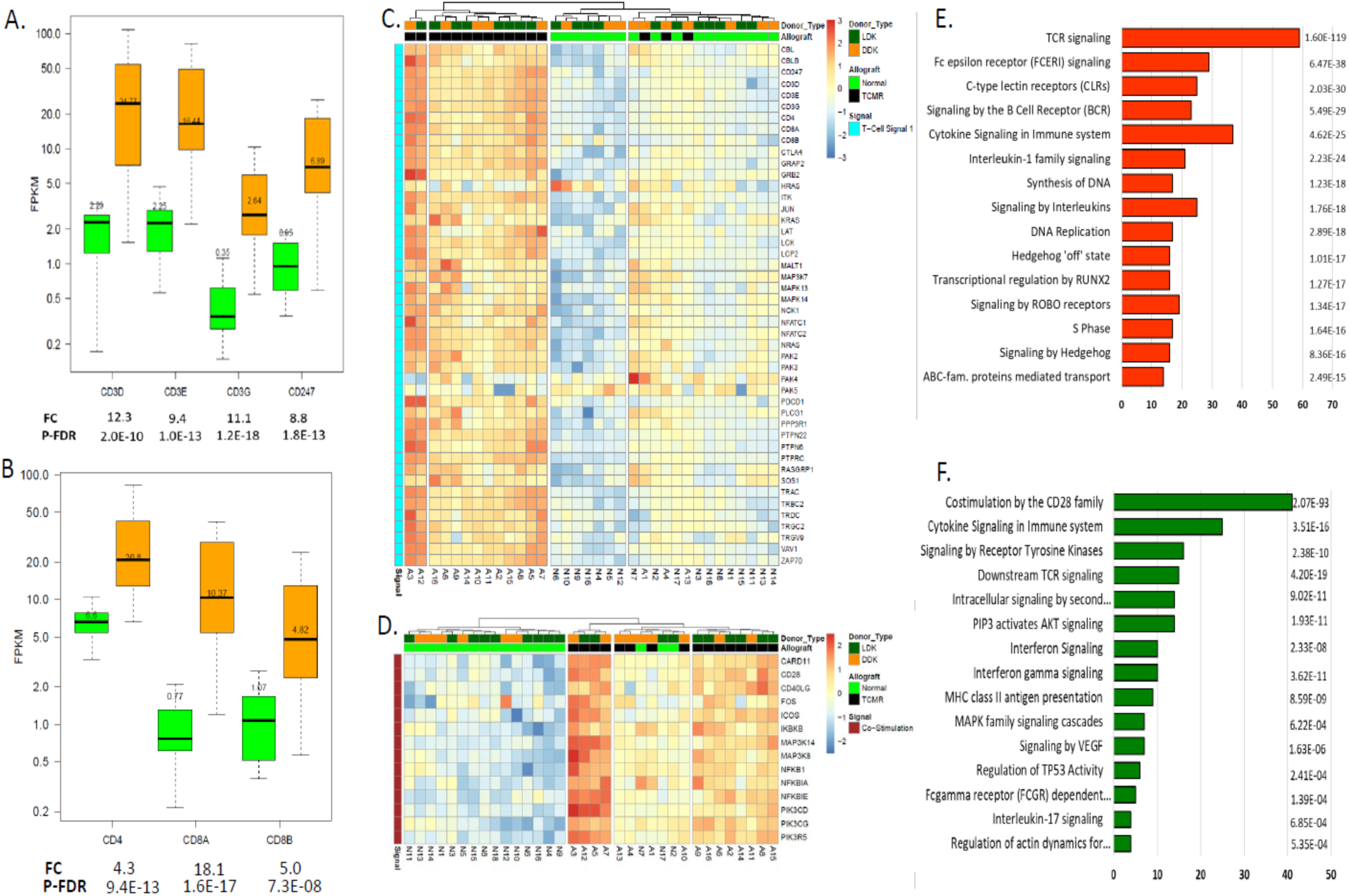
Intragraft abundance of mRNAs for T-Cell Activation Pathways are higher in TCMR biopsies Compared to Normal Biopsies. **(A)** Intragraft abundance of mRNAs encoding T-cell receptor complex proteins in human kidney allograft biopsies (orange: TCMR [N=16]; green: Normal [N=18]). The number within each box plot is the median value of mRNA abundance reported as fragments per kilobase of exon per million reads mapped (FPKM). The fold change (FC) values shown below each mRNA and the P-FDR were calculated using edgeR. **(B)** Intragraft abundance of mRNAs encoding CD4 and CD8A and CD8B T-cell coreceptors. **(C)** Heatmap visualization of hierarchical clustering analysis of intragraft mRNA abundance of genes involved in Signal 1 (antigenic signaling) of T-cells. **(D)** Heatmap visualization of hierarchical clustering analysis of intragraft mRNA abundance of genes involved in Signal 2 (costimulation) of T-cells. **(E)** Top Reactome activation pathways of antigenic signaling T-cells **(F)** Top Reactome activation pathways of costimulation of T-cells.

Among the co-stimulation pathway genes, 27 were expressed in the kidney allograft biopsies and of these, 14 were differentially expressed—all increased in TCMR biopsies. Key genes differentially expressed in the T cell co-stimulation pathway were membrane bound CD28, essential for T-cell proliferation and survival, cytokine production, and T-helper type-2 development; inducible T-cell co-stimulator (ICOS) and CD40LG that regulates B cell functions by engaging CD40 on the surface of B cells (Figure 3D). Figures 3E and F shows the top 15 signal 1-enriched and signal 2-enriched biological pathways. Supplemental Table 3B lists the abundance of co-stimulatory pathway genes, stratified by biopsy diagnosis.

### Expression of T cell subset gene signatures

Multiple T cell subtypes infiltrate a rejecting allograft and we leveraged single-cell RNA sequencing data (11) and investigated graft infiltrating T cell subsets. Figure 4A-C, heat maps and Figure 4D, cumulative distribution function (CDF) plots, demonstrate that the intragraft abundance of mRNAs for CD8^+^ memory T cells (Figure 4A), CD4^+^ memory T cells (Figure 4B), and CD8^+^ γδ mRNAs (Figure 4C) are higher in TCMR biopsies compared to Normal biopsies.

**Figure 4.**
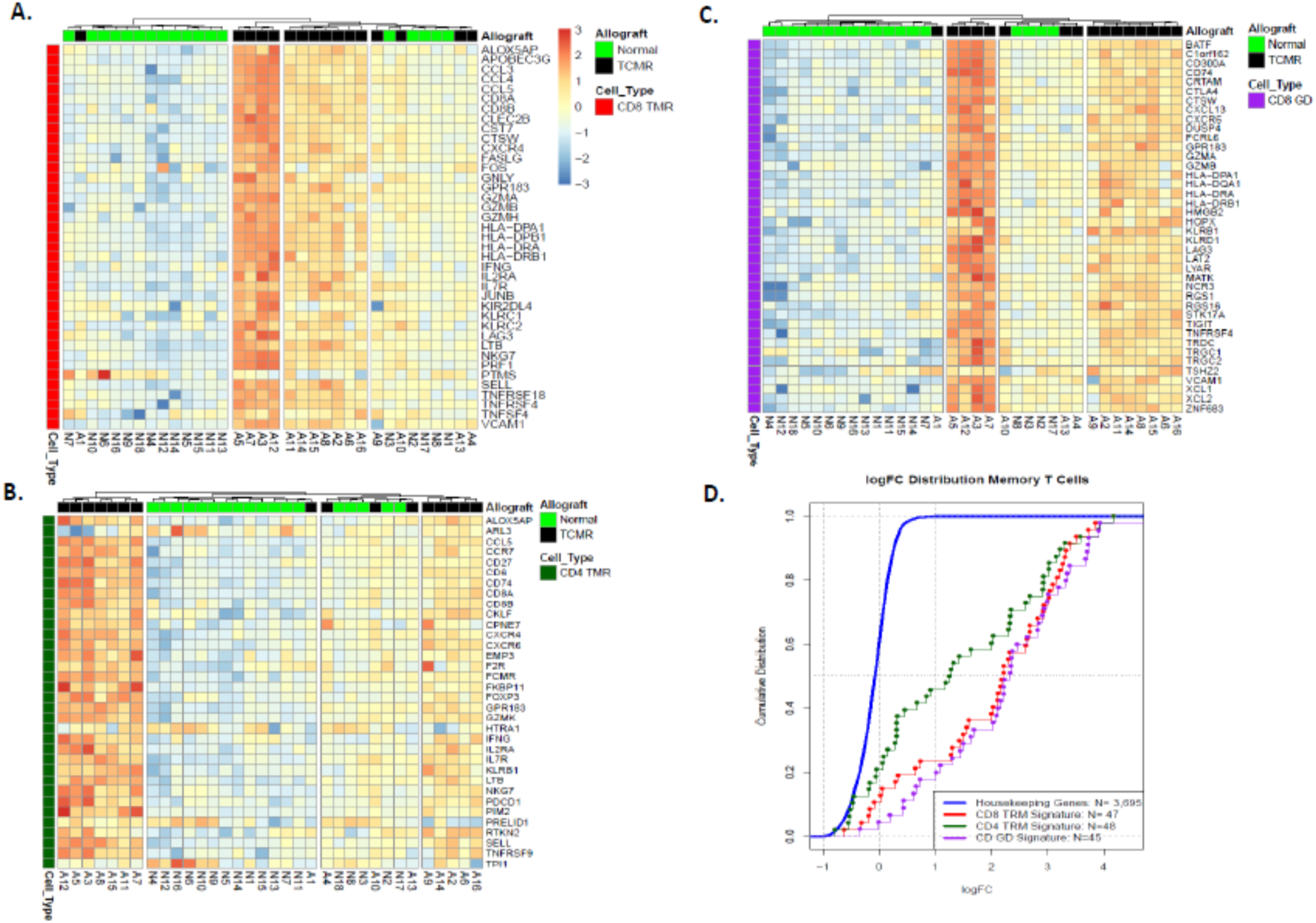
Intragraft abundance of mRNA Gene Set for CD8 Memory T-Cells, CD4 Memory T-Cells, and CD8 γδ T-Cells in TCMR Biopsies Compared to Normal Biopsies. **(A-C)** Heatmap visualization of hierarchical clustering analysis of intragraft mRNA abundance of CD8 memory T-cell gene set (A), CD4 memory T-cell gene set (B), and CD8 γ8 T-cells gene set (C). **(D)** CDF of the ratio of expression of 47 CD8 memory T-cells gene set (red line), 48 CD4 memory T-cells gene set (green line), and 45 CD8 γ8 T-cells gene set (pink line) in TCMR and Normal biopsies. Blue line represents the CDF of the ratio of 3,695 ubiquitously expressed housekeeping genes in TCMR and Normal biopsies. The right shift of the red, green, and pink lines compared to the blue line indicates the increased ratio of expression (TCMR versus Normal) of CD8 memory T-cells gene set, CD4 memory T-cells gene set, and CD8 γ8 T-cells gene set, respectively, compared to the ratio of expression (TCMR versus Normal) of housekeeping genes. This right shift was statistically significant by two-tailed Wilcoxon’s signed rank test.

Figure 5A-B, CDF plots, show heightened intragraft expression of mRNAs for Treg, Th1, Th2 and Th17 in TCMR biopsies compared to Normal biopsies. Furthermore, FGSEA of KEGG Treg, Th1, Th2 and Th17 cell differentiation pathways showed that these pathways were enriched in the up direction (P-FDR <0.0001) (Figure 5C-F).

**Figure 5.**
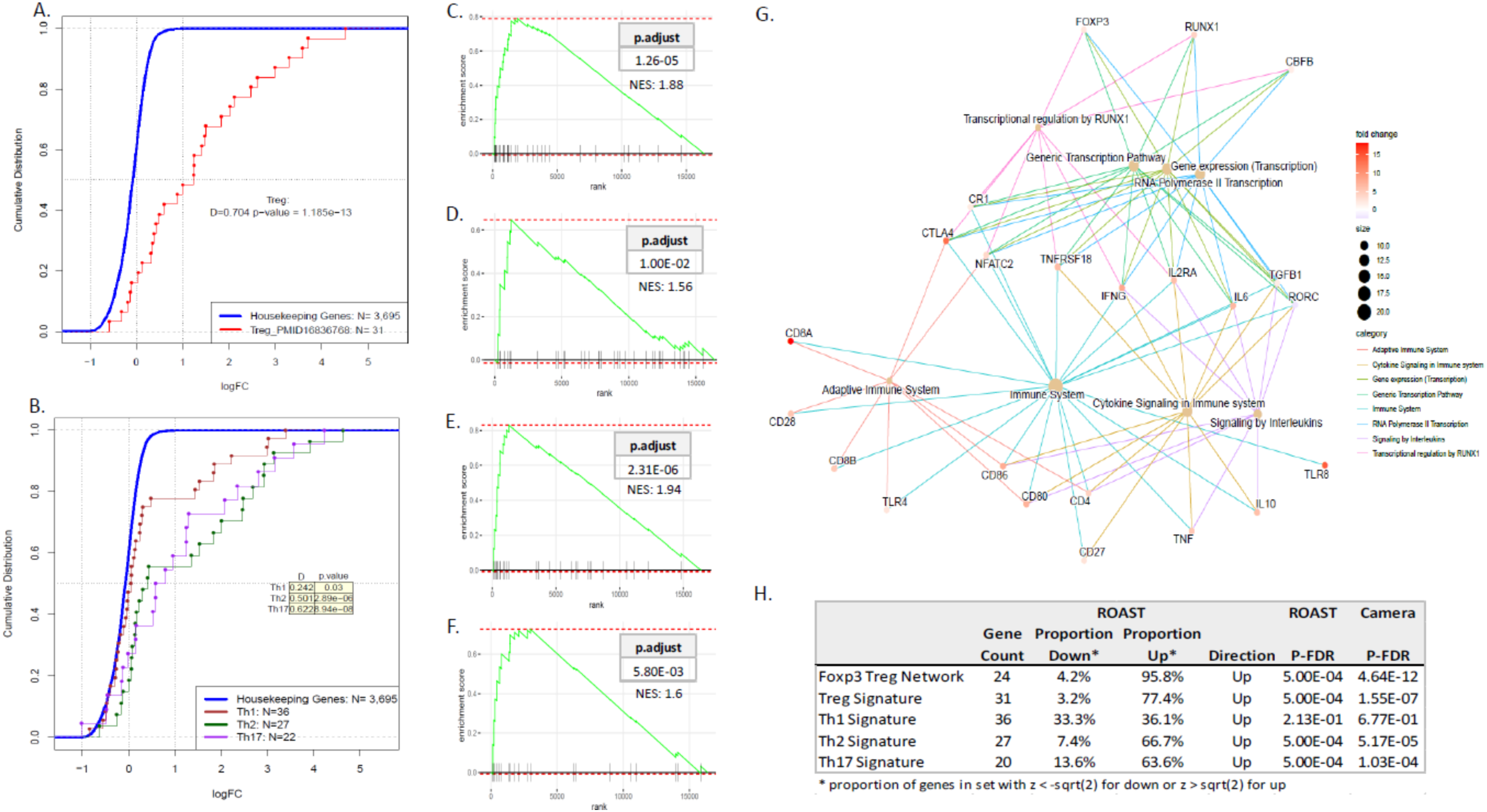
Treg, Th1, Th2 and Th17 mRNA Signatures and FOXP3 Gene Network: TCMR Biopsies vs. Normal Biopsies. **(A)** CDF of intragraft mRNA abundance of 31 Treg genes in TCMR. **(B)** CDF of intragraft mRNA abundance of 36 Th1 genes, 27 Th2 genes and 22 Th17 genes in TCMR. **(C-F)** FGSEA plots of Treg (C), Th1 (D), Th2 (E), and Th17 cells (F). **(G)** Reactome network interaction of FOXP3 genes based on intragraft abundance of 28 differentially expressed mRNAs. **(H)** Gene set testing using ROAST and CAMERA. Gene set testing is a differential expression analysis in which a P value is assigned to a set of genes as a unit. ROAST for gene-set testing uses rotation, a Monte Carlo method, for multivariate regression. CAMERA estimates and incorporates the variance inflation factor associated with inter-gene correlation.

RUNX1 stimulates transcription of IL2 and IFN-gamma (IFNG), and intragraft expression of these two genes is repressed upon binding of FOXP3 to RUNX1. The complex of FOXP3 and RUNX1, on the other hand, stimulates transcription of cell surface markers of Treg such as CD25, CTLA-4 and GITR (Figure 5G-H). We found that the FOXP3 gene network is increased in TCMR biopsies vs. Normal biopsies. There was also a positive association between TCR signaling pathway and Th1, Th2, and Th17 differentiation pathways (Supplemental Figure 3.).

### Expression of B-cell receptor signaling pathway genes

B cells can function as antigen presenting cells (APCs). We investigated whether B cell receptor signaling pathway genes expression is increased during an episode of TCMR. Of the 71 genes reported to participate in B cell activation and downstream signaling, 68 were measurable in the kidney allograft. Of the 41 genes that participate in the signal 1 for B cell activation, the abundance 23 mRNAs were higher and mRNAs for HRAS and RAC3 were lower in TCMR biopsies compared to Normal biopsies (Supplemental Table 4).

The B cell receptor complex (BCR) is made up of an immunoglobulin-alpha (CD79A) and immunoglobulin-beta protein (CD79B), and the abundance of mRNA for CD79A, CD79B and CD22 mRNA were higher in TCMR biopsies vs. Normal biopsies (Figure 6A). mRNAs for receptor related PTKs (BTK, SYK and LYN), and for the adaptor/linker molecules (BLNK and DAPP1) were higher in TCMR biopsies (Supplemental Table 4A). Of the 20 genes involved in the co-stimulation pathway, 7 were higher in TCMR biopsies (Supplemental Table 4B). CD19, which assembles with the antigen receptor of B lymphocytes to decrease the threshold for antigen receptor-dependent stimulation, was not detected in normal biopsies. Co-stimulatory molecules, CD18 and CR2 (CD21) were not differentially expressed between TCMR biopsies and Normal biopsies. There were 7 genes that have co-inhibitory effects on the B cell activation (Supplemental Table 4C). All 7 were expressed in the kidney; 6 of the 7 showed higher abundance in TCMR. Expression of 4 suppressors (checkpoints) of B cell function—CD72, CD22, FCGR2B and PTPN6—were increased in TCMR biopsies. Figure 6B is a heatmap of B cell signal 1 genes, co-stimulary genes and co-inhibitory genes. Figure 6C depict the KEGG B-B-Cell receptor enriched pathways. Our analysis revealed a molecular mechanism for the functionality of B cells during an episode of TCMR, and as found in T cells, mRNAs encoding activating proteins and mRNAs for inhibitory proteins were detected within the rejecting allograft.

**Figure 6.**
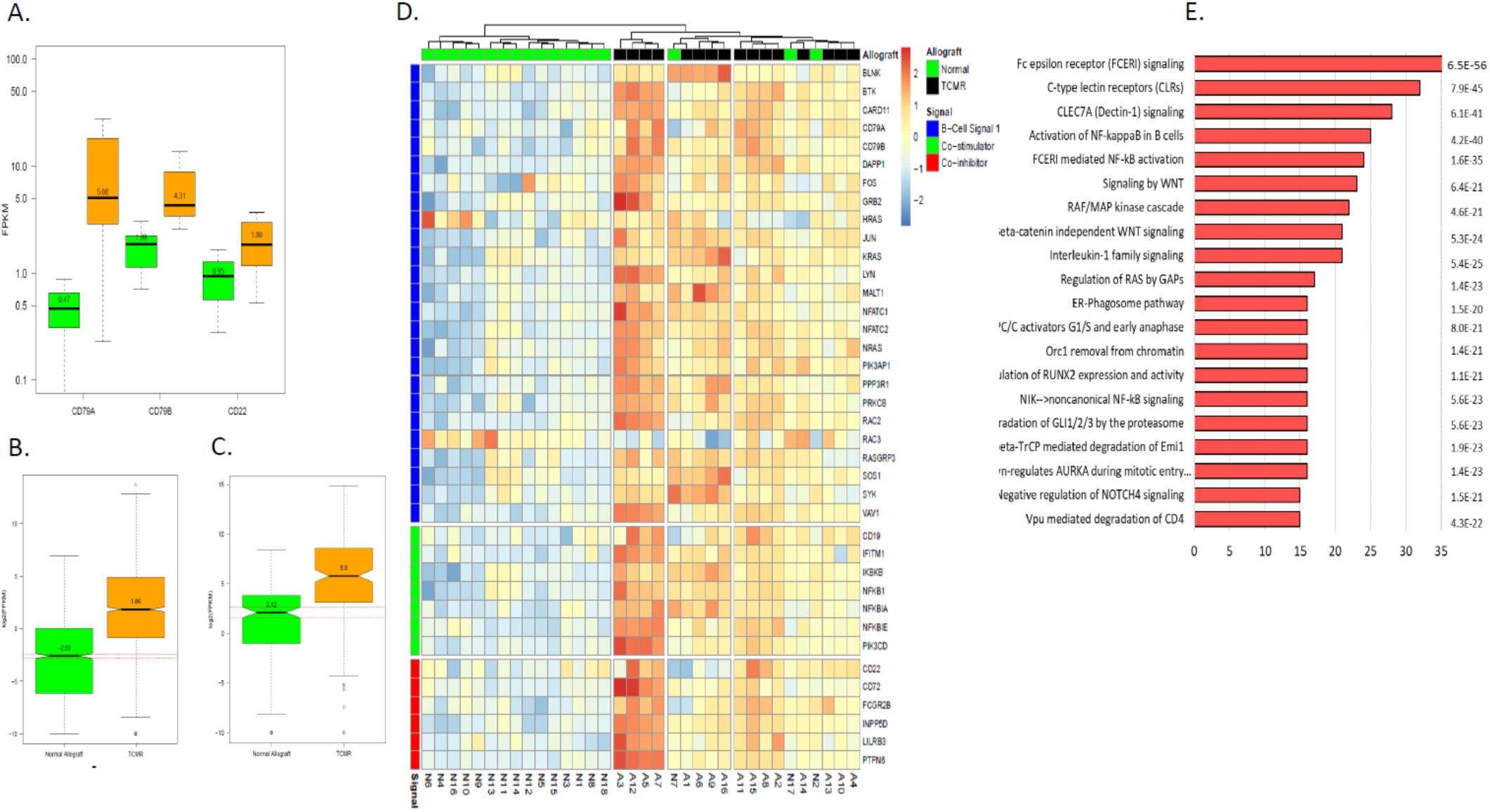
B-Cell Receptor Signaling is Increased in TCMR Biopsies Compared to Normal Biopsies. **(A)** Intragraft abundance of mRNAs encoding B-cell receptor complex genes in human kidney allograft biopsies (orange: TCMR [N=16]; green: Normal [N=18]). **(B)** Genes encoding variable chains of the immunoglobulin molecule. **(C)** Genes encoding constant chains of the immunoglobulin molecule. **(D)** Heatmap visualization of hierarchical clustering analysis of intragraft mRNA abundance of genes involved in B-cell receptor signaling. **(E)** Top Reactome pathways involved in B-cell receptor signaling

### Expression of leukocyte transendothelial migration genes

We queried intragraft expression patterns of mRNAs associated with leukocyte transendothelial migration (TEM). Of the 118 genes that participate in TEM, 100 were detected in the biopsies. Of these, 38 were leukocyte migration machinery genes and 23 of the 38 were differentially expressed with all but one showing higher abundance in TCMR biopsies vs. Normal biopsies (Supplemental Table 5A). Of the other 62 genes involved in facilitating transmigration, 24 were differentially expressed with increased abundance of 19 of 24 genes in TCMR biopsies (Supplemental Table 5B). A heat map of differentially expressed TEM KEGG pathways genes is illustrated in Figure 7A. Expression of levels of NADPH oxidase complexes NOX2 (CYBB) (FC 6.4, FDR 3.75E-16), CYBA (FC 1.2; FDR >0.5), NCF1 (FC 5.4, FDR 7.93E-11), NCF2 (FC 5.9, FDR 4.06E-14), NCF4 (FC 4.0, FDR 5.69E-12) are shown in Figure 7B-D. Key endothelial adhesion molecules ICAM-1 and VCAM-1, essential for leucocyte and endothelial cells physical interactions, showed higher abundance in TCMR biopsies. In addition, the abundance of mRNA for of matrix metallopeptidase 9 (MMP9), a protein involved in the breakdown the extracellular matrix, were significantly higher as well. Altogether, our findings provide a molecular basis for the interstitial infiltration of the kidney allograft by recipient’s leukocytes during an episode of TCMR.

**Figure 7.**
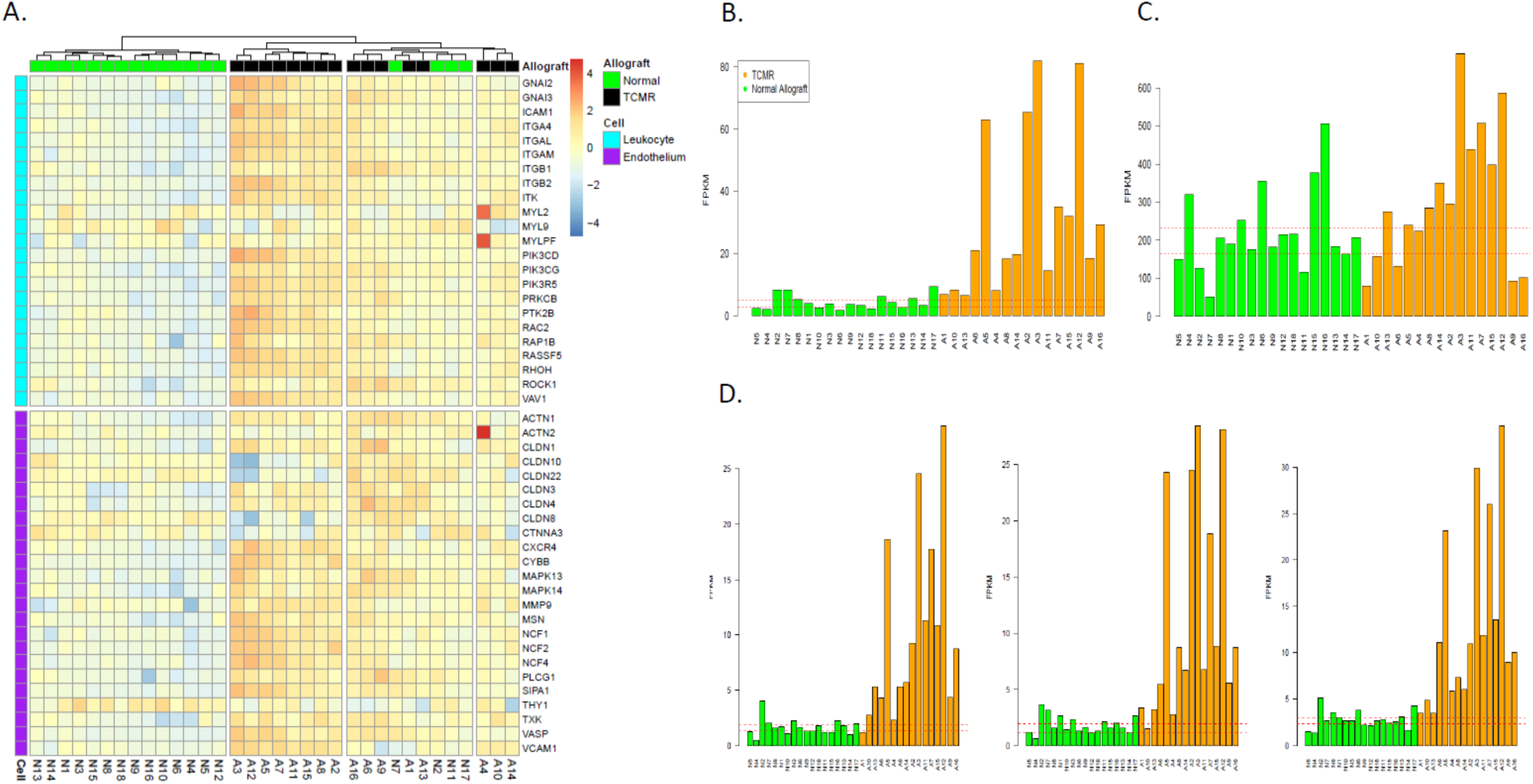
Leukocyte Transendothelial Migration Signature in the Kidney Allograft. **(A)** Leukocyte transendothelial migration KEGG pathway heatmap. **(B)** Expression of NOX2 (CYBB, P-FDR=3.8E-16), part of the reduced form nicotinamide adenine dinucleotide phosphate (NADPH) oxidase complexes in TCMR and Normal biopsies. Dotted redlines in represent the 95% CI expression of normal allograft samples. **(C)** Expression of CYBA(P-FDR>0.05) **(D)** Expression of NCF1 (P-FDR=7.9E-161) NCF2, (P-FDR=4.1E-14) and NCF4 (P-FDR=5.7E-12).

### Expression of glycolytic pathway genes

A kidney allograft undergoing acute rejection is comprised of two different cell types: kidney parenchymal cells and infiltrating immune cells. Interpretation of metabolic gene expression changes in a rejecting kidney— based on bulk RNA-seq data—is, therefore, challenging. We used the expression data of 15,716 genes in 4 normal kidney tissues and 4 normal human lymphoid tissues (4 each of bone marrow, lymph node, and spleen) from the human protein atlas (proteinatlas.org) and derived a ratio of these genes based on their expression in kidney to their expression in lymphoid tissue. A Kidney to Lymphoid tissue ratio (KLR) >1 indicates mRNA expression greater in kidney than in lymphoid tissue (Figure 8A). Of the 15,716 genes, 8314 (53%) were higher in lymphoid tissue (lymphocyte signature) and 7402 (47%) were higher in kidney tissue (kidney signature, Figure 8B). Of the lymphocyte signature genes, 88% were significantly increased in TCMR biopsies. Of the kidney allograft tissue signature genes, 86% showed decreased abundance in TCMR (Figure 8C, X^2^=3334.5, P<0.0001).

**Figure 8.**
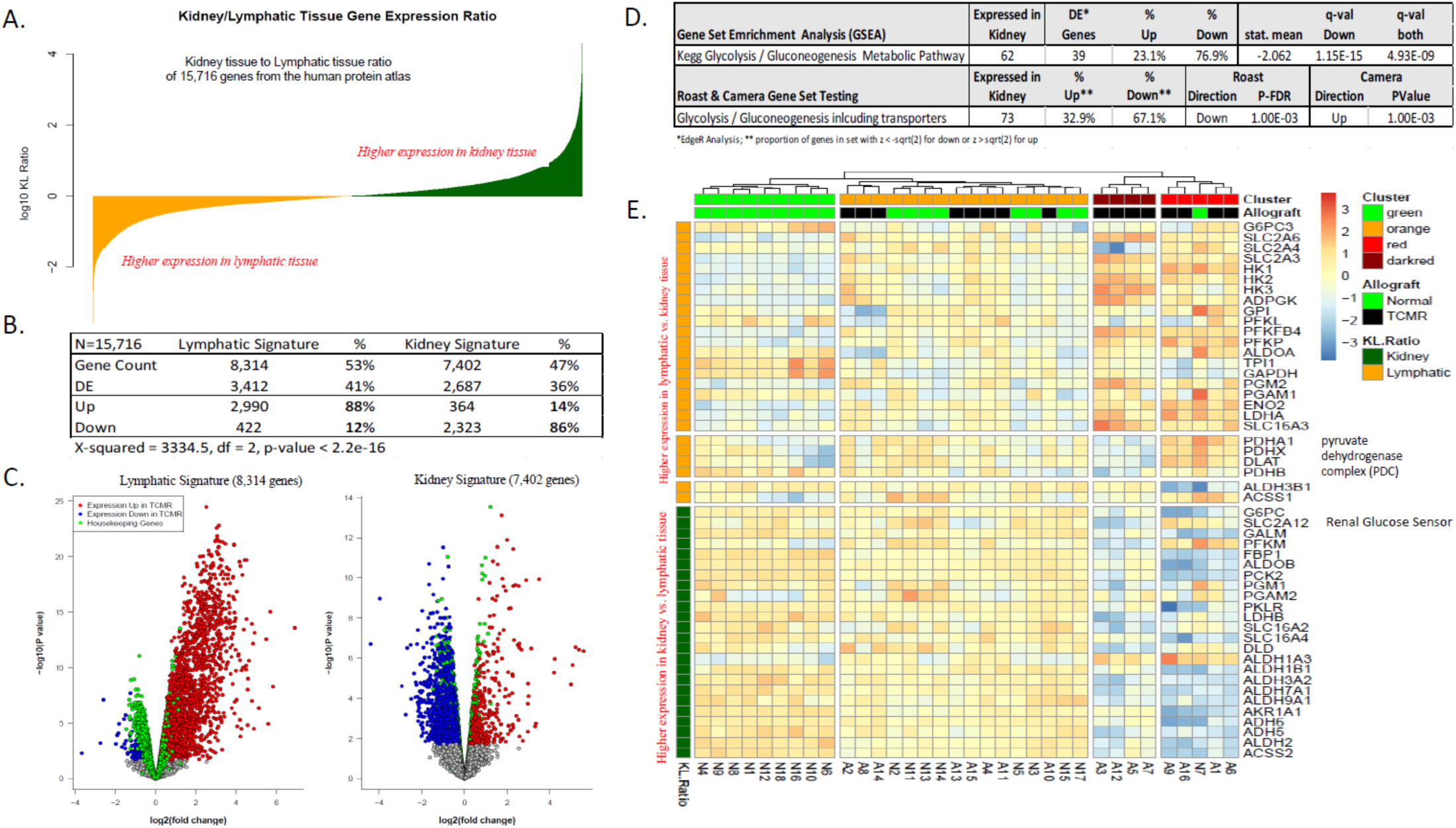
Altered Expression of Glycolytic Pathway Genes in TCMR. **(A)** Expression of 15,716 genes from the human protein atlas (proteinatlas.org). We derived a ratio of these genes based on their expression in normal human kidney tissue (N=4) to their expression in normal human lymphatic tissue (N=4 each of bone marrow, lymph node, and spleen) and represented it as log_10_ kidney tissue to lymphatic tissue ratio. **(B)** Of the 15,716 genes, 8314 (53%) were higher in lymphatic tissue and 7402 (47%) were higher in kidney tissue. **(C)** Volcano plot depicts the fold change in mRNA expression (TCMR vs. Normal) of these 8314 lymphatic tissue genes (left panel) and 7402 kidney tissue genes (right panel). **(D)** Gene set enrichment analysis of KEGG glycolysis and gluconeogenesis pathways in TCMR biopsies using GAGE. For a given pathway, the GAGE algorithm tests whether specific gene sets are significantly differentially expressed (each TCMR vs. all Normal) relative to the background whole gene set of 16,381 genes (each TCMR vs. all Normal) that we identified, using a rank-based two-sample T test. For each gene set, a global P value is derived on a meta-test on the negative log sum of all P values from the individual TCMR vs. Normal comparisons. The stat mean is the mean of the individual statistics from multiple gene set tests. Its absolute value measures the magnitude of gene-set level changes, and its sign indicates direction of the changes (positive: up regulated in TCMR, negative: down regulated in TCMR). The q value is the false discovery rate adjustment, by Benjamini-Hochberg method, on the global P value. Gene set testing using ROAST and CAMERA. **(E)** Heatmap visualization of hierarchical clustering analysis of intragraft mRNA abundance of 50 differentially expressed Glycolysis/Gluconeogenesis metabolic KEGG database pathway genes.

Gene set enrichment analysis of KEGG glycolysis and gluconeogenesis pathways showed 76.9% of genes were downregulated in kidney parenchymal cells (Figure 8D). Of a total of 73 KEGG glycolysis and gluconeogenesis pathway genes, 50 were differentially expressed between TCMR biopsies and Normal biopsies. Heatmap analysis of the 50 genes (Figure 8E) showed 4 different clusters. Cluster red and dark red in TCMR biopsies showed increased expression of glucose metabolism transporters, hexokinases, phosphor-fructokinase, and lactate dehydrogenase A. However, expression of pyruvate dehydrogenase complex (PDC) which converts pyruvate into acetyl-CoA was distinctly different between the two clusters—increased expression in cluster red and reduced expression in cluster dark red. Thus, in cluster dark red, there is likely aerobic glycolysis shifting pyruvate away from the TCA cycle. Furthermore, expression of SLC2A12, a renal glucose sensor, is low in cluster dark red and high in cluster red. Our analysis suggests that the T cells within the kidney allograft are metabolically active while in the kidney parenchymal tissue the metabolic activity is reduced.

### Expression of cell cycle progression and proliferation pathway genes

Prior to data analysis, we subdivided the large number of gene participating in cell cycle control and proliferation into functional units: cyclin-dependent kinases (CDKs) and cyclins required to activate the kinases, genes of the cyclin inhibitor complex which inhibit CDKs and thereby delay or stop the cell cycle progression, genes involved in the transcription and cell division machinery, and cell cycle damage control genes.

Of the 142 cell cycle genes, 117 were measued in the kidney allograft biopsies. Among the 117, 54 (46%) genes were differentially expressed with 47 of 54 increased in TCMR biopsies. Of the 17 cyclins and CDK genes expressed in the kidney, 10 were increased and 1 (CDK4) was decreased in TCMR (Supplemental Table 6A). Of 22 genes of the cell cycle inhibitor complex expressed in the kidney, 9 were increased and 1 decreased in TCMR biopsies. Several of these genes are part of the SCF complex (Skp, Cullin, F-box containing complex), a multi-protein E3 ubiquitin ligase complex, that controls G1/S and G2/M transitions (12). Of the 28 differentially expressed transcription and division machinery genes, 24 were increased in TCMR (Supplement Table 6B).

Cyclins (D1, D2 and D3) links cell environment with the core cell cycle machinery and are stimulated by growth factors and T-Cell activation pathways We found colony stimulating factor 1 ([CSF1], FC: 1.85, P-FDR 3.10E-08) and its receptor CSF1R (FC: 4.11, P-FDR 9.70E-13) to be associated with the D2 and D3 Cyclins (Supplemental Table 7A). IRAK4—essential for T cell immune response (FC: 1.54, P-FDR 6.94E-07), was associated with cyclin D2 and D3 together with several key genes of T cell activation pathways. Induced D cyclins bind and regulate CDK4 and CDK6 to activate the pocket-protein-E2F complex. Expression levels of late G1 E cyclins that activate various cell cycle proteins by phosphorylation were significantly increased and were associated with the E2F transcription factors. Furthermore, increases of cyclin A during the S phase and cyclin B—that drives the progression of many genes involved in DNA replication, centrosome, and chromosome function at the onset of mitosis, were significantly associated with E2F transcription factors (Supplemental Tables 7B). The two genes of the mini chromosome maintenance complex responsible for cell cycle DNA replication initiation and elongation were increased in abundance in TCMR and were significantly associated with cyclins E, A and B, as well as significantly expressed genes of the apoptosis pathway CASP 1, 3, 8, BCL2A1, BIRC3, BCL3 and BTK (Supplemental Tables 7C).

Four key damage control genes, ATM and ATR—the master controllers of cell cycle checkpoint signaling pathways that are required for cell response to DNA damage and for genome stability, and regulators TP53 (p53) and BRCA1, were significantly increased in TCMR. ATM, ATR, and TP53 are direct links of cell cycle genes to apoptosis. Furthermore, two apoptosis regulator CFLAR (FC 1.32, P-FDR 6.72E-03) and PAK2 (FC 1.46, P-FDR 1.81E-05) were associated with M phase genes ANAPC4 and CDC27, the cell cycle checkpoint ATM and the pocket gene RBL1 (Supplemental Table 7C). One of the key genes of the ER stress response with increased abundance, EIF2AK3 (FC 1.61, P-FDR 8.64E-06) was associated with cyclin D2 (r=0.89) and cell division cycle 27 (CDC27) (r=0.91) indicating a link between cell cycle progression and protein assembly.

Figure 9A shows the heatmap of the differentially expressed KEGG cell cycle genes. The pathway assessed with GAGE was significantly enriched in the up direction (P-FDR 3.10E-16). Figure 9B depicts the strong positive association between the cell cycle pathway and the p53 signaling pathway. The cell cycle gene expression profile and enrichment analysis identified here suggest a molecular basis for clonal expansion essential for TCMR, and its limitation by the p53 signaling and apoptosis.

**Figure 9.**
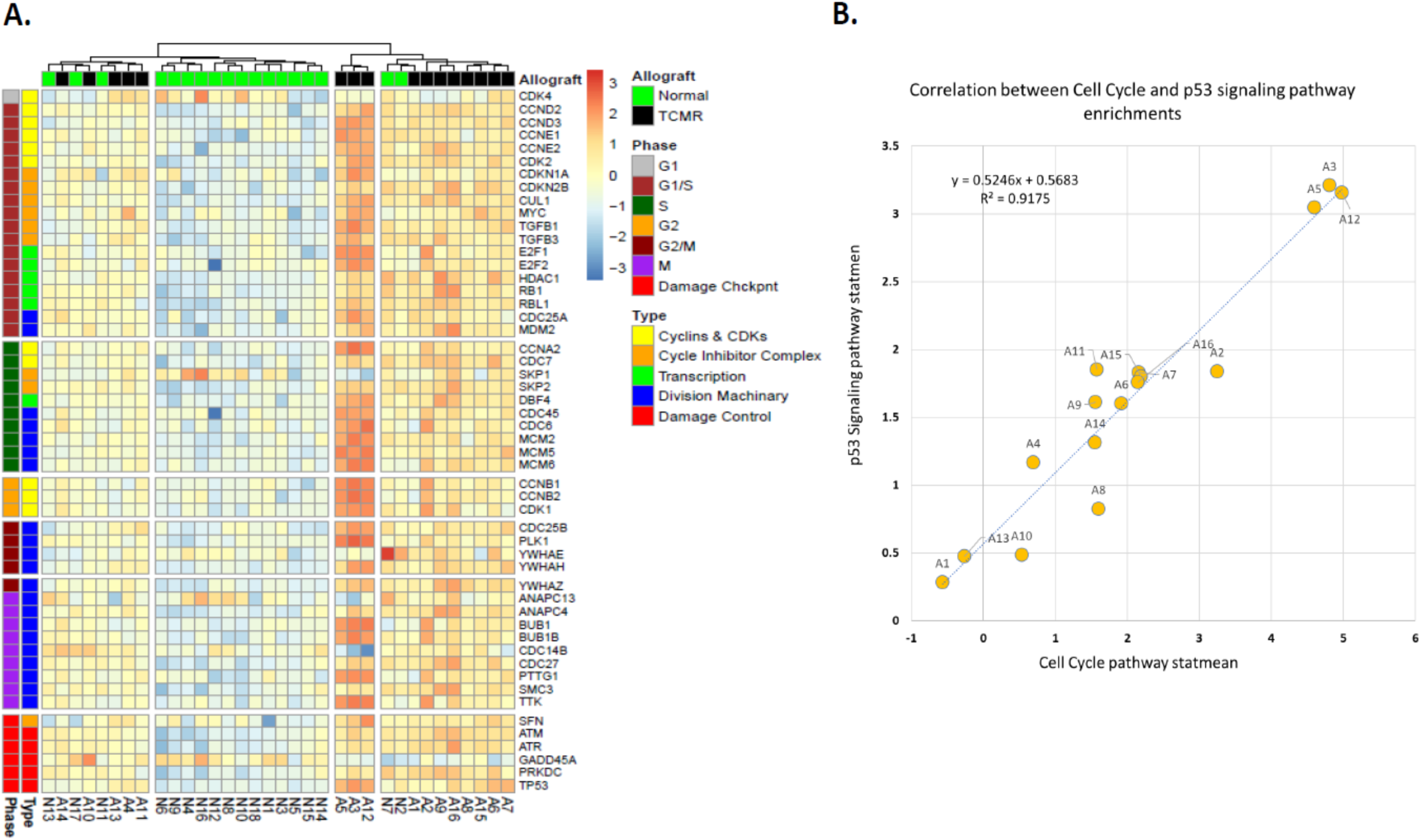
Intragraft Abundance of mRNAs for Cell Cycle Pathway Proteins are Higher in TCMR Biopsies Compared to Normal Biopsies. **(A)** Heatmap visualization of hierarchical clustering analysis of cell cycle pathway genes. **(B)** Positive association between p53 signaling gene-set and cell cycle pathway gene-set. Stat mean is the mean of the individual statistics from multiple gene set tests. Its absolute value measures the magnitude of gene-set level changes, and its sign indicates direction of the changes (positive: up regulated in TCMR, negative: down regulated in TCMR). The strength of the association is expressed as the coefficient.

### Expression of P53 pathway genes

Several stress signals, including DNA damage, oxidative stress, and activated oncogenes activate the P53 pathway in which the p53 protein acts as a transcriptional activator of p53-regulated genes. We identified 67 protein-coding genes implicated in P53 pathway—64 genes were expressed in the kidney allograft—and mRNA abundance was significantly higher for 32 genes and lower for 8 genes in TCMR biopsies (Supplemental Table 8 and Supplemental Figure 4). The key damage control genes, ATM and ATR, together with MDM2, MDM4, GORAB and CDKN2A were increased in TCMR. CDKN1A (p21) and SFN are two P53 target genes with increased abundance in TCMR that cause cell cycle arrest by inhibiting the action of cyclins. Inhibitory action of CDKN1A (p21) on cyclin D causes G1 arrest and the SFN (14-3-3sigma) gene and its product inhibits G2/M progression by cytoplasmatic sequestration of CDC2-cyclin B complexes. Expression of cyclin D2 and cyclin B2 were significantly higher in TCMR. In addition, SFN has been implicated in the transcriptional regulation of CDK-inhibitors as they modulate the transcription factors p53, FOXO and MIZ1 CDKN1A (p21) and DDB2, genes that participate in DNA nucleotide excision repair, are known to induce premature senescence. DDB2 mRNA abundance levels were significantly higher in TCMR. Also, there was a strong positive association between DDB2 mRNA and CDKN1A mRNA (r=0.82). Two additional P53 regulated genes showed higher mRNA abundance in TCMR; RRM2 catalyzes the formation of deoxyribonucleotides from ribonucleotides. Synthesis of the encoded protein (M2) is regulated in a cell-cycle dependent fashion and levels of GTSE1, which is only expressed in the S and G2 phases of the cell cycle. In response to DNA damage, the GTSE1 encoded protein accumulates in the nucleus and binds the tumor suppressor protein p53, shuttling it out of the nucleus and repressing its ability to induce apoptosis. The p53 signaling pathway was significantly enriched (P-FDR 6.35E-11). Our analysis shows an upregulated P53 pathway in TCMR and activation of the damage control and DNA repair machinery, as expected, in a high proliferative state.

### Expression of inhibitory receptor and T cell exhaustion pathway genes

Persistent T cell signaling can result in T cell exhaustion. Inhibitory receptor gene expression has been linked to T cell exhaustion and increased abundance of inhibitory receptor gene expression pattern is referred to as ‘exhaustion markers.’ In addition, increased abundance of genes for cell surface markers CD43/44, SPN, CD69, IL7R, and SELL has been associated with T cell exhaustion (13). Intragraft abundance of mRNAs for key T-cell inhibitory receptor genes linked to T-cell exhaustion (Figure 10A, Supplemental Table 9A) and key T-cell exhaustion markers (Figure 10B, Supplemental Table 9B) were higher TCMR biopsies compared to Normal biopsies. The abundance of the transcriptional repressor PRDM1 (BLIMP-1) which controls the terminal differentiation of cells was also significantly increased together with TBX21 (T-bet) and EOMES known to be co-expressed in CD8+ effector T cells. Our analysis suggest that T cell exhaustion is a molecular feature of TCMR.

**Figure 10.**
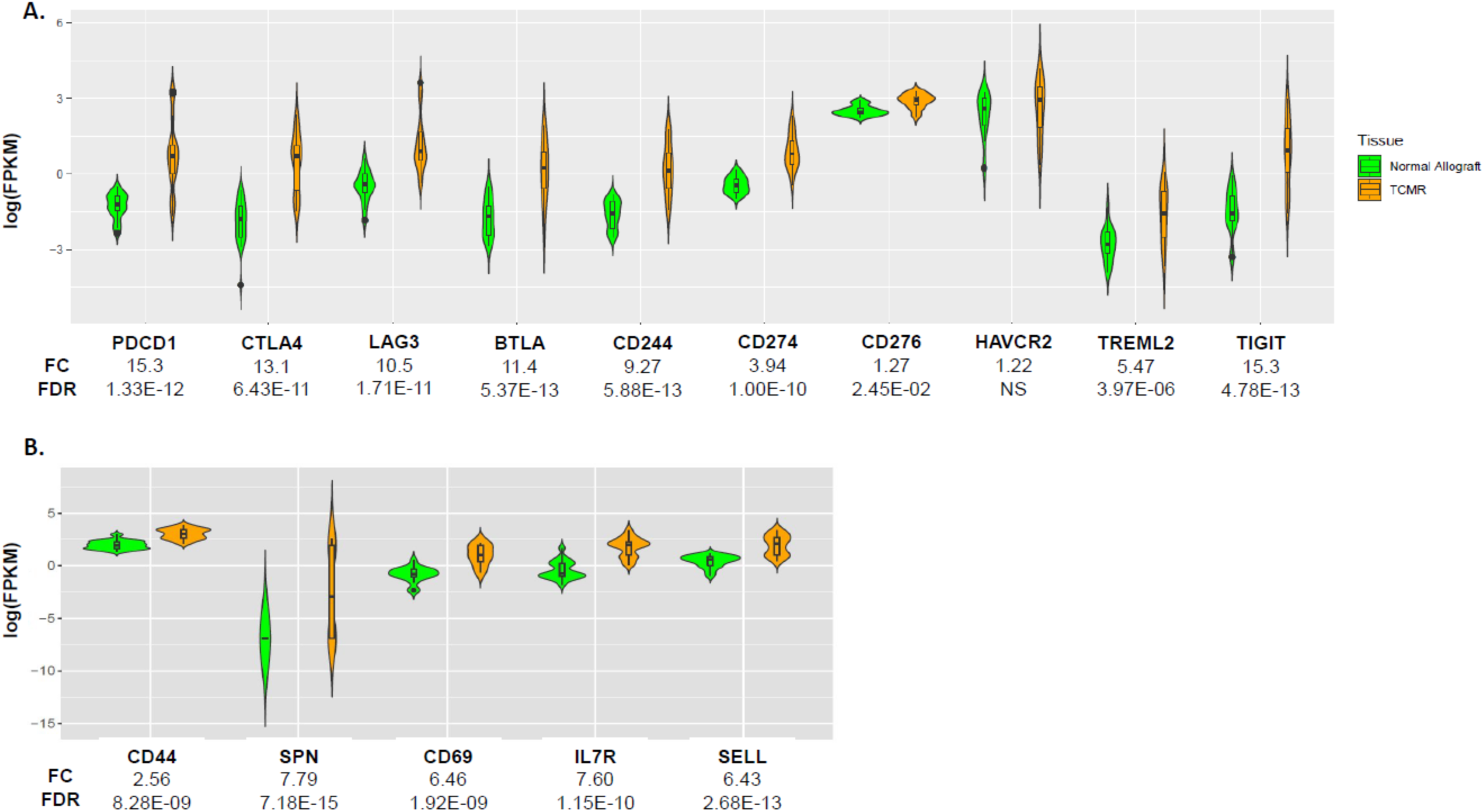
Intragraft Abundance of mRNAs Encoding T-Cell Inhibitory Receptors and Exhaustion Cell Markers in Kidney Allograft Biopsies. **(A)** Intragraft abundance of mRNA encoding T-cell inhibitory receptor proteins linked to T-cell exhaustion. The abundance of all but mRNA for HAVCR2 was higher in TCMR biopsies compared to Normal biopsies. **(B)** Intragraft abundance of mRNA encoding key T-cell exhaustion markers. The abundance of all mRNAs was higher in TCMR biopsies compared to Normal biopsies

### Expression of apoptosis and necroptosis pathway genes

Apoptosis is either due an intrinsic pathway—activated by non-receptor stimuli such as DNA damage or ER stress, or an extrinsic pathway that involves stimulation of members of the tumor necrosis factor receptor subfamily (TNFR). Of 140 genes identified to participate in apoptosis, 134 were expressed in the allograft and 72 of 134 were differentially expressed (Supplemental Table 10 and Supplemental Figure 5). None of the key pro-apoptotic genes of the intrinsic mitochondrial pathway such as BAK, BAX, BID were differentially expressed between TCMR and Normal biopsies, while significant reduction was observed in the expression levels of several intrinsic mitochondrial pathway genes such as the apoptosis initiator cytochrome C (CYCS), BCL2L1 which controls the production of reactive oxygen species and release of cytochrome C by mitochondria, AIFM1, essential for nuclear disassembly in apoptotic cells and ENDO6 which initiates replication of mitochondrial DNA. While members of TNFR subfamily, TNFSF10 and its receptors TNFRSF10A and TNFRSF10B, were significantly increased in TCMR, FADD—required for extrinsic pathway of apoptosis induced by TNFRSF10B was not differentially expressed between TCMR and Normal. Both extrinsic and intrinsic pathways converge to activate effector caspases 3 and 7, which initiate the caspase cascade. Caspase 3 (FC 1.9, P-FDR 2.310E-09) and caspase 7 (FC 1.3, P-FDR 1.6E-02) were significantly increased in TCMR biopsies. Caspase 3 can be directly activated by cytotoxic granules such as perforin and granzyme released by cytotoxic T cells and NK cells. Granzyme and perforin were increased in TCMR suggesting that apoptosis in TCMR is predominately accomplished through the granzyme B/perforin/caspase 3 pathway (Supplemental Figure 6). Several genes that are anti-apoptotic (pro-survival) by inhibiting expression of pro-apoptotic genes were differentially expressed (Supplemental Table 9) between TCMR and Normal. MCL1 is an inhibitor of the BCL-2 apoptotic activator; BCL2A1 which reduces the release of pro-apoptotic cytochrome c from mitochondria and blocks caspase activation was increased in TCMR as were the levels of BIRC3 and BIRC5 that block the action of the initiator caspase 9 and executive caspase 7. Cathepsins are part of the proteolytic machinery of apoptosis; expression of four cathepsins were increased in TCMR. There was a significant upregulation of the apoptosis pathway (P-FDR 2.97E-15). Homeostasis of cellular organisms requires a balance between cell proliferation and apoptosis. We further show the relation between cell cycle pathway and apoptosis pathway genes, by calculating a cell cycle and apoptosis score based on the KEGG pathway profiles. Both, the cell cycle, and apoptosis pathways are tightly associated but shifted towards apoptosis in TCMR (Figure 11A-C).

**Figure 11.**
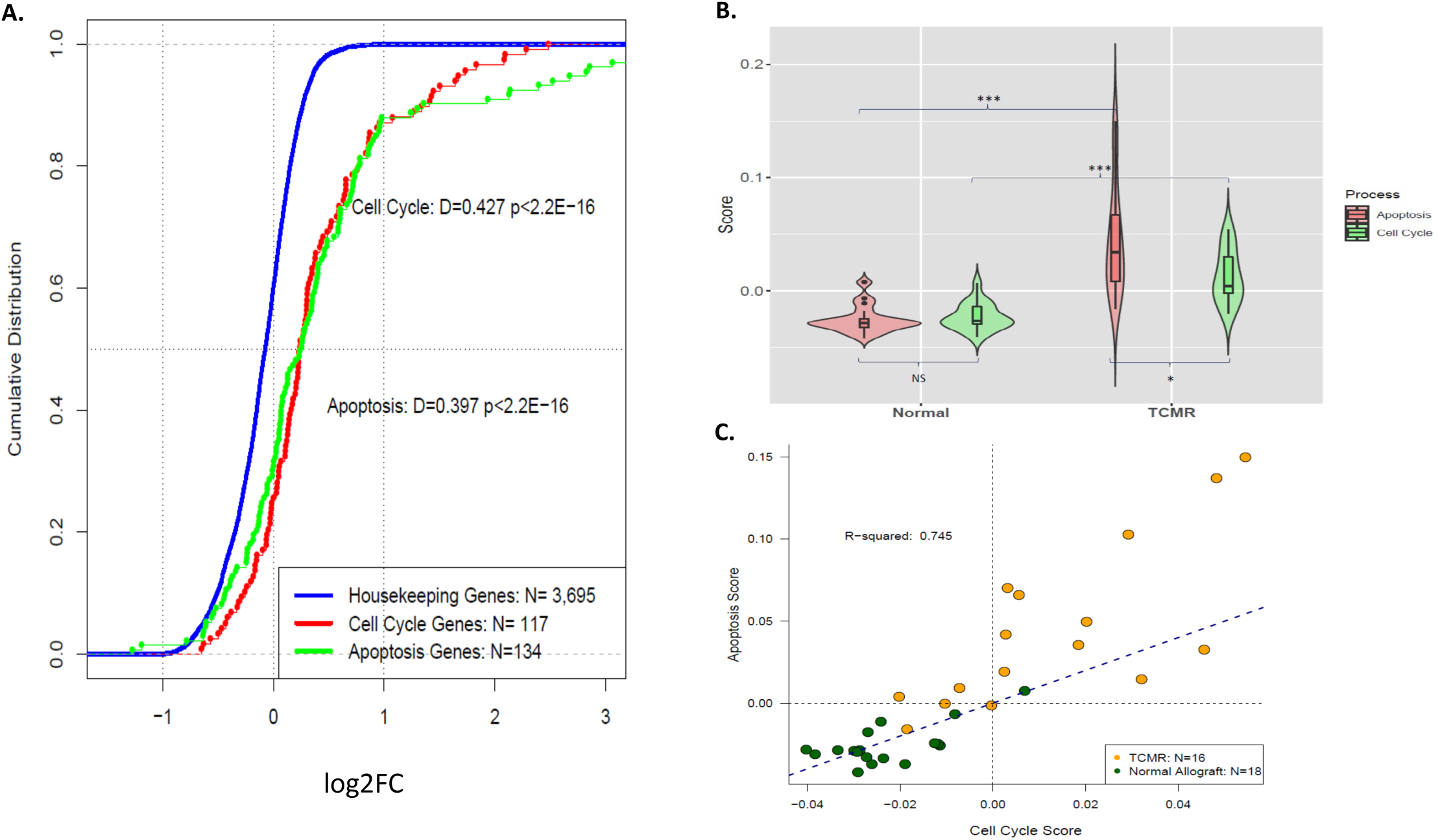
Relationship Between Cell Cycle Genes and Apoptosis Genes in Kidney Allograft Biopsies. **(A)** CDF of cell cycle genes and apoptosis genes. Blue line represents the CDF of the ratio of 3,695 ubiquitously expressed housekeeping genes in TCMR and Normal biopsies. Red line represents the CDF of the ratio of 117 cell cycle genes in TCMR and Normal biopsies. Green line represents the CDF of the ratio of 134 apoptosis genes in TCMR and Normal biopsies. **(B)** Violin plots of cell cycle gene and apoptosis gene scores in kidney allograft biopsies. Scores were calculated by using the ‘AddModuleScore’ function of Seurat (satijalab.org/seurat/) to determine the average expression levels for each gene set at sample level, subtracted by the aggregated expression of control gene sets. All analyzed genes (N=16,381) were binned based on averaged expression and the control genes are randomly selected from each bin. Log_e_-normalized gene counts for each sample were divided by the total counts for that sample and multiplied by the scale factor. Scores of apoptosis genes and cell cycle genes were significantly higher in TCMR biopsies compared to Normal biopsies. **(C)** Positive association between cell cycle genes and apoptosis genes. The line of identity (blue dotted line) represents homeostatic equilibrium where cell cycle and apoptosis are equal.

We also identified 129 genes that participate in necroptosis pathway, with 127 expressed in the kidney of which 77 were differentially expressed with expression levels significantly increased in TCMR for 57 genes (Supplemental Table 11 and Supplemental Figure 7A). Expression levels of receptor interacting protein kinase1 and 3 (RIPK1 and RIPK3), key regulators of necroptosis, were significantly increased, together with MLKL which plays a critical role in tumor necrosis factor (TNF)-induced necroptosis. Expression levels of three key downstream targets of RIPK3, CYBB which generates superoxide in phagocytes, GLUL which catalyzes the synthesis of glutamine from glutamate, PYGL important for glycogenolysis and MLKL were significantly increased. The KEGG necroptosis pathway was significantly enriched in the up direction (P-FDR 8.02E-15) and was significantly associated with the cytosolic DNA sensing pathway activating innate immune system pattern recognition receptors (Supplemental Figure 7B). Taken together, our analysis reveals increased apoptosis and necroptosis machinery in TCMR biopsies.

### Expression of extracellular matrix remodeling during an episode of TCMR

In view of cell injury and cell death during an episode of TCMR, we assessed gene expression pattern of matrix related genes in the biopsies. The matrix genes were detected using the matrisome database (14) comprised of 734 genes categorized into collagens, glycoproteins, proteoglycans, extracellular matrix (ECM) affiliated, regulators, and secreted genes (Supplemental Table 12). Analysis of the whole gene set showed 42% of the 734 genes to be increased in TCMR biopsies (P-FDR 5.00E-04). Activation and degradation were independent of time from transplantation to biopsy (Supplemental Figure 8). Matrisome proteins renew via proteolysis and synthesis. The homeostasis of connective tissue therefore depends on rates of ECM production, rate of ECM removal, and the mechanical properties of the ECM (15). We hypothesized that under normal condition, the ECM gene expression signature will be like those of housekeeping genes but with cellular stress during acute rejection, the expression ECM genes will shift. Figure 12A-B shows the CDF plots of extracellular matrix organization and degradation genes, respectively, during TCMR compared to the housekeeping genes (P<2.2E-16) supporting increased ECM organization and degradation during TCMR. We created a collagen score for both collagen formation and biosynthesis pathway and collagen degradation pathway, based on log-normalized gene counts for each sample using the ‘AddModuleScore’ function in Seurat (16). Figure 12C shows significant increases in collagen formation and degradation scores during TCMR compared to Normal biopsies. Within TCMR or Normal biopsies, however, the collagen degradation score was higher than the collagen formation score, but this difference was not statistically different. Figure 12D shows the positive association between the collagen formation score and the degradation score with the line of identity showing a shift towards degradation in TCMR.

**Figure 12.**
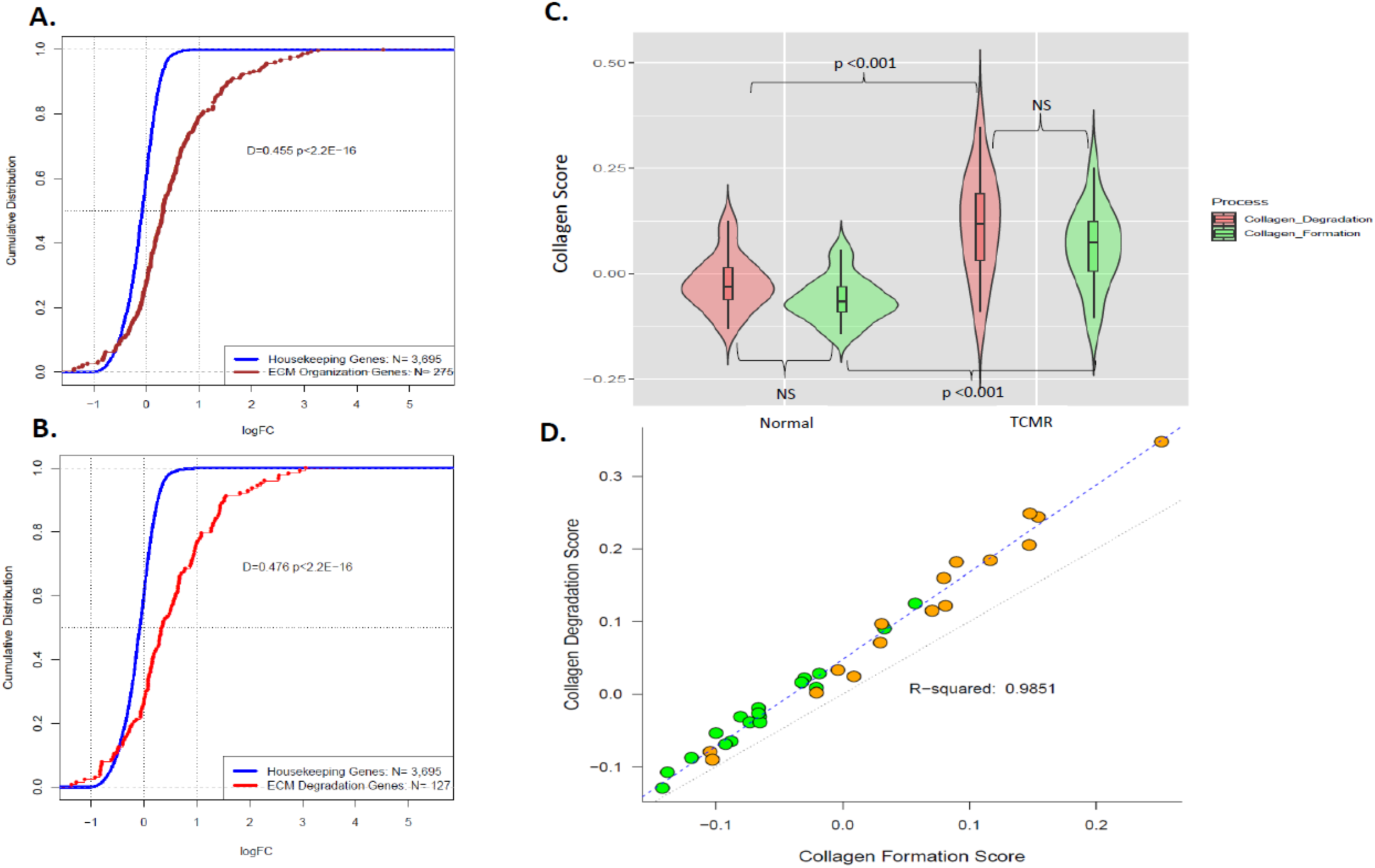
Genes Encoding Extracellular Matrix Regenerative Process are Increased in TCMR Biopsies. **(A)** CDF of the ratio of expression of 275 ECM Organization Reactome pathway genes (brown line) in TCMR and Normal. Blue line represents the CDF of the ratio of 3,695 ubiquitously expressed housekeeping genes in TCMR and Normal biopsies. **(B)** CDF of the ratio of expression of 127 ECM Degradation Reactome pathway genes (red line) in TCMR. Blue line represents the CDF of the ratio of 3,695 ubiquitously expressed housekeeping genes in TCMR and Normal biopsies. **(C)** Violin plots of collagen scores in TCMR biopsies and Normal biopsies. Scores were calculated by using the ‘AddModuleScore’ function of Seurat (satijalab.org/seurat/) to determine the average expression levels for each gene set at sample level, subtracted by the aggregated expression of control gene sets. All analyzed genes (N=16,381) were binned based on averaged expression and the control genes are randomly selected from each bin. Reactome collagen formation and biosynthesis pathway genes are shown in green. Reactome collagen degradation pathway genes are shown in red. Log_e_-normalized gene counts for each sample were divided by the total counts for that sample and multiplied by the scale factor. Collagen formation and degradation gene set scores were significantly higher in TCMR biopsies compared to Normal biopsies. The was no significant difference in the scores between collagen formation and degradation gene sets in either type of biopsies. **(D)** Association between collagen formation genes and collagen degradation genes. The line of identity (black dotted line) represents homeostatic equilibrium where collagen formation and collagen degradation are equal. Normal Allograft green, TCMR orange dots.

We expanded the matrisome matrix by including 43 additional genes and created a fibrosis gene expression matrix of 777 genes. We used this matrix to perform a weighted correlation network analysis to correlate mRNA gene expression with clinical phenotype information yielding a 242 gene expression matrix highly correlated to acute rejection Banff Scores and validated by STRING protein-protein interaction database (17). Figure 13A, immune system and ECM pathway interaction network, is based on the significantly expressed genes of the 242 gene matrix shows a protein-protein interaction network grouping the ECM differentially expressed genes into 6 network clusters: extracellular matrix (B), immune system (C) with chemokine signaling (D), glycosylation (E), platelet pathway related genes (F) and 18 highly connected nodes (genes) at the center (Figure 13A and Supplemental Table 13). The network illustrates how the immune system (C) and chemokine activation (D) relates to ECM regeneration (B) by increased ECM degradation and ECM formation while platelet degranulation promotes vasodilation and increased blood vessel permeability (F). Figure 13B shows the heatmap of this network by relative gene expression values. Our analysis of the impact of acute rejection on the ECM reveals strong turnover of ECM characterized by an increased matrix degradation and matrix formation expression signature. In addition, the ECM turnover appears to be blunted in allografts with preexisting fibrosis.

**Figure 13.**
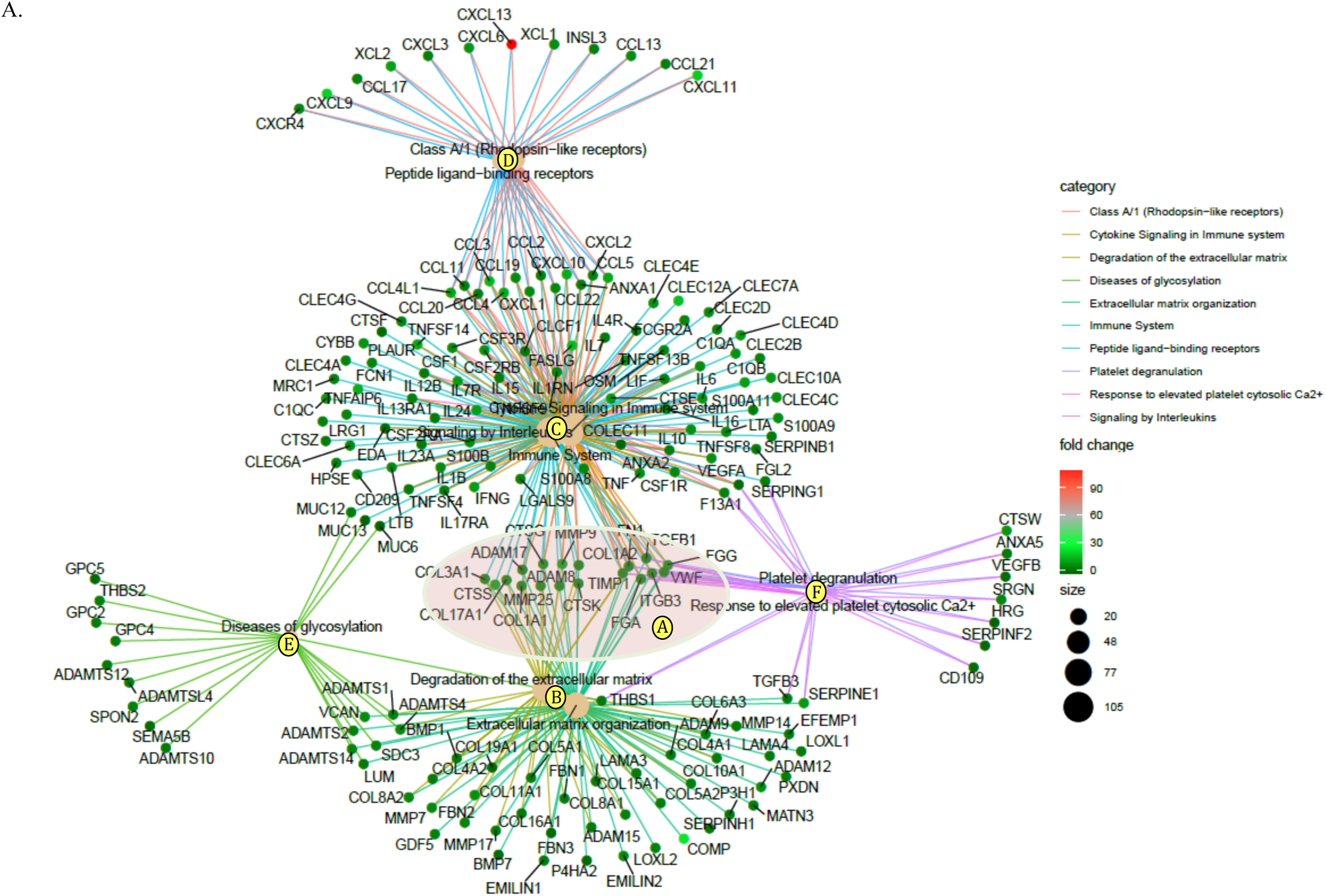
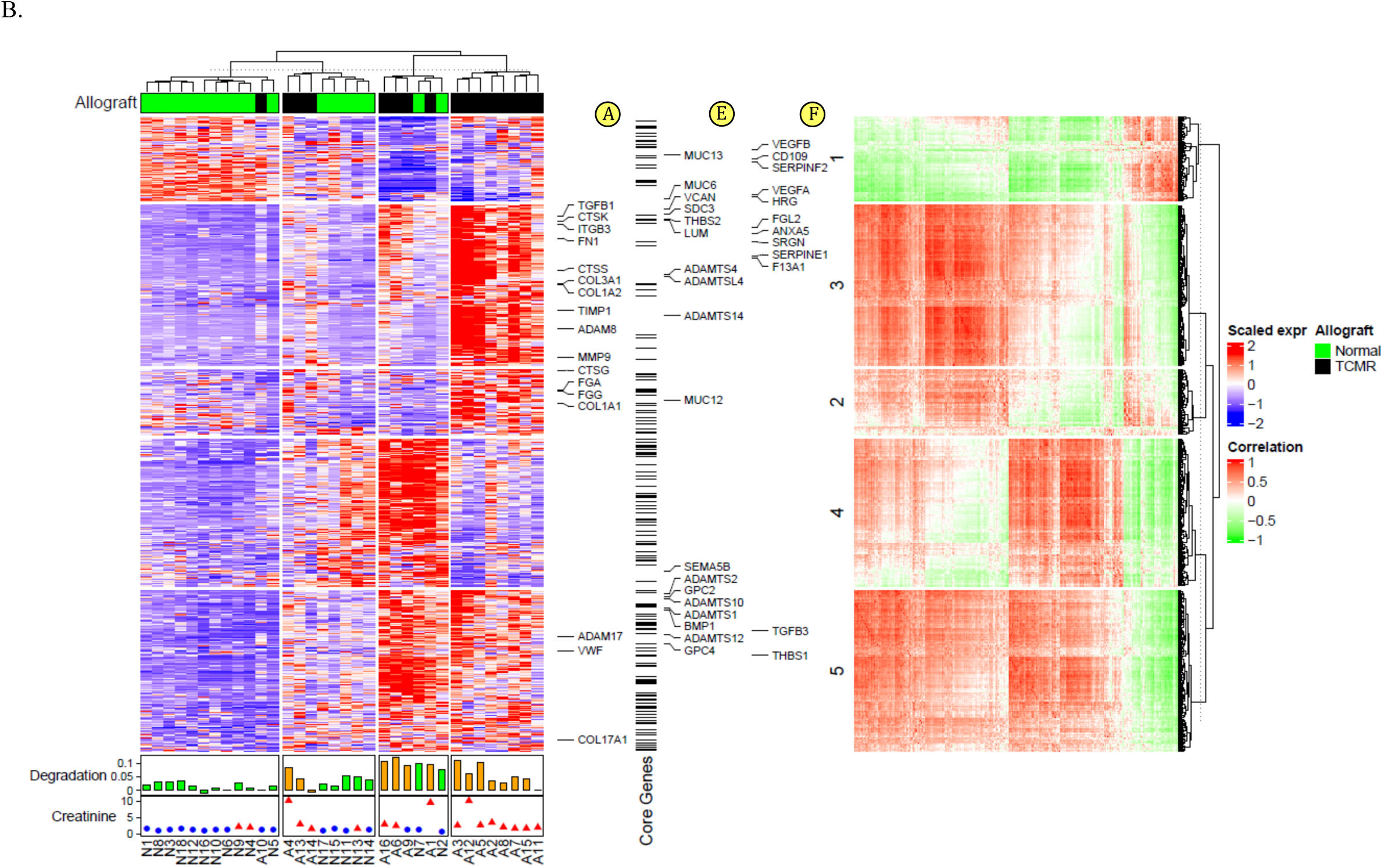
Immune System and Extracellular Matrix Pathway Interaction Network in TCMR. **(A)** Top 10 out of 23 enriched Reactome pathways based on 242 differentially expressed genes matching the profibrotic gene expression matrix, and with confirmed protein-protein interactions according to the STRING protein-protein network interaction database. The gene-concept network graph depicts a linkage of genes as a network to extract the complex association in which a gene may belong to multiple annotation categories. Genes are shown as dots; the color of the dot represent the Fold Change value. The statistical significance of the pathways is an over-representation (enrichment) analysis that tests whether a specific pathway is over-represented among the differentially expressed genes—more genes of a particular pathway than would be expected by chance. All ten Reactome pathways were statistically significant (P-FDR<0.05). **(B)** Heatmap of this network by relative gene expression values (left panel). Biopsy samples (columns) clustered into four clusters and gene expression (rows) into five clusters. The bottom of the heatmap shows shift towards degradation as sample bar plots (Degradation Score – Formation Score) and relation to serum creatinine levels at the time of biopsy. Genes of network nodes A, E, and F from Figure 12A are shown in relation to the five gene expression clusters (middle panel). Correlation plot (right panel) shows the five gene expression clusters now clustered by gene correlations > 0.5 to more than 20 other genes.

To assess the impact of TCMR on ECM changes in allografts with or without IFTA, we stratified the biopsy samples by Banff score for IFTA (Supplemental Figure 9). Only 1.26% or 204 genes were differentially expressed when comparing TCMR with IFTA and TCMR without IFTA. The IFTA subgroup analysis did show decreased matrix formation and degradation pattern in TCMR with IFTA suggesting reduced injury repair and ECM remodeling in TCMR with IFTA.

## Discussion

We identified genome-wide transcriptional changes in human kidney allografts by RNA sequencing prototypic human kidney allograft biopsies and deciphered adaptive immune molecular circuits operative during an episode of TCMR. The key gene-based pathways we found enriched in TCMR biopsies include antigen processing and presentation, T cell and B cell signaling, costimulation, second messenger generation and phosphorylation, metabolism, cell cycle progression and proliferation, apoptosis and necroptosis, leukocyte transendothelial migration, and extracellular remodeling. Intriguingly, we identified that the immune repertoire was not monotonic and counter regulatory molecular signatures were evident in TCMR biopsies-enrichment of proinflammatory cytokine and cytotoxic effector pathways genes countered by enrichment of anti-inflammatory and T cell inhibitory receptor gene signatures; cell cycle progression and proliferation gene sets offset by apoptosis and necroptosis pathway genes. The immune perturbations were associated with gene signatures of heighted T cell metabolism and T cell exhaustion. Our findings confirm and extend in very important ways TCMR-associated transcripts previously identified using microarrays and based in part on mouse kidney transplants and cultured cells (8-10, 18, 19).

Antigen processing and presentation to T cells and B cells occur within specialized lymphoid organs. The abundance of mRNA for HLA class I and class II antigens and mRNA for several of the minor histocompatibility antigens were all higher in TCMR biopsies compared to Normal biopsies. Also, mRNA for IFNG and TNFA, two potent inducers of HLA expression, were over expressed in TCMR biopsies.

Our finding that the antigen processing and presentation machinery resides within the allograft raises the intriguing possibility that the allograft itself could function as a novel lymphoid structure. Direct as well as indirect allorecognition could occur within the allograft since the rejecting allograft is infiltrated not only by recipient’s T cells and B cells but also by antigen presenting cells.

Allorecognition is contingent upon recipient’s T cells and B cells, the primary effector cells of the adaptive immune system. Surprisingly, we found not only enrichment of T cell receptor signaling pathway genes but also enrichment of B cell receptor signaling pathway genes in prototypic TCMR biopsies compared to Normal biopsies. Plenary T cell activation requires antigenic signaling via the TCR/CD3 complex and costimulation via the CD28 and related molecules. We document in TCMR biopsies higher abundance of mRNAs encoding proteins for T cell transmembrane and downstream signaling and the machinery for the phosphorylation of CD3 zeta chain. Our cataloguing of genes and pathways offers also novel targets including PTKs to regulate allorecognition and to prevent and/or treat TCMR.

Cells with cytolytic activity have been eluted from rejected experimental and clinical allografts and the granzymes and perforin are the main components of the granule exocytosis cytolytic machinery. We demonstrate in this study that all four species of granzymes- A, B, M and K, and perforin are expressed at a higher abundance in TCMR biopsies compared to normal biopsies. Our leveraging of single cell RNA sequencing data (11) identified that CD8^+^ memory T cells, CD4^+^ memory T cells, and CD8^+^ γδ T cells are all enriched in TCMR biopsies. Since memory T cells and CD8^+^ γδ T cells are relatively resistant to immunosuppressive drugs, our new findings suggest a mechanistic basis for almost 40% of TCMR episodes in humans being refractory to anti-rejection therapy.

T cell subsets with opposing effects were identified within the rejecting allograft using subset specific gene expression signatures. We observed not only higher abundance of mRNA for proinflammatory Th1, Th2 and Th17 T cell subsets in TCMR biopsies but also mRNA for the anti-inflammatory FOXP3 Treg subset. The T h1, Th2 and Th17 cell subsets, via production of inflammatory cytokines and by promoting lytic activity of CTL and NK cells, are likely to be significant mediators of TCMR. On the other hand, FOXP3+ via its binding of RUNX transcription factors and inhibiting IL2 and IFNG secretion and induction negative regulatory molecules such as CTLA4 is likely to counter the effect of proinflammatory T cell subsets. The fine balance between the proinflammatory and anti-inflammatory T cell subsets may determine TCMR reversal and ultimate graft survival, as observed in our earlier urinary cell mRNA profiling studies (20, 21).

Our study demonstrates that B cells are among the graft infiltrating cells in biopsies classified as TCMR. The abundance of mRNA for BCR and mRNA for costimulatory proteins such as CD40 were higher in TCMR compared to Normal biopsies. BCR downstream signaling pathway gene set analysis identified higher abundance of both canonical and noncanonical NFKB gene sets. As found with T cells, not only mRNAs for costimulatory but also mRNAs for coinhibitory pathways were enriched in TCMR biopsies.

Interstitial infiltration and tubulitis are the histologic hallmarks of TCMR. An intriguing question is the processes that aid transendothelial migration of T cells from their vascular niche into the interstitial space via paracellular and transcellular pathways (22). It has been reported that ICAM1-mediated signals activate an endothelial cell calcium flux required for leukocyte migration (23) and VCAM1 signals activate endothelial cell NADPH oxidase involved in the opening of an endothelial passage through which leukocytes can extravasate (24). Thus, endothelial cells serve not only as a scaffold for lymphocyte adhesion but also play an active role in promoting lymphocyte migration (25). Our finding of heightened expression of mRNAs for leucocyte adhesion and migration machinery genes such as ICAM1 and endothelial genes such as CYBB, VCAM1 and NADPH oxidases complex in TCMR provide a mechanistic basis for the histologic features of TCMR, and importantly suggest target molecules to prevent TCMR. Four-dimensional imaging of T cells in kidney rejection in mouse models has shown that intragraft graft DCs extend their cellular processes into capillary lumina and contact blood-borne T cells to initiate transmigration of T cells as well as arrest T cells within the allograft interstitial space (26). The high abundance of intragraft DCs in TCMR biopsies (7, 27) () represents another mechanism for the graft reactive cells to emigrate from their vascular niche into the interstitial space.

Clonal expansion is characterized by cell growth, a high cell division rate and a highly active metabolic state. Activated cells expand through successive rounds of cell divisions from a limited number of naïve T cells (about 10^5^-10^6^) to up to 10^4^ fold (28). The T cell response follows a triphasic pathway of activation, proliferation and differentiation before becoming functionally and phenotypically “exhausted” as shown in the settings of chronic infection, autoimmunity and in cancer (29). Importantly, the extent and magnitude of clonal expansion in vivo is directly related to the amount of antigenic stimulation provided to the naive T cell during primary activation (28). We document for the first time in humans, how different amounts of antigen stimulation, as shown in gene expression heatmaps of MHC I and MHC II KEGG and Autosomal Minor HC Antigens Pathways (Figure S1and S2), impact the various degrees of gene expression of T cell and B activation pathway heatmaps (Figure 3 and Figure 8) at the individual sample level. Animal studies have shown that a key molecular process of T cell proliferation is the entry of cells into the cell cycle, a complex process tightly controlled by the ordered expression of cyclins, the activation of cyclin-dependent kinase enzymatic activity, and the subsequent phosphorylation of relevant substrates (30). Our analysis of TCMR in humans of the cell cycle pathway underscores a gene expression pattern of high proliferation with increased activation of close to 60% of all cyclin and cyclin-dependent kinases, in particular D Cyclins regarded as links between the cell environment and the core cell cycle machinery (31). These D cyclins are stimulated by growth factors and T-Cell activation pathways. Colony stimulating factor 1 and its receptor highly correlated with the D2 and D3 Cyclins. The abundance of MYC oncogene, known to promote early T cell growth and glycolytic and glutaminolytic metabolism (32) was significantly increased. In addition, abundance was increased in more than 60% of all differentially expressed transcription and division machinery genes and less than 50% of the cell cycle inhibitor complex genes were differentially expressed, as such, p27 (CDKN1B) a known inhibitor of D Cyclins, was not differentially expressed. Altogether, the cell cycle gene expression profile and enrichment analysis suggest a significant upregulation of cell cycle activity as required by high proliferating cells. Furthermore, we found in TCMR biopsies a close to 19 times fold change of the transcription factor IRF4, that was found in animal studies to act as a dose dependent regulator of the metabolic reprogramming of activated T cells, thereby linking metabolic function with the clonal selection and effector differentiation of T cells (33). Investigating the different adaptive immune system pathways, we found that the abundance of many genes that characterize T cell effector function significantly elevated, among them cytotoxic granules (Granzyme A, B, H, K, M and Perforin, major effectors of cytotoxic T cells functions, in which fold change increases ranged between 7 and 10 times.

Proliferation, development, and sustaining T cell effector functions requires a significant uptake of cellular nutrients (32). While the metabolic demand for naïve T cells is relatively low and oxidative phosphorylation is sufficient to cover their energy obligation (34) many in vitro and animal studies document that the main energy source of activated T cells is glucose, catabolized through the glycolytic cascade to pyruvate (35). Pyruvate can either be reduced to lactate to regenerate NAD+ in active T cells or is converted by pyruvate dehydrogenase to acetyl-CoA to nourish the tricarboxylic acid (TCA) cycle for further oxidation in naïve and memory T cells. Although we show highly increased abundance of regulators of metabolic reprogramming in activated T cells and a high cell cycle pathway enrichment consistent with proliferation, interpretation of metabolic gene expression changes in a rejecting kidney—based on bulk RNA-seq data—is challenging. The kidney microenvironment of allografts undergoing acute rejection is comprised of two fundamentally different cell types: kidney parenchymal cells and infiltrating immune cells with different metabolic demand and not unlike as described in cancer, when metabolites are restricted by exacerbated consumptions, both, proliferating lymphocytes and renal parenchymal cells, compete for the same nutrients. Renal cells may struggle to sustain their metabolism and function, clinically manifested by rapidly deteriorating renal function in acute rejection. To better understand the metabolic changes that occur during TCMR and to segregate kidney parenchymal metabolic changes from the metabolic demands of graft infiltrating cells, we sorted the KEGG Glycolysis/ Gluconeogenesis pathway genes by a Kidney to Lymphatic Tissue Ratio (KL ratio) derived from gene expression data of the Protein Atlas. Our analysis revealed that in TCMR biopsies, the Lymphocyte ratio signature showed markedly increased gene expressions in glucose metabolism transporters, hexokinases, phosphor-fructokinase (PFKFB4) and Lactate dehydrogenase A (LDHA). LDHA converts pyruvate to lactate, characteristic of anaerobic glycolysis with a shift to lactate to provide nutrition for the expanding biomass of activated immune cells. However, Kidney ratio signature genes in TCMR biopsies showed reduced gene expression indicating low metabolic activity of the kidney parenchyma.

Our analysis further shows how in adaptive immune system T cell activation phosphatidylinositol 3 kinase (PI3K) and phospholipase C gamma 1 (PLCG1) together with DAG propagate the signal to three major pathways: PI3K–AKT, RAS-RAF-ERK and the NF-kB signaling and activate three nuclear transcription factors gene sets: NF-kappa B, NFAT and AP-1(FOS, JUN) responsible for cell proliferation, survival and apoptosis. Downstream targets of these pathways differentially expressed in TCMR participate in cell cycle progression, cell survival and apoptosis.

Following clearance of antigen, expanded effector T cells massively contract and ∼90% die and are eliminated mainly via apoptosis (36). Homeostasis of cellular organisms requires a balance between cell proliferation and apoptosis. Animal and in vitro studies from cancer research suggest that the connection of proliferation and apoptosis is ether positive (37, 38), the more cells proliferate the more apoptosis occurs or, alternatively, an increased level of demand can be met by reduced apoptosis (39). Our human data in TCMR documents that many of the genes that directly link the cell cycle to apoptosis were significantly increased suggesting a strong interplay between the two pathways. Furthermore, key damage control genes, ATM and ATR, master controllers of cell cycle checkpoints and required for cell response to DNA damage and for genome stability and regulators p53 (TP53) and BRCA1, were significantly increased and link cell cycle genes to apoptosis and we show that the degree of cell cycle activation is correlated with the p53 regulator pathway activation. We show, by calculating a cell cycle and apoptosis score based on the KEGG pathway profiles, how both, the cell cycle and apoptosis pathways are tightly associated together and is shifted towards apoptosis in TCMR. Apoptosis that eliminates damaged or redundant cells by activation of caspases can be mapped by three often interrelated pathways: the extrinsic, intrinsic and a third pathway that activates caspase-3 by cytotoxic granules (e.g., Perforin and Granzyme B) directly released form cytotoxic T-cells and natural killer cells (NK). While some genes of the extrinsic and intrinsic apoptosis pathways were differentially expressed, key genes such as TNFRSF1A, TRADD and FADD (extrinsic) and none of the key pro-apoptotic genes of the intrinsic mitochondrial pathway such as BAK, BAX, BID were differentially expressed between TCMR biopsies and Normal biopsies while expression levels of apoptosis initiator cytochrome C (CYCS), BCL2L1 which controls the production of reactive oxygen species and release of cytochrome C by mitochondria, AIFM1, essential for nuclear disassembly in apoptotic cells as well ENDO6 which initiates replication of mitochondrial DNA were significantly reduced. However, Caspases 3, 7, 8, and 10 were all significantly increased together with GZMB (Granzyme B) and PRF1 (Perforin) suggestive that apoptosis during an episode of TCMR is predominately activated through the Granzyme B pathway and that the Granzyme B/ Perforin are the major apoptosis pathway in TCMR.

Studies in mice investigating the adaptive T cell response during infection under conditions of prolonged antigen exposure, discovered that T cells follow a distinct pathway of differentiation and become functionally and phenotypically “exhausted” (40). In exhausted cells, levels of expressions are not only increased in one inhibitory receptor, such as PD-1 (PDCD1), but multiple inhibitory receptors induced during T-Cell activation are persistently elevated which leads to a profound inability of T cells to respond to activation signals and as described first in chronic infection a progressive dysfunction worsening over time. The importance of T cell exhaustion in organ transplantation is twofold: (i) early after the discovery of T cell exhaustion it was postulated that T cell exhaustion might contribute to long-term graft survival (41) and could be leveraged to improve long-term graft outcome (42), (ii) although reactivating the exhausted T cells in cancer as a strategy of cancer treatment has been highly successful, using this intervention in transplant patients suffering from cancer was associated with an increased risk of rejection (43). A recent study in serially collected peripheral blood mononuclear cells in 26 kidney transplant recipients using mass-cytometry identified a distinct subset of circulating exhausted T cells and their relationship to allograft function (44). We report here for the first time in human kidney allografts in acute rejection, the expression profile of five cell markers described as being activated in T cell exhaustion and significant abundance increases in in nine of 10 inhibitory receptors associated with exhaustion. Furthermore, the transcription factor BLIMP-1 that mediates a transcriptional program in various innate and adaptive immune tissue-resident lymphocyte T cell types such as tissue-resident memory T (Trm), NK and natural killer T (NKT) cells was also significantly increased together with TBX21 (T-bet) and EOMES known to be co-expressed in CD8+ effector T cells (45). Overall, the gene expression signatures of inhibitory receptors and T cell exhaustion indicate a pattern of a high proliferative state as observed in rapid clonal expansion together with exhaustion and activation of tissue resident memory cells.

We investigated the impact of TCMR on the gene expression pattern of the extracellular matrix (ECM).. We observed an increase in matrisome genes in TCMR. This increase was not related to time from transplantation to biopsy. Furthermore, we found a shift towards ECM degradation in TCMR, a tissue regenerative process. Often tissue regenerative phase is in homeostasis and has no long-term sequala. However, during tissue repair, excess accumulation of connective tissue results in the replacement of the normal parenchyma. Several cell types are critical to repair damage, restore tissue integrity, normal function, and control tissue inflammation. Fibroblasts associate with macrophages and lymphocytes at sites of injury and respond directly to immune mediated signals. Macrophages integrate signals from the tissue microenvironment, the innate and adaptive immune responses and their associated fibroblasts promote, inhibit or reverse fibrosis. We have previously shown an increased abundance of key inflammatory M1 and tissue repair M2 macrophages gene markers and a significant shift of the respective distribution functions of the xCell gene sets for these cell types (7). While macrophages show a marked shift, the fibroblast gene signature was not increased in TCMR. Furthermore, the expression of α-smooth muscle actin (ACTA2) involved in the transition of fibroblasts to myofibroblasts required for fibrotic tissue generation was significantly decreased in TCMR suggesting that while tissue restoration and repair is increased in TCMR, the homeostatic balance is maintained with no increase in fibrosis. However, there was a blunted ECM regeneration in TCMR allografts with preexisting fibrosis.

Our study has limitations. We demonstrate differences at the mRNA level between TCMR and normal and not at the protein or functional level. Not every differentially abundant mRNA needs to be different at the protein level. However, the coordinated expression of multiple genes in multiple pathways suggests mRNA expression patterns are informative of underlying biology. Moreover, several molecular circuits we resolved have been demonstrated in elegant animal models. Also, we provide a cross sectional view of the transcriptome; hence, our findings should be interpreted as associative rather than causative. Our intragraft gene expression patterns were characterized using ensemble RNA sequencing and single cell RNA sequencing may yield more granular insights.

In summary, utilizing RNA sequencing to investigate human kidney allografts, we developed a comprehensive landscape of adaptive immune system gene expression patterns and strength during an episode of TCMR of human kidney allografts. We identified heterogeneity in gene expression within the normal and TCMR biopsies. Our data provides an unbiased view of the intragraft molecular circuits during TCMR and highlights the key events that includes increased antigen processing and presentation, T cell and B cell signaling, costimulation, second messenger generation and phosphorylation, metabolism, cell cycle progression and proliferation, apoptosis and necroptosis, leukocyte transendothelial migration, and extracellular remodeling. The deciphered immune repertoire was not monotonic and counter regulation was evident in TCMR. Our data suggest the hypothesis that molecular categorization of TCMR is needed for transplant precision therapeutics and prioritization of therapeutic targets to prevent and/or treat TCMR.

## Methods

### Study group and kidney allograft biopsies

We performed RNA-Seq of 34 kidney allograft biopsies from 34 adult recipients of kidney allografts, a subset of kidney allograft recipients transplanted and followed at our center. The biopsies were retrieved from our Biobank since they represented prototypic examples of biopsies classified as Banff TCMR or Banff Normal/ Nonspecific categories. The percutaneous needle core biopsies were done under ultrasound guidance and were read and reported independently by two renal pathologists at our center (S.S and S.V.S), based on the Banff 2017 update of the Banff ‘97 classification of allograft pathology (46, 47) and blinded to gene expression data. Biopsy tissue sections were stained with hematoxylin and eosin, periodic acid–Schiff, and Masson trichrome, as well as for polyomavirus, and for complement factor 4 degradation product d (C4d). Each patient provided a single biopsy sample for this study.

### RNA sequencing of kidney allograft biopsies

At the time of allograft biopsy, a portion of the biopsy tissue was immediately immersed in RNAlater^®^ RNA stabilization solution (Life Technologies, Grand Island, NY) and stored in −80°C for subsequent batch processing. From the stored sample, we isolated total RNA using the miRNeasy mini kit (Qiagen, Inc., Valencia, CA). We used NanoDrop 1000 spectrophotometer (ThermoScientific, Wilmington, DE) to measure the quantity and purity of the RNA and Agilent 2100 Bioanalyzer (Agilent Technologies Inc., Santa Clara, CA) to measure the integrity of RNA. We used 400 ng total RNA from each biopsy sample for mRNA sequencing. We used the TruSeq™ sample preparation kit v2 (Illumina, Inc., San Diego, CA) to prepare individual cDNA libraries. Briefly, this consists of poly-A selection of mRNAs and conversion to single-stranded cDNA using random hexamer primer followed by second strand generation to create double stranded cDNA. Sequencing adapters were then ligated to the fragmented cDNA. This was followed by PCR amplification and pooling of the libraries. The biopsy samples were sequenced in two different batches (batch 1 and batch 2) as part of a larger RNA sequencing studies in our laboratory. The first batch contained of 15 samples (10 TCMR and 5 Normal) were sequenced single-end, 6 pooled libraries per lane of a flow cell, on a HiSeq 2500 (Illumina, Inc., San Diego, CA) sequencer. The second batch contained 19 samples (6 TCMR and 13 Normal) and were sequenced paired-end, 10 pooled libraries per lane of a flow cell, on a HiSeq 4000 sequencer. The raw sequencing data were stored in FASTQ format.

### Sequencing data analysis

Data processing of all sample used the same analysis pipeline. Quality control of FASTQ data was performed using FastQC:Read QC (48) and adapter sequences, if any, removed with trimmomatic (49). The 101bp single and paired-end reads were then aligned to human genome (GRCh37/hg19 assembly) using Tophat2 (50). The human UCSC reference annotation was used to align and quantitate the transcripts. Overall mapping rate was 91.8%±3.9% (mean±SD). Transcript assembly and transcript abundance was determined using Cufflinks (51). The hg19 genes file was used as reference annotation guide to determine the FPKM for each transcript with Cufflinks and raw counts with HTseq (52). We further established that normalization by library size, TMM (weighted trimmed mean) and log2 transformation of raw counts, as described in the edgeR user guide, and sufficient to remove the potential batch effects. Differential RNA-seq analysis was performed with edgeR (53). After removal of duplicate transcripts and keeping genes with at least 1 cpm in one sample, the initial transcript size of 22,582 reduced to 16,381. We used R packages GAGE (54), limma (55), and pathview (56) for gene set or pathway analysis, pheatmap for creating heatmaps, WGCNA package (57) for weighted co-expression analysis using, and Reactome pathway enrichment analysis by Reactome (58). We used ROAST (59) and CAMERA (60) for gene set testing. For assessing gene set enrichment, we used FGSEA (fast gene set enrichment analysis) (61). We used the *matrisome* database to assess the genes encoding the ‘matrisome’—an ensemble of extracellular matrix and matrix-associated proteins (14), and created a profibrotic gene expression matrix by adding 43 additional genes based on literature review. We used this matrix to construct a gene co-expression network for the 34 biopsy samples (62).

### Statistical analysis

We used edgeR to determine the fold changes; edgeR fits a negative binomial model of the sequence data and derives probability values for differential expression using an exact test (63). Probability values were adjusted for FDR; P-FDR < 0.05 was considered statistically significant.

## Study approval

Kidney transplant recipients reported herein provided written informed consent to participate in the study and the informed consent was obtained prior to their inclusion in the study. Our Institutional Review Board approved the study. The clinical and research activities that we report here are consistent with the principles of the “World Medical Association Declaration of Helsinki on Ethical Principles for Medical Research Involving Human Subjects” (64), and the “Declaration of Istanbul on Organ Trafficking and Transplant Tourism” (65).

## Author Contributions

Designed research: M.S., T.M.

Conducting experiments: H.Y., C.L., C.S., T.M.,

Acquiring data: F.B.M., H.Y., C.L., C.S., D.M.D., J.Z.X., V.K.S., T.M.

Analyzing data: F.B.M., H.Y., J.Z.X., S.S., S.V.S., V.K.S., O.E., T.M.

Providing reagents: J.Z.X., M.S.,

Writing the manuscript: F.B.M., M.S., T.M.

## Acknowledgements

Supported in part by awards from the National Institutes of Health to M. Suthanthiran (NIH MERIT Award, R37-AI051652), T. Muthukumar (K08-DK087824), and Weill Cornell Medical College (Clinical and Translational Science Center Award UL1TR000457).

## Notes

**Conflict of Interest Statement** The authors have declared that no conflict of interest exists.

### Competing Interest Statement

The authors have declared no competing interest.

## References

1. Hariharan S, Johnson CP, Bresnahan BA, Taranto SE, McIntosh MJ, and Stablein D. Improved graft survival after renal transplantation in the United States, 1988 to 1996. N Engl J Med. 2000;342(9):605–12.

2. Cole EH, Johnston O, Rose CL, and Gill JS. Impact of acute rejection and new-onset diabetes on long-term transplant graft and patient survival. Clin J Am Soc Nephrol. 2008;3(3):814–21.

3. Alkadi MM, Kim J, Aull MJ, Schwartz JE, Lee JR, Watkins A, et al. Kidney allograft failure in the steroid-free immunosuppression era: A matched case-control study. Clin Transplant. 2017;31(11).

4. Bouatou Y, Viglietti D, Pievani D, Louis K, Duong Van Huyen JP, Rabant M, et al. Response to treatment and long-term outcomes in kidney transplant recipients with acute T cell-mediated rejection. Am J Transplant. 2019;19(7):1972–88.

5. Rampersad C, Balshaw R, Gibson IW, Ho J, Shaw J, Karpinski M, et al. The negative impact of T cell-mediated rejection on renal allograft survival in the modern era. Am J Transplant. 2022;22(3):761–71.

6. Benichou G, and Kim J. Editorial: Allorecognition by Leukocytes of the Adaptive Immune System. Front Immunol. 2017;8:1555.

7. Mueller FB, Yang H, Lubetzky M, Verma A, Lee JR, Dadhania DM, et al. Landscape of innate immune system transcriptome and acute T cell-mediated rejection of human kidney allografts. JCI Insight. 2019;4(13).

8. Sarwal M, Chua MS, Kambham N, Hsieh SC, Satterwhite T, Masek M, et al. Molecular heterogeneity in acute renal allograft rejection identified by DNA microarray profiling. N Engl J Med. 2003;349(2):125–38.

9. Spivey TL, Uccellini L, Ascierto ML, Zoppoli G, De Giorgi V, Delogu LG, et al. Gene expression profiling in acute allograft rejection: challenging the immunologic constant of rejection hypothesis. J Transl Med. 2011;9:174.

10. Hennessy C, Lewik G, Cross A, Hester J, and Issa F. Recent advances in our understanding of the allograft response. Fac Rev. 2021;10:21.

11. Savas P, Virassamy B, Ye C, Salim A, Mintoff CP, Caramia F, et al. Single-cell profiling of breast cancer T cells reveals a tissue-resident memory subset associated with improved prognosis. Nat Med. 2018;24(7):986–93.

12. Ou Y, and Rattner JB. The centrosome in higher organisms: structure, composition, and duplication. Int Rev Cytol. 2004;238:119–82.

13. Yi JS, Cox MA, and Zajac AJ. T-cell exhaustion: characteristics, causes and conversion. Immunology. 2010;129(4):474–81.

14. Naba A, Clauser KR, Hoersch S, Liu H, Carr SA, and Hynes RO. The matrisome: in silico definition and in vivo characterization by proteomics of normal and tumor extracellular matrices. Mol Cell Proteomics. 2012;11(4):M111 014647.

15. Humphrey JD, Dufresne ER, and Schwartz MA. Mechanotransduction and extracellular matrix homeostasis. Nat Rev Mol Cell Biol. 2014;15(12):802–12.

16. Ji AL, Rubin AJ, Thrane K, Jiang S, Reynolds DL, Meyers RM, et al. Multimodal Analysis of Composition and Spatial Architecture in Human Squamous Cell Carcinoma. Cell. 2020;182(2):497–514 e22.

17. Franceschini A, Szklarczyk D, Frankild S, Kuhn M, Simonovic M, Roth A, et al. STRING v9.1: protein-protein interaction networks, with increased coverage and integration. Nucleic Acids Res. 2013;41(Database issue):D808–15.

18. Einecke G, Melk A, Ramassar V, Zhu LF, Bleackley RC, Famulski KS, et al. Expression of CTL associated transcripts precedes the development of tubulitis in T-cell mediated kidney graft rejection. Am J Transplant. 2005;5(8):1827–36.

19. Mueller TF, Einecke G, Reeve J, Sis B, Mengel M, Jhangri GS, et al. Microarray analysis of rejection in human kidney transplants using pathogenesis-based transcript sets. Am J Transplant. 2007;7(12):2712–22.

20. Muthukumar T, Dadhania D, Ding R, Snopkowski C, Naqvi R, Lee JB, et al. Messenger RNA for FOXP3 in the urine of renal-allograft recipients. N Engl J Med. 2005;353(22):2342–51.

21. Luan D, Dadhania DM, Ding R, Muthukumar T, Lubetzky M, Lee JR, et al. FOXP3 mRNA Profile Prognostic of Acute T Cell-mediated Rejection and Human Kidney Allograft Survival. Transplantation. 2021;105(8):1825–39.

22. Muller WA. Getting leukocytes to the site of inflammation. Vet Pathol. 2013;50(1):7–22.

23. Muller WA. Transendothelial migration: unifying principles from the endothelial perspective. Immunol Rev. 2016;273(1):61–75.

24. Cook-Mills JM, Marchese ME, and Abdala-Valencia H. Vascular cell adhesion molecule-1 expression and signaling during disease: regulation by reactive oxygen species and antioxidants. Antioxid Redox Signal. 2011;15(6):1607–38.

25. Matheny HE, Deem TL, and Cook-Mills JM. Lymphocyte migration through monolayers of endothelial cell lines involves VCAM-1 signaling via endothelial cell NADPH oxidase. J Immunol. 2000;164(12):6550–9.

26. Hughes AD, Lakkis FG, and Oberbarnscheidt MH. Four-Dimensional Imaging of T Cells in Kidney Transplant Rejection. J Am Soc Nephrol. 2018;29(6):1596–600.

27. Batal I, De Serres SA, Safa K, Bijol V, Ueno T, Onozato ML, et al. Dendritic Cells in Kidney Transplant Biopsy Samples Are Associated with T Cell Infiltration and Poor Allograft Survival. J Am Soc Nephrol. 2015;26(12):3102–13.

28. van Stipdonk MJ, Hardenberg G, Bijker MS, Lemmens EE, Droin NM, Green DR, et al. Dynamic programming of CD8+ T lymphocyte responses. Nat Immunol. 2003;4(4):361–5.

29. Verdon DJ, Mulazzani M, and Jenkins MR. Cellular and Molecular Mechanisms of CD8(+) T Cell Differentiation, Dysfunction and Exhaustion. Int J Mol Sci. 2020;21(19).

30. Smith KA. T-cell growth factor. Immunol Rev. 1980;51:337–57.

31. Laphanuwat P, and Jirawatnotai S. Immunomodulatory Roles of Cell Cycle Regulators. Front Cell Dev Biol. 2019;7:23.

32. Wang R, Dillon CP, Shi LZ, Milasta S, Carter R, Finkelstein D, et al. The transcription factor Myc controls metabolic reprogramming upon T lymphocyte activation. Immunity. 2011;35(6):871–82.

33. Man K, Miasari M, Shi W, Xin A, Henstridge DC, Preston S, et al. The transcription factor IRF4 is essential for TCR affinity-mediated metabolic programming and clonal expansion of T cells. Nat Immunol. 2013;14(11):1155–65.

34. Fox CJ, Hammerman PS, and Thompson CB. Fuel feeds function: energy metabolism and the T-cell response. Nat Rev Immunol. 2005;5(11):844–52.

35. Oberholtzer N, Quinn KM, Chakraborty P, and Mehrotra S. New Developments in T Cell Immunometabolism and Implications for Cancer Immunotherapy. Cells. 2022;11(4).

36. Kaech SM, Wherry EJ, and Ahmed R. Effector and memory T-cell differentiation: implications for vaccine development. Nat Rev Immunol. 2002;2(4):251–62.

37. Mason EF, and Rathmell JC. Cell metabolism: an essential link between cell growth and apoptosis. Biochim Biophys Acta. 2011;1813(4):645–54.

38. Traver D, Akashi K, Weissman IL, and Lagasse E. Mice defective in two apoptosis pathways in the myeloid lineage develop acute myeloblastic leukemia. Immunity. 1998;9(1):47–57.

39. Testa NG. Erythroid progenitor cells: their relevance for the study of haematological disease. Clin Haematol. 1979;8(2):311–33.

40. Moskophidis D, Lechner F, Pircher H, and Zinkernagel RM. Virus persistence in acutely infected immunocompetent mice by exhaustion of antiviral cytotoxic effector T cells. Nature. 1993;362(6422):758–61.

41. Starzl TE, and Zinkernagel RM. Antigen localization and migration in immunity and tolerance. N Engl J Med. 1998;339(26):1905–13.

42. Steger U, Denecke C, Sawitzki B, Karim M, Jones ND, and Wood KJ. Exhaustive differentiation of alloreactive CD8+ T cells: critical for determination of graft acceptance or rejection. Transplantation. 2008;85(9):1339–47.

43. Angeletti A, Cantarelli C, Riella LV, Fribourg M, and Cravedi P. T-cell Exhaustion in Organ Transplantation. Transplantation. 2022;106(3):489–99.

44. Fribourg M, Anderson L, Fischman C, Cantarelli C, Perin L, La Manna G, et al. T-cell exhaustion correlates with improved outcomes in kidney transplant recipients. Kidney Int. 2019;96(2):436–49.

45. Martins G, and Calame K. Regulation and functions of Blimp-1 in T and B lymphocytes. Annu Rev Immunol. 2008;26:133–69.

46. Haas M, Loupy A, Lefaucheur C, Roufosse C, Glotz D, Seron D, et al. The Banff 2017 Kidney Meeting Report: Revised diagnostic criteria for chronic active T cell-mediated rejection, antibody-mediated rejection, and prospects for integrative endpoints for next-generation clinical trials. Am J Transplant. 2018;18(2):293–307.

47. Racusen LC, Solez K, Colvin RB, Bonsib SM, Castro MC, Cavallo T, et al. The Banff 97 working classification of renal allograft pathology. Kidney Int. 1999;55(2):713–23.

48. Andrews S. FastQC: a quality control tool for high throughput sequence data. http://www.bioinformatics.babraham.ac.uk/projects/fastqc.

49. Bolger AM, Lohse M, and Usadel B. Trimmomatic: a flexible trimmer for Illumina sequence data. Bioinformatics. 2014;30(15):2114–20.

50. Kim D, Pertea G, Trapnell C, Pimentel H, Kelley R, and Salzberg SL. TopHat2: accurate alignment of transcriptomes in the presence of insertions, deletions and gene fusions. Genome Biol. 2013;14(4):R36.

51. Trapnell C, Roberts A, Goff L, Pertea G, Kim D, Kelley DR, et al. Differential gene and transcript expression analysis of RNA-seq experiments with TopHat and Cufflinks. Nat Protoc. 2012;7(3):562–78.

52. Anders S, Pyl PT, and Huber W. HTSeq--a Python framework to work with high-throughput sequencing data. Bioinformatics. 2015;31(2):166–9.

53. Robinson MD, and Oshlack A. A scaling normalization method for differential expression analysis of RNA-seq data. Genome Biol. 2010;11(3):R25.

54. Luo W, Friedman MS, Shedden K, Hankenson KD, and Woolf PJ. GAGE: generally applicable gene set enrichment for pathway analysis. BMC Bioinformatics. 2009;10:161.

55. Ritchie ME, Phipson B, Wu D, Hu Y, Law CW, Shi W, et al. limma powers differential expression analyses for RNA-sequencing and microarray studies. Nucleic Acids Res. 2015;43(7):e47.

56. Luo W, and Brouwer C. Pathview: an R/Bioconductor package for pathway-based data integration and visualization. Bioinformatics. 2013;29(14):1830–1.

57. Zhang B, and Horvath S. A general framework for weighted gene co-expression network analysis. Stat Appl Genet Mol Biol. 2005;4:Article17.

58. Yu G, and He QY. ReactomePA: an R/Bioconductor package for reactome pathway analysis and visualization. Mol Biosyst. 2016;12(2):477–9.

59. Wu D, Lim E, Vaillant F, Asselin-Labat ML, Visvader JE, and Smyth GK. ROAST: rotation gene set tests for complex microarray experiments. Bioinformatics. 2010;26(17):2176–82.

60. Wu D, and Smyth GK. Camera: a competitive gene set test accounting for inter-gene correlation. Nucleic Acids Res. 2012;40(17):e133.

61. Korotkevich G, Sukhov V, Budin N, Shpak B, Artyomov MN, and Sergushichev A. Fast gene set enrichment analysis. bioRxiv. 2021:060012.

62. Langfelder P, and Horvath S. WGCNA: an R package for weighted correlation network analysis. BMC Bioinformatics. 2008;9:559.

63. Robinson MD, McCarthy DJ, and Smyth GK. edgeR: a Bioconductor package for differential expression analysis of digital gene expression data. Bioinformatics. 2010;26(1):139–40.

64. World Medical A. World Medical Association Declaration of Helsinki: ethical principles for medical research involving human subjects. JAMA. 2013;310(20):2191–4.

65. International Summit on Transplant T, and Organ T. The Declaration of Istanbul on Organ Trafficking and Transplant Tourism. Clin J Am Soc Nephrol. 2008;3(5):1227–31.

