## Supplemental Figures & Tables for "Adaptive Immune Landscape of T-Cell Mediated Rejection of Human Kidney Allografts"

Supplemental Figure S1. Heatmaps of Major Histocompatibility I and II Pathway Genes.

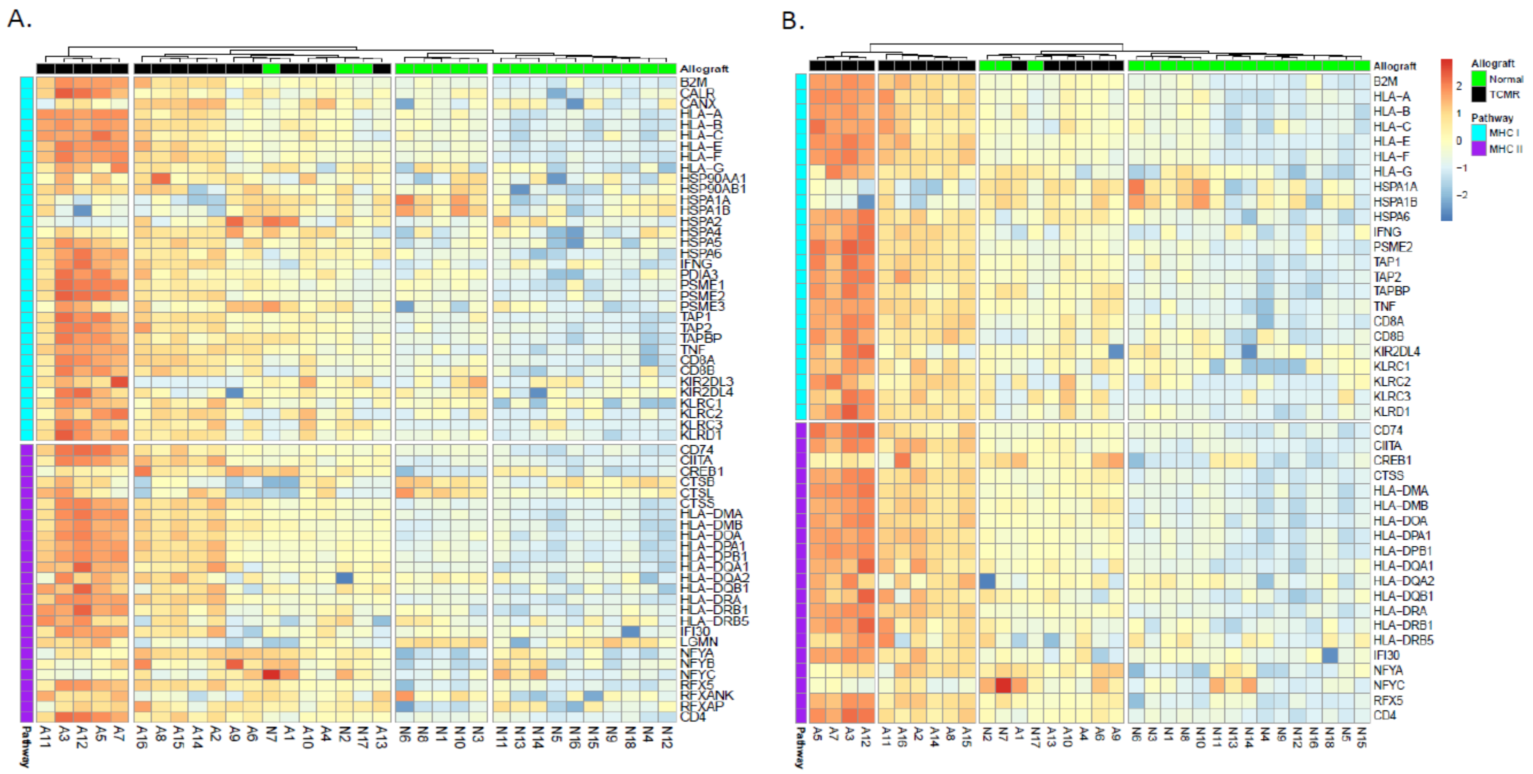


Heatmaps of Major Histocompatibility I and II KEGG Pathways. (A) Gene expression of MHC I and MHC II pathway, (B) Differentially expressed genes only at P-FDR <0.05.

Supplemental Figure S2. Heatmaps of Autosomal Minor Histocompatibility Antigens.

A.

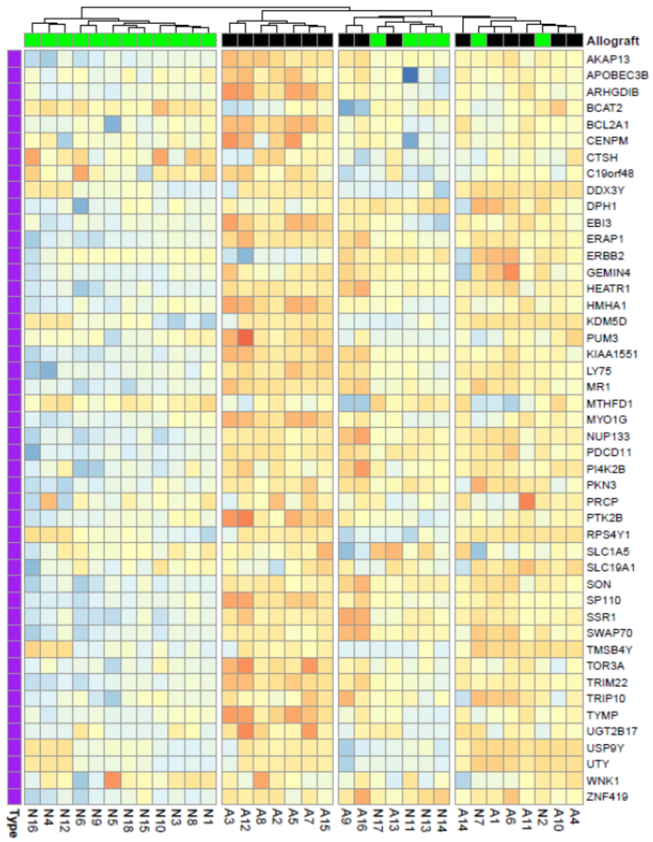

B.

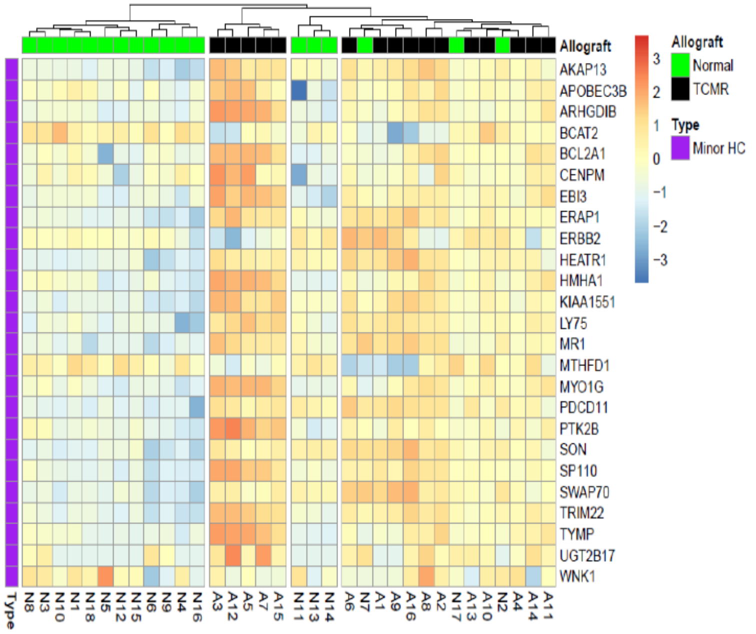

Heatmaps of Autosomal Minor Histocompatibility Antigens Pathways. (A.) mRNA expression of all pathway genes. (B.) Differentially expressed genes only (P-FDR <0.05).

Supplemental Figure S3. Relation Between TCR Signaling and Th1, Th2, and Th17 Differentiation Pathways.

A.

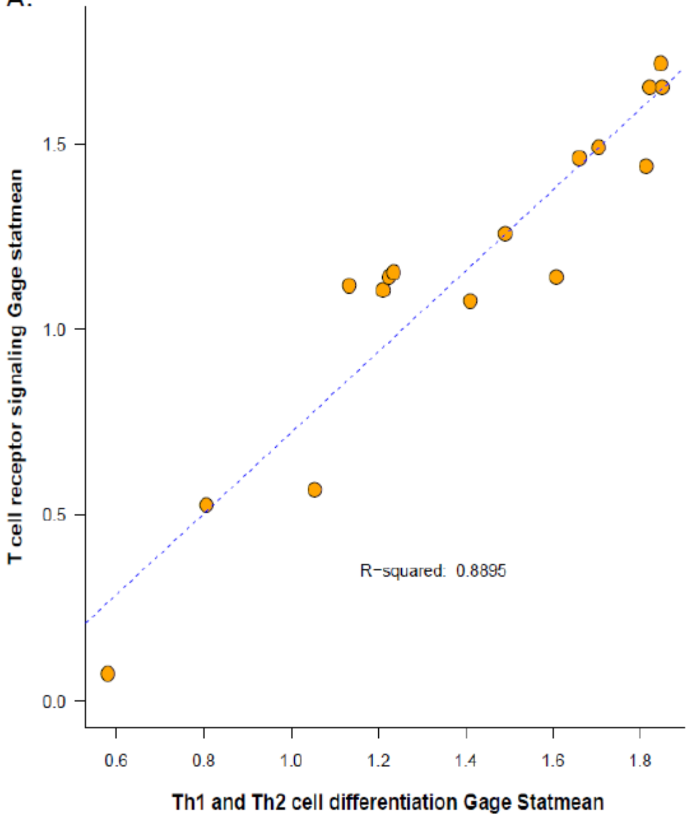


B.

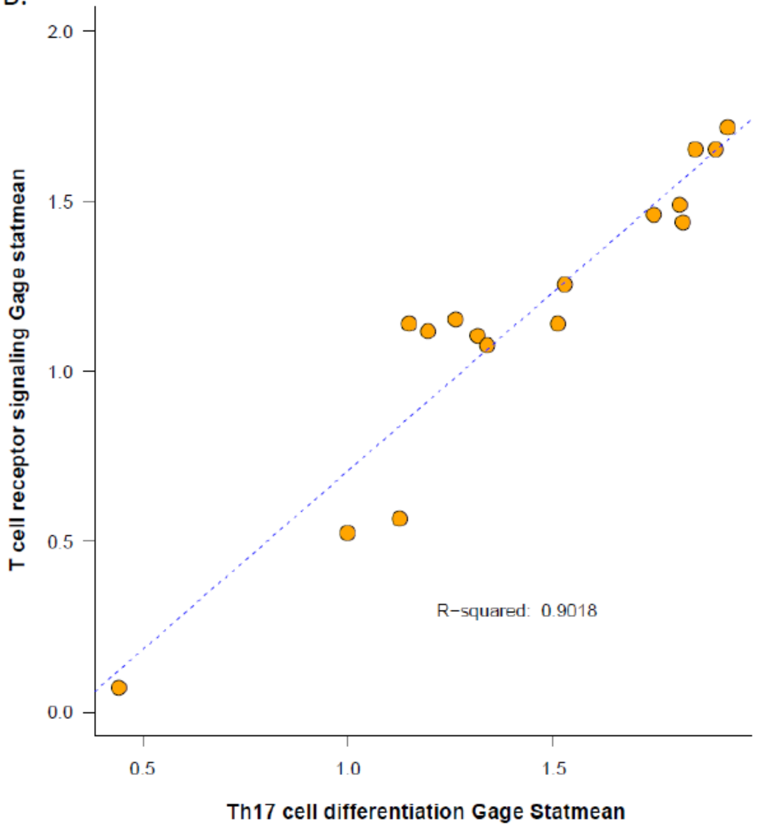


KEGG pathway GAGE enrichment mean (stat-mean) of Th1/Th2 (A) and Th17 cell differentiation (B) against the stat-mean of T cell receptor signaling pathway. The stat mean is the mean of the individual statistics from multiple gene-set tests. Normally, its absolute value measures the magnitude of gene-set level changes, and its sign indicates direction of the changes.

Supplemental Figure S4. Heatmap of p53 Signaling Pathways Genes.

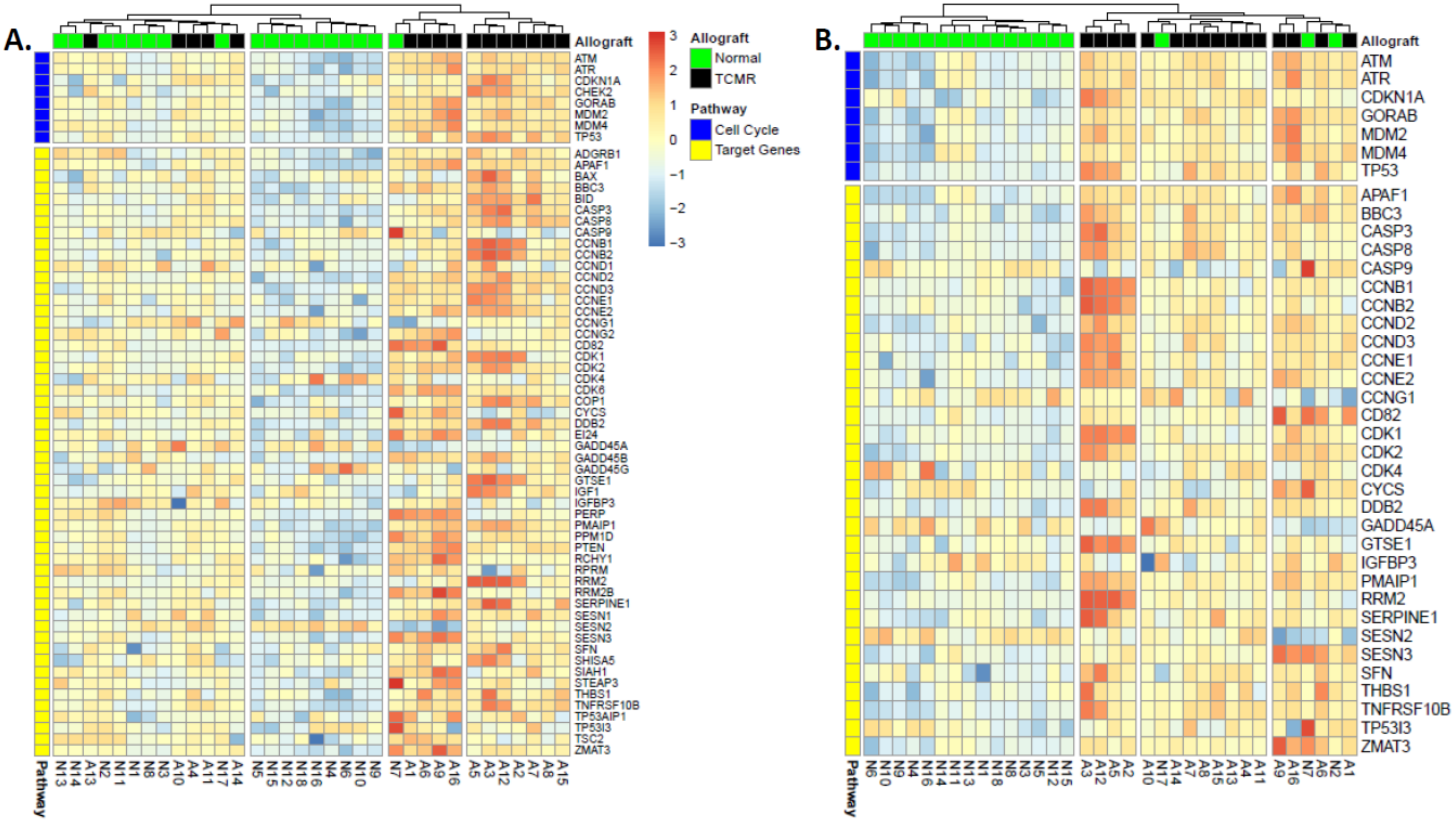


Heatmaps of all KEGG (A) and differentially expressed p53 pathway signaling genes (B).

Supplemental Figure 5. Heatmap of Apoptosis Pathways Genes.

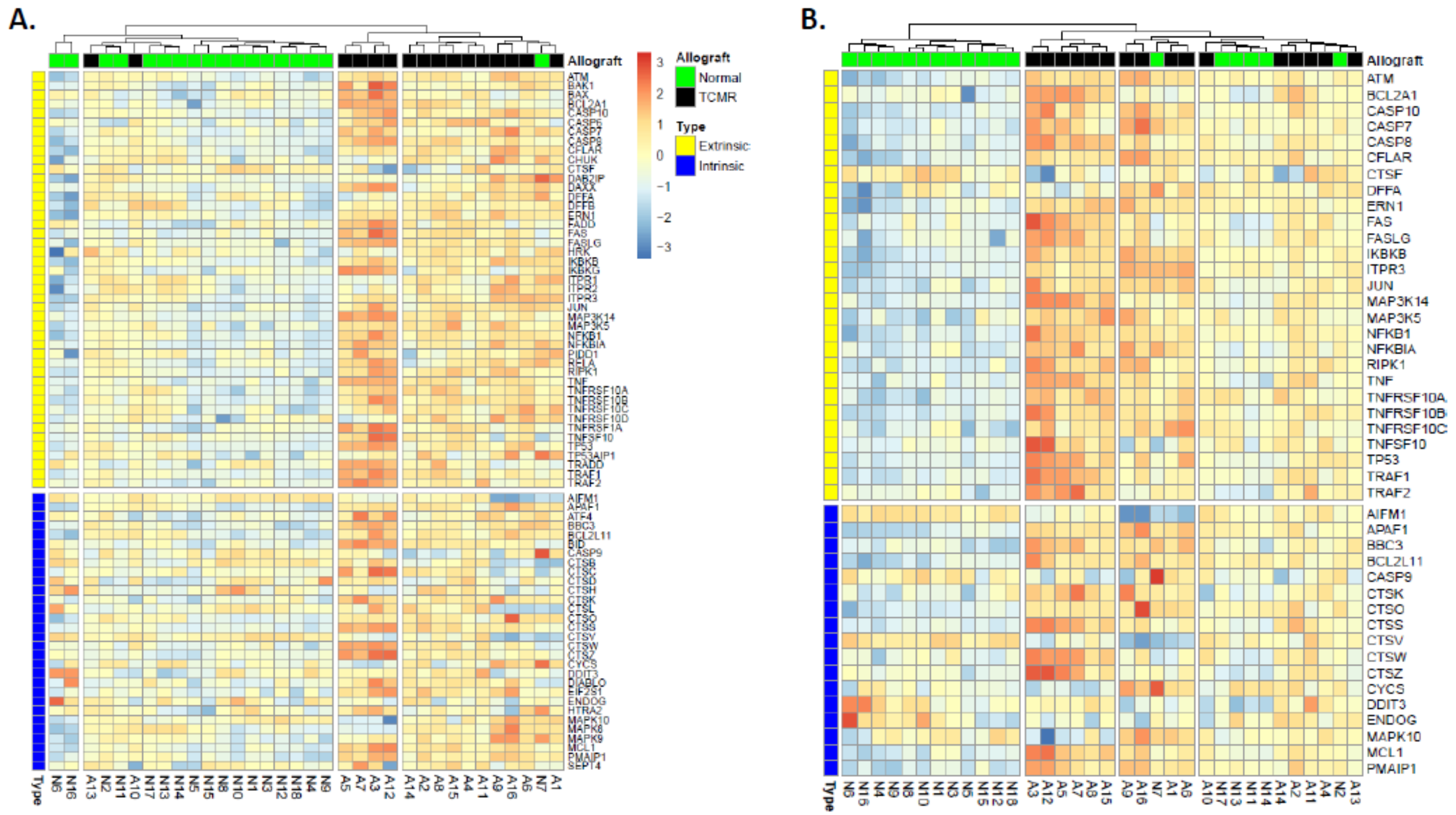

Heatmaps of all KEGG (A) and differentially expressed (B) extrinsic and intrinsic apoptosis pathway signaling genes.

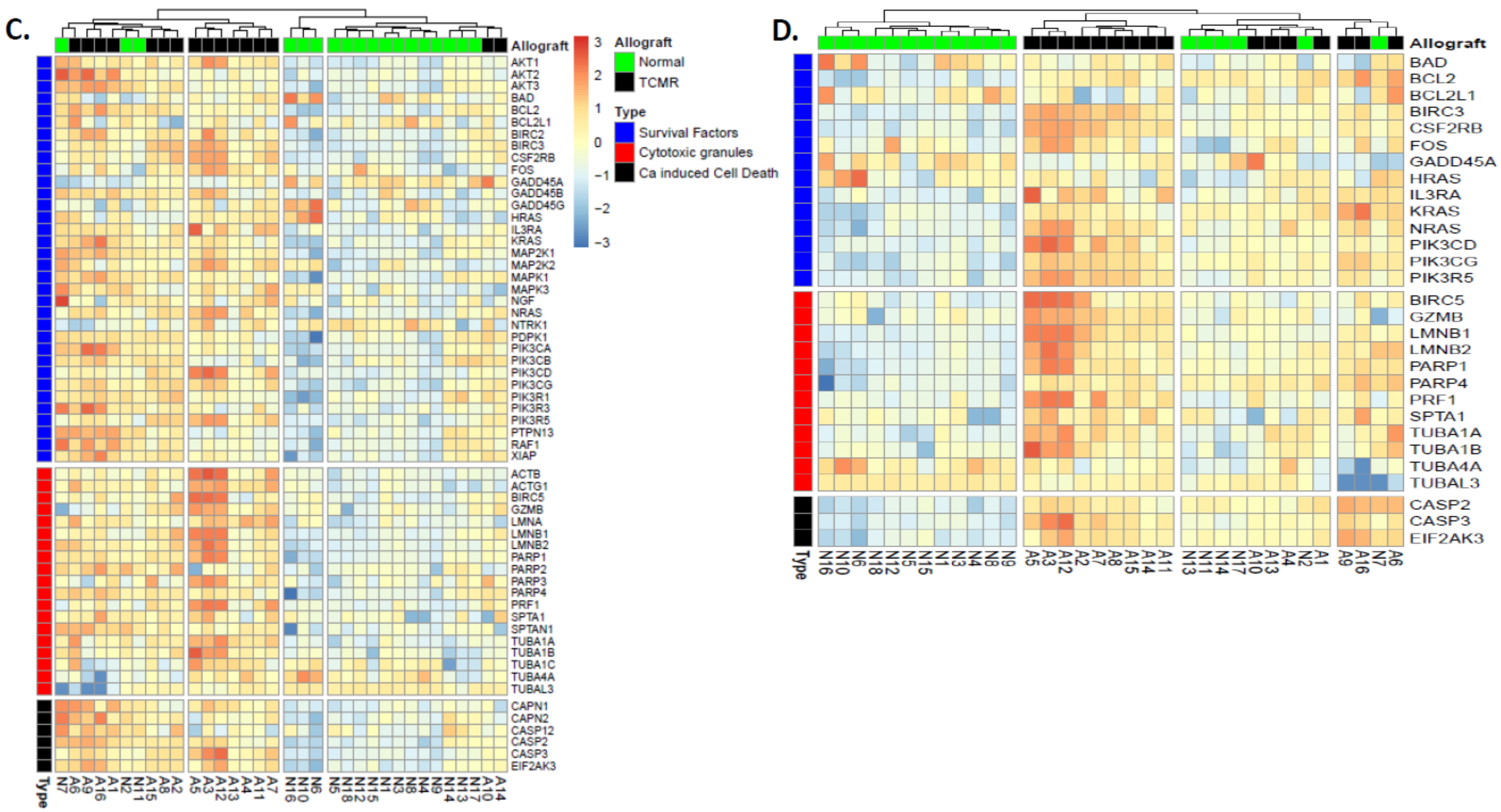


Heatmaps of all KEGG (C) and differentially expressed (D) survival factors, cytotoxic granules and calcium induced cell death gene of apoptosis pathway signaling genes.

Supplemental Figure 6. Abundance of Granzyme and Perforin mRNA in TCMR.

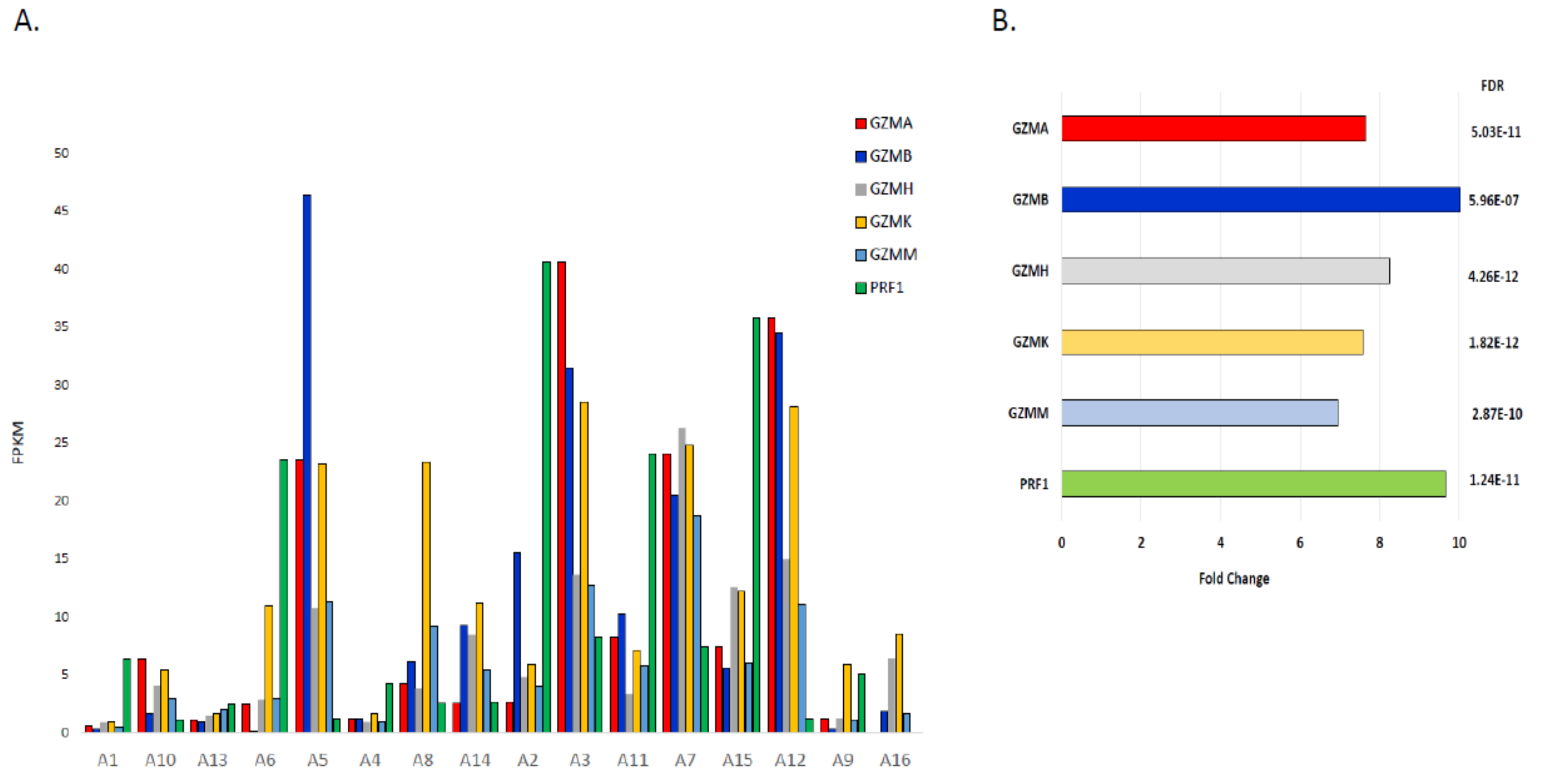

Supplemental Figure S7. Heatmap of Differentially Expressed Necroptosis Pathway Gene and Correlation of Cytosolic DNA Sensing Pathway with Necroptosis Signaling Pathway Enrichment in TCMR.

A.

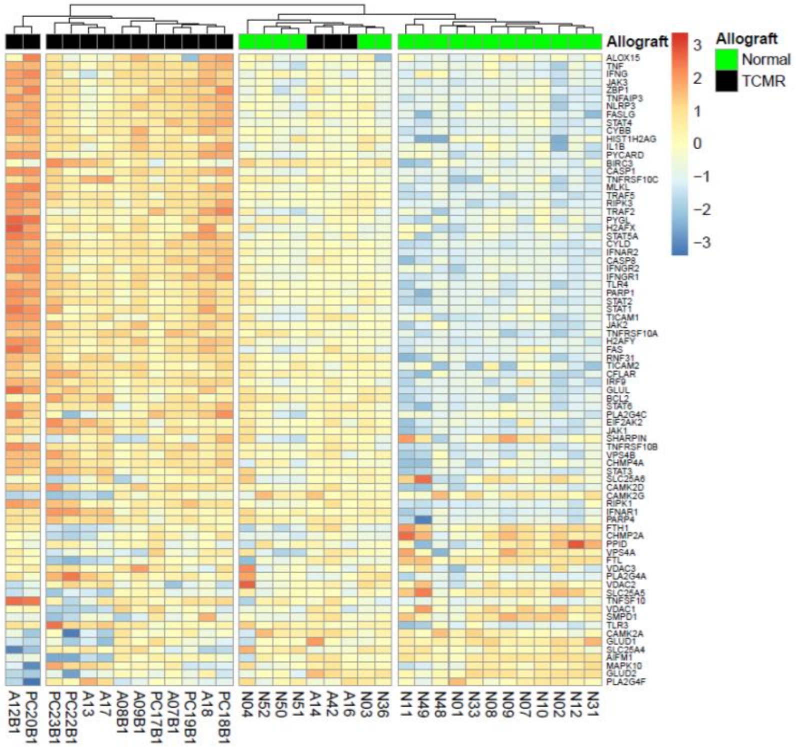


B.

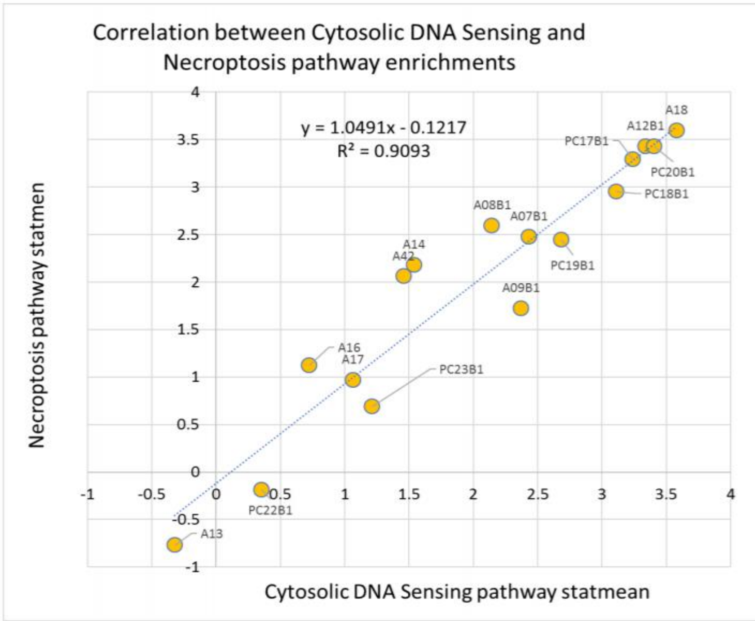


(A) Heatmap of differentially expressed necroptosis KEGG pathway genes. (B) Relationship of GAGE enriched cytosolic DNA sensing pathway stat-mean in TCMR to necroptosis pathway stat-mean.

Supplemental Figure 8.  Impact of Time Transplantation to Biopsy on Matrisome Gene Expression.

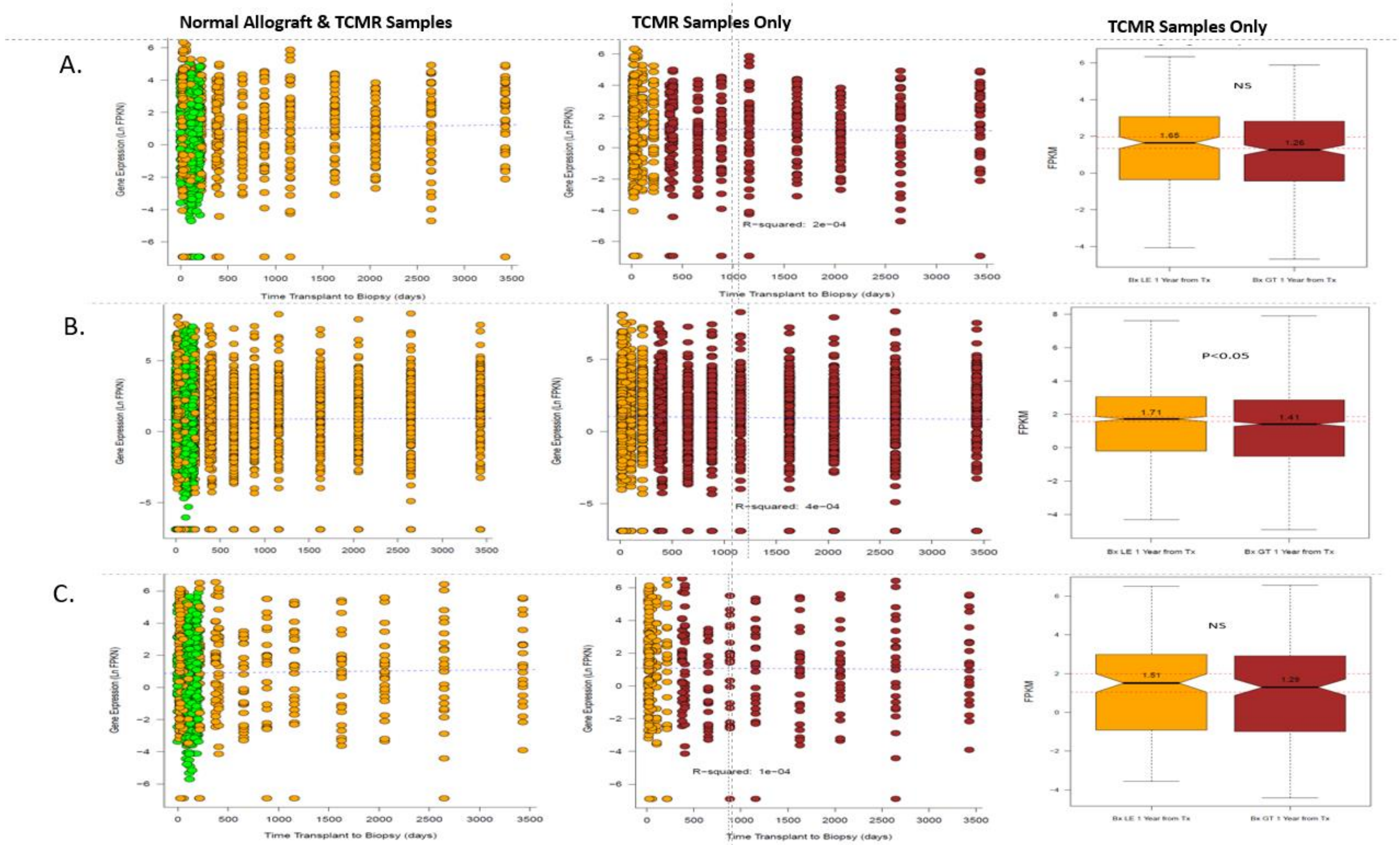


(A) Collagen genes, (B) Glycoprotein genes and (C) Proteoglycan genes. Boxplots show genes expressions for biopsies performed 1 year or less after transplantation (orange) or greater than 1 year after transplantation.

Supplemental Figure S9. Differential Expression Analysis in Interstitial Fibrosis positive and negative TCMR Samples compared to Normal Allografts.

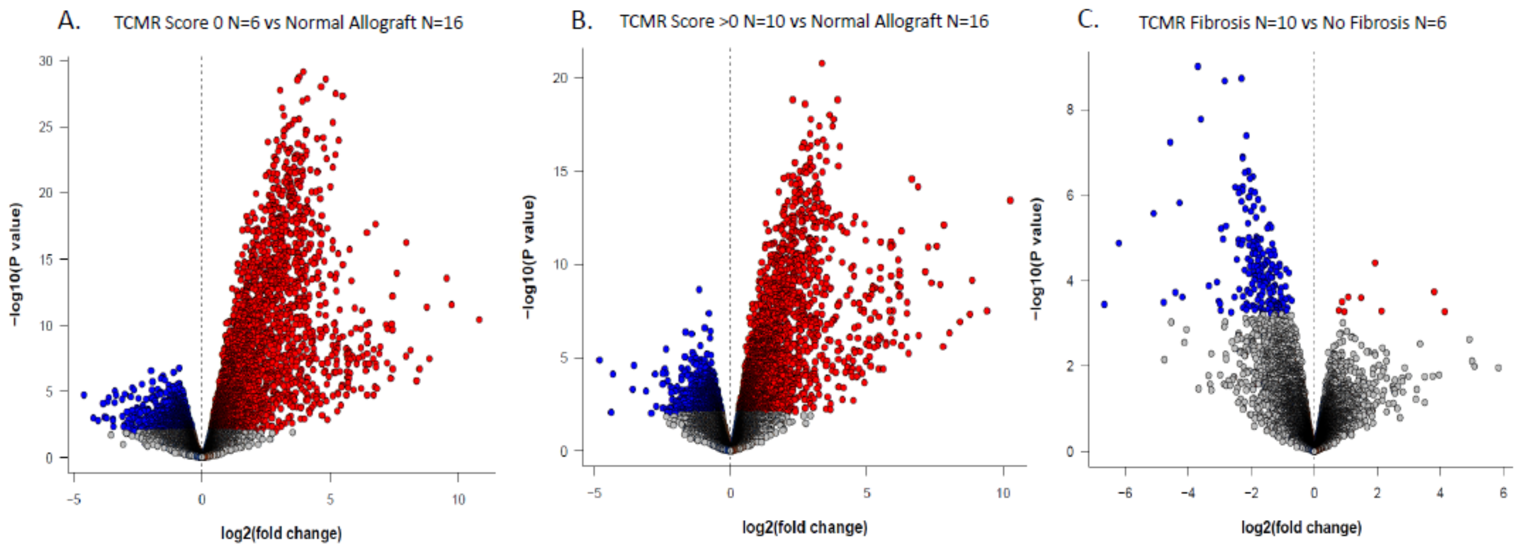


| Banff TCM Fibrosis Score 0 vs Normal Allograft | | | |
|---|---|---|---|
| | FDR Significance Level | | |
| Gene Expression | p<0.05 | p<0.01 | p<0.001 |
| Decreased in Biopsies | 1,273 | 524 | 120 |
| Increased | 2,844 | 2,253 | 1,778 |
| Differentially Expressed | **4,117** | **2,777** | **1,898** |
| Unchanged | 12,242 | 13,582 | 14,461 |
| % DE | 33.63% | 20.45% | 13.12% |
| Dispersion | 19.46% | | |
| BCV | 44.11% | | |

| Banff TCMR Fibrosis Score >0 vs Normal Allograft | | | |
|---|---|---|---|
| | FDR Significance Level | | |
| Gene Expression | p<0.05 | p<0.01 | p<0.001 |
| Decreased in Biopsies | 1,129 | 328 | 51 |
| Increased | 2,451 | 1,646 | 1,149 |
| Differentially Expressed | **3,580** | **1,974** | **1,200** |
| Unchanged | 12,779 | 14,385 | 15,159 |
| % DE | 28.01% | 13.72% | 7.92% |
| Dispersion | 19.46% | | |
| BCV | 44.11% | | |

| Banff TCMR Fibrosis Score >0 vs TCMR Fibrosis Score 0 | | | |
|---|---|---|---|
| | FDR Significance Level | | |
| Gene Expression | p<0.05 | p<0.01 | p<0.001 |
| Decreased in Biopsies | 195 | 84 | 22 |
| Increased | 9 | 1 | 0 |
| Differentially Expressed | **204** | **85** | **22** |
| Unchanged | 16,155 | 16,274 | 16,337 |
| % DE | 1.26% | 0.52% | 0.13% |
| Dispersion | 19.46% | | |
| BCV | 44.11% | | |

Volcano plots of differentially expressed genes in TCMR stratified by Banff interstitial fibrosis score. Differentially expressed genes (P-FDR<0.05) with positive $\log_2$(fold change) of ≥2 is indicated in red; with negative $\log_2$(fold change) of <2 is indicated in blue. Non-significant genes are in grey. (A) Comparison of Normal allografts with no fibrosis (Banff ci + ct score = 0, N=16) to TCMR samples with no fibrosis (Banff ci + ct score = 0, N=6). (B) Comparison of Normal allografts with no fibrosis (Banff ci + ct score = 0, N=16) to TCMR samples with fibrosis (Banff ci + ct score >0, mean score 1.3± 0.67 SD, N=10). (C) Comparison between TCMR samples with fibrosis and TCMR samples without fibrosis.

### Adaptive Immune Landscape of T-Cell Mediated Rejection of Human Kidney Allografts

#### Supplemental Tables

Supplemental Table S1. Characteristics of Kidney Transplant Recipients.

| Characteristics | Kidney Allograft Biopsies | |
|---|---|---|
| | **Acute T Cell Mediated Rejection (N=16)** | **Normal / Non-Specific (N=18)** |
| ***At the time of transplantation*** | | |
| Recipient information | | |
| Age, years, mean (SD) | 47 (12) | 49 (12) |
| Women, n (%) | 8 (50) | 9 (50) |
| Black, n (%) | 6 (38) | 4 (22) |
| Donor information | | |
| Age, years, mean (SD) | 42 (17) | 46 (13) |
| Women, n (%) | 10 (63) | 14 (78) |
| Black, n (%) | 1 (16) | 1 (6) |
| Deceased donor, n (%) | 8 (50) | 7 (39) |
| HLA-ABDR mismatch, median (IQR) | 5 (3-6) | 5 (3-6) |
| Induction therapy, n (%) | 16 (100) | 18 (100) |
| Antithymocyte globulin, n (%) | 12 (75) | 18 (100) |
| ***After transplantation and before the index biopsy*** | | |
| Delayed graft function, n (%) | 4 (25) | 0 (0) |
| Tacrolimus, n (%) | 15 (94)* | 18 (100) |
| Mycophenolate Mofetil, n (%) | 16 (100) | 18 (100) |
| Corticosteroid maintenance, n (%) | 3 (19) | 2 (11) |
| BK virus nephropathy, n (%) | 0 (0) | 0 (0) |
| Biopsy proven acute rejection, n (%) | 0 (0) | 0 (0) |
| ***At the time of index allograft biopsy*** | | |
| Clinically indicated: Surveillance biopsy, n | 16: 0 | 0:18 |
| Time from transplantation to biopsy, months, median (IQR) | 12.5 (1-49) | 3.5 (3-6) |
| Serum creatinine, mg/dl, median (IQR) | 2.55 (1.77-3.83) | 1.34 (1.05-1.48) |
| Serum tacrolimus trough, ng/dl, median (IQR) | 5.2 (4.2-6.7) | 7.0 (6.0-8.7) |

* One recipient received ASKP1240, a fully human antibody targeting CD40 in antigen presenting cells.

Supplemental Table S2. Intragraft Expression of mRNAs Encoding Antigen Processing and Presentation KEGG Pathway Genes in TCMR and Normal Kidney Allograft Biopsies.

Table S2A. MHC 1 Pathway

| | Normal Allograft | | TCMR | | | | |
|---|---|---|---|---|---|---|---|
| Symbol | Median | IQR | Median | IQR | FC | PValue | FDR |
| B2M | 2196.5 | 737.7 | 8176.2 | 7549.0 | 2.7 | 5.6E-08 | 8.7E-07 |
| CALR | 244.3 | 34.2 | 290.7 | 90.9 | 1.1 | 5.1E-01 | NS |
| CANX | 108.4 | 21.6 | 130.1 | 32.2 | 1.0 | 9.3E-01 | NS |
| HLA-A | 369.4 | 164.9 | 831.7 | 1068.1 | 2.4 | 4.8E-06 | 4.6E-05 |
| HLA-B | 408.3 | 255.3 | 1355.3 | 1524.6 | 3.1 | 6.5E-09 | 1.3E-07 |
| HLA-C | 366.7 | 186.1 | 743.1 | 872.2 | 2.2 | 3.4E-05 | 2.5E-04 |
| HLA-E | 161.6 | 58.4 | 352.1 | 346.0 | 2.1 | 1.4E-06 | 1.5E-05 |
| HLA-F | 45.6 | 20.4 | 146.0 | 197.8 | 3.5 | 1.7E-12 | 6.8E-11 |
| HLA-G | 5.7 | 4.5 | 13.7 | 13.9 | 2.3 | 5.1E-03 | 1.7E-02 |
| HSP90AA1 | 201.4 | 41.3 | 211.5 | 31.1 | -1.1 | 5.1E-01 | NS |
| HSP90AB1 | 200.9 | 27.1 | 224.5 | 31.0 | -1.1 | 3.6E-01 | NS |
| HSPA1A | 76.0 | 21.2 | 68.7 | 26.6 | -1.4 | 3.2E-03 | 1.1E-02 |
| HSPA1B | 68.3 | 23.9 | 64.5 | 21.2 | -1.5 | 6.6E-04 | 3.1E-03 |
| HSPA2 | 14.8 | 19.2 | 14.2 | 38.2 | 1.1 | 7.7E-01 | NS |
| HSPA4 | 19.7 | 2.9 | 21.0 | 2.1 | -1.1 | 2.6E-01 | NS |
| HSPA5 | 75.5 | 18.2 | 99.5 | 18.3 | 1.2 | 1.2E-01 | NS |
| HSPA6 | 0.8 | 0.5 | 3.1 | 3.1 | 3.6 | 1.8E-10 | 4.7E-09 |
| IFNG | 0.1 | 0.2 | 1.2 | 1.8 | 8.1 | 2.2E-08 | 3.8E-07 |
| PDIA3 | 146.2 | 21.9 | 163.4 | 41.4 | 1.1 | 5.2E-01 | NS |
| PSME1 | 141.0 | 37.2 | 193.9 | 103.1 | 1.3 | 3.5E-02 | NS |
| PSME2 | 83.2 | 102.3 | 349.4 | 489.7 | 1.7 | 7.8E-05 | 5.1E-04 |
| PSME3 | 20.9 | 3.7 | 23.0 | 4.6 | -1.0 | 8.5E-01 | NS |
| TAP1 | 16.9 | 7.0 | 72.1 | 58.4 | 4.2 | 1.5E-13 | 7.3E-12 |
| TAP2 | 9.5 | 4.6 | 17.7 | 16.1 | 3.1 | 5.5E-14 | 3.1E-12 |
| TAPBP | 21.8 | 5.6 | 46.6 | 23.4 | 1.9 | 2.8E-08 | 4.7E-07 |
| TNF | 0.4 | 0.3 | 2.4 | 2.0 | 5.3 | 2.3E-14 | 1.4E-12 |

Table S2B. MHC 1 Pathway

| Symbol | Normal Allograft Median | Normal Allograft IQR | TCMR Median | TCMR IQR | FC | PValue | FDR |
|---|---|---|---|---|---|---|---|
| CD74 | 782.0 | 362.6 | 1858.1 | 2497.0 | 3.1 | 6.4E-08 | 9.9E-07 |
| CIITA | 1.6 | 0.9 | 13.2 | 14.7 | 2.6 | 2.0E-12 | 7.9E-11 |
| CREB1 | 2.7 | 1.7 | 4.4 | 1.4 | 1.3 | 7.9E-04 | 3.6E-03 |
| CTSB | 689.6 | 432.1 | 497.7 | 584.7 | -1.5 | 5.1E-02 | NS |
| CTSL | 137.5 | 32.4 | 133.8 | 72.9 | -1.3 | 5.6E-02 | NS |
| CTSS | 9.6 | 4.9 | 85.8 | 126.7 | 8.3 | 1.3E-22 | 5.5E-19 |
| HLA-DMA | 0.0 | 56.9 | 146.2 | 107.5 | 2.8 | 1.1E-12 | 4.8E-11 |
| HLA-DMB | 0.0 | 0.0 | 124.1 | 90.3 | 4.1 | 4.2E-20 | 2.2E-17 |
| HLA-DOA | 5.7 | 3.4 | 25.8 | 27.2 | 4.3 | 6.9E-17 | 9.9E-15 |
| HLA-DPA1 | 111.1 | 59.0 | 532.3 | 690.3 | 4.5 | 8.4E-14 | 4.5E-12 |
| HLA-DPB1 | 83.9 | 47.2 | 462.8 | 552.5 | 4.3 | 1.5E-13 | 7.3E-12 |
| HLA-DQA1 | 23.6 | 10.9 | 139.3 | 255.8 | 7.2 | 2.8E-13 | 1.3E-11 |
| HLA-DQA2 | 4.4 | 4.7 | 19.0 | 37.0 | 4.1 | 5.1E-06 | 4.9E-05 |
| HLA-DQB1 | 22.5 | 26.2 | 118.9 | 344.7 | 7.5 | 1.2E-08 | 2.3E-07 |
| HLA-DRA | 211.8 | 89.9 | 1137.8 | 1243.4 | 4.7 | 2.3E-14 | 1.4E-12 |
| HLA-DRB1 | 139.6 | 90.7 | 300.9 | 576.4 | 2.8 | 1.4E-06 | 1.6E-05 |
| HLA-DRB5 | 36.2 | 47.2 | 125.7 | 155.0 | 2.7 | 5.6E-03 | 1.8E-02 |
| IFI30 | 66.9 | 29.1 | 125.3 | 168.3 | 4.7 | 6.8E-07 | 8.2E-06 |
| LGMN | 207.6 | 142.0 | 93.2 | 164.3 | -1.6 | 4.1E-02 | NS |
| NFYA | 5.6 | 8.2 | 10.1 | 2.7 | 1.4 | 2.2E-03 | 8.4E-03 |
| NFYB | 9.5 | 7.7 | 16.6 | 8.4 | -1.0 | 1.0E+00 | NS |
| NFYC | 24.8 | 5.2 | 24.3 | 5.2 | -1.2 | 3.2E-04 | 1.7E-03 |
| RFX5 | 9.5 | 2.7 | 17.9 | 6.2 | 1.7 | 4.9E-09 | 9.7E-08 |
| RFXANK | 20.1 | 5.9 | 22.2 | 4.5 | -1.2 | 1.5E-01 | NS |
| RFXAP | 1.6 | 0.7 | 2.1 | 0.4 | 1.1 | 4.4E-01 | NS |
| CD4 | 6.6 | 2.1 | 20.8 | 27.5 | 4.3 | 1.4E-14 | 9.4E-13 |

Table S2C. Autosomal Minor Histocompatibility Antigens

| Symbol | Normal Allograft Median | Normal Allograft IQR | TCMR Median | TCMR IQR | FC | PValue | FDR |
|---|---|---|---|---|---|---|---|
| MYO1G | 0.7 | 0.5 | 5.0 | 8.7 | 7.4 | 9.7E-17 | 1.3E-14 |
| TRIM22 | 11.5 | 6.5 | 38.9 | 27.6 | 3.3 | 1.3E-16 | 1.7E-14 |
| TYMP | 6.3 | 4.7 | 36.5 | 89.6 | 7.8 | 6.6E-16 | 6.6E-14 |
| KIAA1551 | 8.1 | 2.8 | 16.6 | 20.9 | 2.7 | 1.8E-15 | 1.5E-13 |
| HMHA1 | 0.0 | 0.0 | 17.9 | 32.4 | 3.3 | 2.7E-12 | 1.0E-10 |
| BCL2A1 | 0.5 | 0.2 | 4.2 | 9.4 | 9.4 | 1.5E-11 | 5.0E-10 |
| SP110 | 4.9 | 1.7 | 11.5 | 7.6 | 2.2 | 3.2E-11 | 9.9E-10 |
| EBI3 | 1.0 | 0.7 | 3.5 | 4.5 | 4.2 | 4.4E-11 | 1.3E-09 |
| ERAP1 | 8.6 | 5.0 | 22.2 | 11.8 | 1.8 | 1.2E-10 | 3.4E-09 |
| LY75 | 0.0 | 2.4 | 5.0 | 2.6 | 2.6 | 3.1E-10 | 7.9E-09 |
| ARHGDIB | 41.4 | 13.3 | 98.1 | 99.4 | 2.7 | 1.4E-09 | 3.2E-08 |
| MR1 | 3.8 | 0.9 | 7.3 | 2.7 | 1.7 | 6.2E-07 | 7.6E-06 |
| AKAP13 | 6.8 | 2.5 | 12.1 | 3.8 | 1.5 | 1.2E-06 | 1.4E-05 |
| APOBEC3B | 0.6 | 0.5 | 1.9 | 2.2 | 3.0 | 1.3E-05 | 1.1E-04 |
| BCAT2 | 41.3 | 9.5 | 21.7 | 7.8 | -1.6 | 1.6E-05 | 1.3E-04 |
| HEATR1 | 3.4 | 2.2 | 6.2 | 1.9 | 1.4 | 3.8E-05 | 2.7E-04 |
| MTHFD1 | 50.3 | 13.8 | 29.6 | 15.5 | -1.5 | 2.5E-04 | 1.4E-03 |
| UGT2B17 | 0.1 | 0.0 | 0.1 | 0.1 | 4.4 | 1.6E-03 | 6.6E-03 |
| PTK2B | 10.6 | 2.0 | 15.3 | 7.9 | 1.4 | 1.8E-03 | 7.1E-03 |
| CENPM | 0.9 | 0.5 | 1.5 | 1.2 | 2.2 | 4.1E-03 | 1.4E-02 |
| SWAP70 | 6.6 | 4.9 | 11.1 | 5.3 | 1.3 | 6.2E-03 | 2.0E-02 |
| SON | 37.4 | 26.6 | 66.9 | 48.0 | 1.3 | 1.0E-02 | 2.9E-02 |
| ERBB2 | 69.6 | 21.9 | 60.4 | 22.2 | -1.3 | 1.1E-02 | 3.2E-02 |
| PDCD11 | 3.8 | 1.1 | 5.8 | 0.6 | 1.2 | 1.2E-02 | 3.4E-02 |
| WNK1 | 22.3 | 9.3 | 18.0 | 6.3 | -1.3 | 1.5E-02 | 4.1E-02 |
| CTSH | 464.1 | 227.9 | 449.6 | 146.4 | -1.4 | 2.7E-02 | NS |
| DPH1 | 19.4 | 7.3 | 19.0 | 4.3 | -1.2 | 2.8E-02 | NS |
| TOR3A | 8.8 | 1.3 | 12.6 | 3.0 | 1.2 | 2.9E-02 | NS |
| C19orf48 | 10.7 | 3.1 | 10.4 | 3.6 | -1.3 | 3.2E-02 | NS |
| NUP133 | 7.3 | 2.1 | 9.3 | 1.8 | 1.1 | 4.0E-02 | NS |
| PKN3 | 1.9 | 1.3 | 3.6 | 1.3 | 1.3 | 6.8E-02 | NS |
| SSR1 | 20.5 | 3.6 | 32.4 | 12.5 | 1.1 | 6.8E-02 | NS |
| PRCP | 76.8 | 17.8 | 80.6 | 22.4 | -1.2 | 1.6E-01 | NS |
| KDM5D | 0.3 | 5.2 | 6.4 | 7.0 | 1.9 | 2.2E-01 | NS |
| DDX3Y | 0.2 | 5.5 | 6.2 | 8.0 | 1.7 | 3.1E-01 | NS |
| UTY | 0.1 | 1.2 | 1.1 | 1.3 | 1.6 | 3.3E-01 | NS |
| PUM3 | 7.9 | 1.8 | 9.9 | 3.3 | 1.1 | 3.3E-01 | NS |
| TRIP10 | 25.9 | 6.8 | 31.9 | 11.8 | 1.1 | 3.8E-01 | NS |
| USP9Y | 0.1 | 6.7 | 4.7 | 9.7 | 1.6 | 3.8E-01 | NS |
| SLC19A1 | 103.5 | 22.4 | 117.0 | 37.5 | -1.1 | 4.4E-01 | NS |
| ZNF419 | 3.2 | 1.1 | 4.1 | 0.7 | 1.1 | 5.4E-01 | NS |
| SLC1A5 | 33.2 | 18.9 | 30.4 | 21.6 | -1.1 | 5.4E-01 | NS |
| PI4K2B | 5.8 | 1.1 | 7.2 | 1.2 | 1.0 | 6.8E-01 | NS |
| GEMIN4 | 3.6 | 0.5 | 4.4 | 0.9 | 1.0 | 8.2E-01 | NS |
| TMSB4Y | 0.1 | 0.9 | 0.6 | 0.6 | -1.1 | 8.7E-01 | NS |
| RPS4Y1 | 3.1 | 80.8 | 60.1 | 107.2 | 1.0 | 9.9E-01 | NS |

Supplemental Table S3. Intragraft Expression of mRNAs Encoding of T Cell Receptor Signaling KEGG Pathway in TCMR and Normal Kidney Allograft Biopsies.

Table S3A. T Cell Receptor Activation (Signal 1)

| | Normal Allograft | | TCMR | | | | |
|---|---|---|---|---|---|---|---|
| Symbol | Median | IQR | Median | IQR | FC | PValue | FDR |
| CBL | 1.3 | 0.7 | 3.1 | 1.1 | 1.8 | 5.7E-09 | 1.1E-07 |
| CBLB | 3.5 | 1.3 | 5.5 | 1.8 | 1.6 | 2.2E-05 | 1.7E-04 |
| CD247 | 1.0 | 0.9 | 6.9 | 12.6 | 8.8 | 2.1E-15 | 1.8E-13 |
| CD3D | 2.3 | 1.3 | 24.8 | 40.5 | 12.3 | 1.7E-16 | 2.0E-14 |
| CD3E | 2.3 | 1.5 | 16.4 | 34.3 | 9.4 | 1.1E-15 | 1.0E-13 |
| CD3G | 0.3 | 0.3 | 2.6 | 3.9 | 11.1 | 6.8E-22 | 1.2E-18 |
| CD4 | 6.6 | 2.1 | 20.8 | 27.5 | 4.3 | 1.4E-14 | 9.4E-13 |
| CD8A | 0.8 | 0.6 | 10.4 | 21.0 | 18.1 | 2.8E-20 | 1.6E-17 |
| CD8B | 1.1 | 1.1 | 4.8 | 9.5 | 5.0 | 1.3E-07 | 1.9E-06 |
| CTLA4 | 0.2 | 0.2 | 2.0 | 2.5 | 13.1 | 1.6E-12 | 6.4E-11 |
| GRAP2 | 0.3 | 0.4 | 2.0 | 3.4 | 7.9 | 9.1E-19 | 2.4E-16 |
| GRB2 | 24.1 | 6.0 | 28.3 | 9.3 | 1.2 | 1.7E-02 | 4.5E-02 |
| ITK | 0.0 | 0.0 | 1.5 | 5.3 | 9.3 | 2.0E-16 | 2.4E-14 |
| JUN | 7.4 | 4.2 | 16.4 | 3.6 | 1.9 | 1.5E-09 | 3.2E-08 |
| KRAS | 4.8 | 2.0 | 6.9 | 2.4 | 1.2 | 1.7E-02 | 4.5E-02 |
| LAT | 0.0 | 0.0 | 0.0 | 0.0 | 2.3 | 1.8E-05 | 1.5E-04 |
| LCK | 1.0 | 1.2 | 11.2 | 13.4 | 10.0 | 2.2E-19 | 7.5E-17 |
| LCP2 | 3.2 | 1.8 | 24.0 | 26.3 | 6.9 | 6.3E-20 | 3.2E-17 |
| MALT1 | 5.4 | 1.6 | 7.9 | 3.6 | 1.5 | 1.2E-05 | 1.0E-04 |
| MAP3K7 | 6.4 | 2.7 | 8.8 | 1.8 | 1.2 | 1.9E-02 | 4.8E-02 |
| MAPK13 | 6.1 | 1.3 | 9.3 | 4.3 | 1.5 | 7.1E-04 | 3.3E-03 |
| MAPK14 | 10.5 | 3.2 | 14.9 | 2.5 | 1.2 | 1.6E-02 | 4.3E-02 |
| NCK1 | 6.9 | 1.3 | 11.3 | 6.1 | 1.4 | 7.4E-05 | 4.9E-04 |
| NFATC1 | 2.2 | 0.7 | 3.5 | 2.2 | 1.6 | 1.3E-04 | 7.9E-04 |
| NFATC2 | 0.5 | 0.8 | 2.5 | 2.0 | 3.7 | 2.7E-13 | 1.3E-11 |
| NRAS | 7.3 | 1.2 | 10.9 | 2.6 | 1.3 | 6.1E-04 | 2.9E-03 |
| PAK2 | 7.0 | 3.2 | 12.5 | 4.4 | 1.5 | 1.7E-06 | 1.8E-05 |
| PAK3 | 0.3 | 0.4 | 0.3 | 0.3 | 1.8 | 7.2E-04 | 3.3E-03 |
| PDCD1 | 0.3 | 0.2 | 2.0 | 2.1 | 15.3 | 2.1E-14 | 1.3E-12 |
| PLCG1 | 11.5 | 3.2 | 18.8 | 4.6 | 1.3 | 9.9E-04 | 4.3E-03 |
| PPP3R1 | 6.9 | 1.0 | 9.8 | 1.6 | 1.2 | 1.1E-02 | 3.0E-02 |
| PTPN22 | 0.0 | 0.0 | 2.4 | 1.8 | 9.4 | 1.8E-20 | 1.2E-17 |
| PTPN6 | 12.9 | 5.7 | 26.6 | 32.5 | 2.6 | 2.3E-10 | 6.0E-09 |
| PTPRC | 3.0 | 1.8 | 25.3 | 28.1 | 8.8 | 1.4E-21 | 1.8E-18 |
| RASGRP1 | 2.2 | 1.8 | 3.9 | 2.4 | 1.6 | 4.6E-04 | 2.3E-03 |
| SOS1 | 3.8 | 2.0 | 5.6 | 1.4 | 1.3 | 7.0E-03 | 2.2E-02 |
| TRAC | 6.7 | 2.3 | 39.8 | 77.2 | 6.5 | 5.1E-15 | 3.8E-13 |
| TRBC2 | 8.3 | 7.6 | 80.7 | 151.5 | 9.7 | 1.1E-16 | 1.4E-14 |
| TRDC | 1.1 | 0.8 | 5.4 | 6.2 | 4.7 | 4.6E-08 | 7.3E-07 |
| TRGC2 | 0.7 | 0.6 | 4.5 | 4.1 | 8.1 | 5.3E-11 | 1.6E-09 |
| TRGV9 | 0.0 | 0.0 | 0.3 | 0.3 | 8.1 | 9.6E-07 | 1.1E-05 |
| VAV1 | 1.1 | 0.8 | 5.4 | 6.7 | 6.5 | 5.3E-17 | 7.9E-15 |
| ZAP70 | 2.0 | 1.5 | 11.0 | 19.8 | 7.3 | 7.8E-16 | 7.7E-14 |

Table S3A continued. T Cell Receptor Activation (Signal 1)

| Symbol | Normal Allograft Median | Normal Allograft IQR | TCMR Median | TCMR IQR | FC | PValue | FDR |
|---|---|---|---|---|---|---|---|
| HRAS | 26.8 | 5.5 | 25.6 | 7.1 | -1.3 | 1.6E-02 | 4.3E-02 |
| PAK4 | 18.7 | 4.7 | 16.5 | 5.8 | -1.3 | 2.6E-04 | 1.4E-03 |
| PAK5 | 0.1 | 0.1 | 0.1 | 0.0 | -2.2 | 1.2E-02 | 3.3E-02 |
| BCL10 | 3.8 | 0.7 | 5.0 | 1.2 | 1.1 | 9.9E-02 | NS |
| CBLC | 3.1 | 1.1 | 3.3 | 1.6 | -1.1 | 4.3E-01 | NS |
| CDC42 | 75.3 | 8.2 | 89.9 | 11.7 | 1.0 | 7.6E-01 | NS |
| DLG1 | 10.4 | 4.4 | 12.8 | 4.4 | 1.0 | 8.6E-01 | NS |
| FYN | 36.6 | 10.4 | 35.7 | 21.4 | 1.2 | 1.1E-01 | NS |
| MAP2K1 | 19.3 | 10.1 | 29.3 | 9.0 | 1.0 | 4.3E-01 | NS |
| MAP2K2 | 41.0 | 14.1 | 43.5 | 12.7 | -1.1 | 3.5E-01 | NS |
| MAP2K7 | 9.2 | 2.9 | 12.2 | 1.9 | 1.0 | 6.7E-01 | NS |
| MAPK1 | 7.7 | 3.2 | 10.3 | 2.7 | 1.1 | 2.6E-01 | NS |
| MAPK11 | 6.4 | 1.1 | 8.0 | 2.8 | 1.1 | 5.4E-01 | NS |
| MAPK12 | 0.0 | 14.6 | 0.0 | 0.0 | -1.2 | 2.7E-01 | NS |
| MAPK3 | 34.1 | 13.3 | 37.8 | 14.8 | -1.2 | 3.6E-02 | NS |
| MAPK9 | 7.3 | 2.1 | 9.3 | 5.5 | 1.0 | 8.5E-01 | NS |
| NCK2 | 11.9 | 3.1 | 14.9 | 3.2 | 1.1 | 1.5E-01 | NS |
| NFATC3 | 0.0 | 0.0 | 0.0 | 0.0 | 1.1 | 9.3E-02 | NS |
| PAK1 | 11.9 | 2.5 | 18.7 | 18.5 | 1.2 | 3.1E-02 | NS |
| PAK6 | 3.0 | 0.8 | 2.3 | 1.6 | -1.4 | 2.1E-02 | NS |
| PPP3CA | 4.4 | 1.4 | 6.0 | 1.1 | 1.1 | 7.7E-02 | NS |
| PPP3CB | 16.3 | 3.7 | 20.1 | 3.7 | 1.0 | 5.1E-01 | NS |
| PPP3CC | 4.2 | 0.9 | 6.5 | 1.8 | 1.2 | 9.3E-02 | NS |
| PRKCQ | 1.9 | 5.4 | 5.0 | 3.4 | 1.3 | 2.0E-01 | NS |
| RAF1 | 23.6 | 5.6 | 26.2 | 4.1 | -1.1 | 1.8E-01 | NS |
| RHOA | 130.0 | 25.0 | 197.5 | 61.9 | 1.2 | 1.4E-01 | NS |
| SOS2 | 7.1 | 1.5 | 8.0 | 2.5 | 1.0 | 8.8E-01 | NS |
| TEC | 0.8 | 0.2 | 1.0 | 0.3 | 1.2 | 2.5E-01 | NS |
| VAV2 | 13.2 | 2.2 | 13.0 | 2.9 | -1.1 | 2.8E-01 | NS |
| VAV3 | 23.5 | 29.8 | 34.8 | 30.7 | -1.3 | 7.7E-02 | NS |

Table S3B. Co-Stimulation (Signal 2)

| Symbol | Normal Allograft Median | Normal Allograft IQR | TCMR Median | TCMR IQR | FC | PValue | FDR |
|---|---|---|---|---|---|---|---|
| CARD11 | 0.5 | 0.3 | 3.0 | 4.3 | 6.2 | 2.7E-18 | 6.2E-16 |
| CD28 | 0.4 | 0.3 | 1.9 | 1.4 | 5.5 | 6.4E-14 | 3.5E-12 |
| CD40LG | 0.4 | 0.4 | 1.8 | 1.1 | 2.7 | 5.3E-07 | 6.6E-06 |
| FOS | 2.2 | 1.9 | 6.0 | 3.7 | 1.7 | 1.4E-02 | 3.8E-02 |
| ICOS | 0.1 | 0.1 | 1.6 | 2.0 | 15.2 | 1.1E-15 | 9.8E-14 |
| IKBKB | 9.5 | 4.4 | 16.7 | 4.0 | 1.5 | 2.9E-06 | 3.0E-05 |
| MAP3K14 | 6.1 | 2.1 | 8.9 | 3.9 | 1.6 | 6.1E-07 | 7.5E-06 |
| MAP3K8 | 2.9 | 0.7 | 5.9 | 4.8 | 2.4 | 1.8E-10 | 4.7E-09 |
| NFKB1 | 8.3 | 3.7 | 15.3 | 4.6 | 1.5 | 7.5E-07 | 9.0E-06 |
| NFKBIA | 27.7 | 11.1 | 54.8 | 19.9 | 1.5 | 4.2E-05 | 3.0E-04 |
| NFKBIE | 4.1 | 1.3 | 13.5 | 10.8 | 2.1 | 4.8E-08 | 7.7E-07 |
| PIK3CD | 0.0 | 0.0 | 8.0 | 6.5 | 2.0 | 1.3E-07 | 2.0E-06 |
| PIK3CG | 0.3 | 0.4 | 1.6 | 1.2 | 4.4 | 1.2E-10 | 3.2E-09 |
| PIK3R5 | 0.7 | 0.4 | 3.3 | 4.1 | 7.2 | 3.9E-20 | 2.1E-17 |
| AKT1 | 33.7 | 5.5 | 43.9 | 12.4 | 1.1 | 2.3E-01 | NS |
| AKT2 | 35.1 | 15.7 | 39.0 | 12.1 | -1.0 | 6.7E-01 | NS |
| AKT3 | 4.0 | 2.9 | 6.2 | 3.5 | 1.2 | 9.1E-02 | NS |
| CHUK | 8.6 | 1.1 | 7.8 | 2.1 | -1.1 | 2.9E-01 | NS |
| GSK3B | 4.2 | 1.4 | 5.5 | 1.7 | 1.2 | 2.5E-02 | NS |
| IKBKG | 7.3 | 2.9 | 9.1 | 4.5 | 1.0 | 6.7E-01 | NS |
| NFKBIB | 12.0 | 5.0 | 13.5 | 5.4 | -1.2 | 8.7E-02 | NS |
| PDPK1 | 7.8 | 2.4 | 9.3 | 2.0 | 1.0 | 5.9E-01 | NS |
| PIK3CA | 2.2 | 1.2 | 3.2 | 0.9 | 1.2 | 2.6E-02 | NS |
| PIK3CB | 10.7 | 2.4 | 10.8 | 2.7 | -1.1 | 2.2E-01 | NS |
| PIK3R1 | 9.1 | 3.0 | 11.4 | 4.1 | 1.1 | 4.4E-01 | NS |
| PIK3R3 | 5.7 | 1.5 | 7.1 | 3.3 | 1.1 | 4.1E-01 | NS |
| RELA | 18.1 | 3.1 | 23.1 | 6.8 | 1.1 | 3.0E-01 | NS |

Supplemental Table S4. Intragraft Expression of mRNAs Encoding B Cell Receptor Signaling KEGG Pathway Genes in TCMR and Normal kidney allograft biopsies.

Table S4A. B Cell Activation (Signal 1)

| | Normal Allograft | | TCMR | | | | |
|---|---|---|---|---|---|---|---|
| Symbol | Median | IQR | Median | IQR | FC | PValue | FDR |
| BLNK | 8.865 | 3.021 | 15.46 | 5.967 | 1.388 | 1.86E-03 | 7.30E-03 |
| BTK | 0.781 | 0.69 | 3.86 | 6.623 | 6.968 | 4.25E-18 | 9.11E-16 |
| CARD11 | 0.492 | 0.335 | 2.95 | 4.261 | 6.249 | 2.72E-18 | 6.19E-16 |
| CD79A | 0.468 | 0.334 | 5.06 | 13.669 | 13.734 | 2.40E-13 | 1.17E-11 |
| CD79B | 1.878 | 1.009 | 4.31 | 5.29 | 3.288 | 3.59E-09 | 7.33E-08 |
| DAPP1 | 0.427 | 0.259 | 3.74 | 5.315 | 8.949 | 4.23E-22 | 9.89E-19 |
| FOS | 2.155 | 1.853 | 5.97 | 3.727 | 1.666 | 1.39E-02 | 3.78E-02 |
| GRB2 | 24.143 | 6.006 | 28.35 | 9.296 | 1.24 | 1.71E-02 | 4.47E-02 |
| JUN | 7.354 | 4.246 | 16.43 | 3.551 | 1.896 | 1.47E-09 | 3.24E-08 |
| KRAS | 4.784 | 1.959 | 6.92 | 2.361 | 1.193 | 1.74E-02 | 4.53E-02 |
| LYN | 3.676 | 2.754 | 9.15 | 11.2 | 3.069 | 4.38E-12 | 1.61E-10 |
| MALT1 | 5.36 | 1.6 | 7.92 | 3.557 | 1.494 | 1.20E-05 | 1.02E-04 |
| NFATC1 | 2.201 | 0.681 | 3.46 | 2.182 | 1.631 | 1.33E-04 | 7.91E-04 |
| NFATC2 | 0.538 | 0.758 | 2.45 | 1.952 | 3.681 | 2.75E-13 | 1.31E-11 |
| NRAS | 7.272 | 1.189 | 10.89 | 2.562 | 1.291 | 6.10E-04 | 2.91E-03 |
| PIK3AP1 | 3.753 | 2.855 | 10.08 | 8.54 | 2.528 | 1.67E-09 | 3.63E-08 |
| PPP3R1 | 6.886 | 1.015 | 9.84 | 1.568 | 1.2 | 1.05E-02 | 3.00E-02 |
| PRKCB | 0.416 | 0.326 | 3.25 | 4.664 | 6.439 | 8.64E-12 | 2.99E-10 |
| RAC2 | 3.312 | 2.607 | 18.39 | 26.754 | 8.366 | 3.89E-17 | 6.07E-15 |
| RASGRP3 | 2.75 | 1.69 | 5.02 | 1.839 | 1.317 | 1.01E-02 | 2.89E-02 |
| SOS1 | 3.808 | 2.042 | 5.6 | 1.397 | 1.252 | 7.01E-03 | 2.17E-02 |
| SYK | 4.968 | 2.415 | 12.05 | 8.086 | 1.529 | 3.23E-03 | 1.15E-02 |
| VAV1 | 1.065 | 0.761 | 5.43 | 6.665 | 6.47 | 5.31E-17 | 7.91E-15 |

Table S4A continued. B Cell Activation (Signal 1)

| Symbol | Normal Allograft Median | Normal Allograft IQR | TCMR Median | TCMR IQR | FC | PValue | FDR |
|---|---|---|---|---|---|---|---|
| HRAS | 26.779 | 5.523 | 25.59 | 7.057 | -1.314 | 1.64E-02 | 4.31E-02 |
| RAC3 | 6.306 | 4.065 | 4.86 | 2.435 | -1.602 | 7.42E-03 | 2.27E-02 |
| BCL10 | 3.844 | 0.687 | 4.99 | 1.17 | 1.141 | 9.85E-02 | 1.81E-01 |
| MAP2K1 | 19.309 | 10.149 | 29.34 | 8.969 | 1.049 | 4.29E-01 | 5.57E-01 |
| MAP2K2 | 41.022 | 14.126 | 43.51 | 12.726 | -1.086 | 3.50E-01 | 4.79E-01 |
| MAPK1 | 7.705 | 3.19 | 10.31 | 2.733 | 1.093 | 2.57E-01 | 3.80E-01 |
| MAPK3 | 34.096 | 13.34 | 37.81 | 14.838 | -1.178 | 3.64E-02 | 8.23E-02 |
| NFATC3 | 0 | 0 | 0 | 0 | 1.124 | 9.32E-02 | 1.74E-01 |
| PLCG2 | 14.634 | 3.942 | 14 | 4.325 | -1.241 | 6.95E-02 | 1.38E-01 |
| PPP3CA | 4.407 | 1.421 | 6 | 1.074 | 1.137 | 7.75E-02 | 1.50E-01 |
| PPP3CB | 16.272 | 3.734 | 20.07 | 3.693 | 1.039 | 5.08E-01 | 6.32E-01 |
| PPP3CC | 4.213 | 0.865 | 6.47 | 1.759 | 1.164 | 9.26E-02 | 1.73E-01 |
| RAC1 | 66.655 | 5.07 | 77.72 | 18.315 | 1.027 | 7.28E-01 | 8.16E-01 |
| RAF1 | 23.56 | 5.576 | 26.23 | 4.101 | -1.095 | 1.76E-01 | 2.84E-01 |
| RELA | 18.062 | 3.128 | 23.09 | 6.845 | 1.081 | 3.01E-01 | 4.28E-01 |
| SOS2 | 7.119 | 1.507 | 8.03 | 2.511 | 1.011 | 8.80E-01 | 9.23E-01 |
| VAV2 | 13.178 | 2.167 | 13 | 2.901 | -1.076 | 2.80E-01 | 4.06E-01 |
| VAV3 | 23.548 | 29.806 | 34.78 | 30.674 | -1.259 | 7.72E-02 | 1.49E-01 |

Table S4B. B Cell Activation (Co-Stimulator)

| Symbol | Normal Allograft Median | Normal Allograft IQR | TCMR Median | TCMR IQR | FC | PValue | FDR |
|---|---|---|---|---|---|---|---|
| CD19 | 0 | 0 | 0 | 0 | 9.289 | 1.21E-11 | 4.05E-10 |
| IFITM1 | 63.024 | 34.624 | 145.02 | 55.393 | 2.204 | 1.29E-08 | 2.37E-07 |
| IKBKB | 9.515 | 4.376 | 16.74 | 4.046 | 1.46 | 2.90E-06 | 2.96E-05 |
| NFKB1 | 8.332 | 3.719 | 15.28 | 4.597 | 1.521 | 7.55E-07 | 8.99E-06 |
| NFKBIA | 27.722 | 11.116 | 54.82 | 19.852 | 1.511 | 4.22E-05 | 3.01E-04 |
| NFKBIE | 4.096 | 1.272 | 13.47 | 10.8 | 2.111 | 4.83E-08 | 7.68E-07 |
| PIK3CD | 0 | 0 | 8.01 | 6.482 | 1.969 | 1.35E-07 | 1.96E-06 |
| AKT1 | 33.658 | 5.525 | 43.9 | 12.404 | 1.093 | 2.33E-01 | 3.53E-01 |
| AKT2 | 35.086 | 15.667 | 39.04 | 12.126 | -1.039 | 6.65E-01 | 7.67E-01 |
| AKT3 | 4.002 | 2.904 | 6.25 | 3.511 | 1.21 | 9.07E-02 | 1.70E-01 |
| CD81 | 240.129 | 55.515 | 300.89 | 63.075 | -1.016 | 8.80E-01 | 9.23E-01 |
| CHUK | 8.56 | 1.134 | 7.82 | 2.088 | -1.083 | 2.87E-01 | 4.13E-01 |
| CR2 | 2.912 | 6.748 | 0.41 | 1.154 | -2.407 | 4.63E-02 | 9.99E-02 |
| GSK3B | 4.178 | 1.408 | 5.5 | 1.676 | 1.182 | 2.53E-02 | 6.14E-02 |
| IKBKG | 7.333 | 2.932 | 9.12 | 4.534 | 1.041 | 6.75E-01 | 7.74E-01 |
| NFKBIB | 12.012 | 5.041 | 13.51 | 5.369 | -1.201 | 8.73E-02 | 1.65E-01 |
| PIK3CA | 2.16 | 1.208 | 3.23 | 0.925 | 1.2 | 2.64E-02 | 6.34E-02 |
| PIK3CB | 10.714 | 2.388 | 10.77 | 2.678 | -1.1 | 2.19E-01 | 3.36E-01 |
| PIK3R1 | 9.074 | 2.971 | 11.42 | 4.053 | 1.072 | 4.36E-01 | 5.65E-01 |

Table S4C. B Cell Activation (Co-Inhibitor)

| Symbol | Normal Allograft Median | Normal Allograft IQR | TCMR Median | TCMR IQR | FC | PValue | FDR |
|---|---|---|---|---|---|---|---|
| CD22 | 0.947 | 0.696 | 1.86 | 1.391 | 2.459 | 4.77E-04 | 2.36E-03 |
| CD72 | 4.775 | 1.149 | 7.51 | 8.711 | 2.418 | 1.48E-06 | 1.62E-05 |
| FCGR2B | 0 | 0 | 0 | 4.942 | 5.396 | 1.08E-10 | 2.99E-09 |
| INPP5D | 2.835 | 1.844 | 10.32 | 10.59 | 3.764 | 2.16E-14 | 1.36E-12 |
| LILRB3 | 0.462 | 0.514 | 3.78 | 2.875 | 4.853 | 1.67E-08 | 2.99E-07 |
| PTPN6 | 12.925 | 5.663 | 26.62 | 32.454 | 2.569 | 2.31E-10 | 5.97E-09 |
| INPPL1 | 26.832 | 3.895 | 31.57 | 8.032 | 1.009 | 9.07E-01 | 9.41E-01 |

Supplemental Table S5. Intragraft Expression of mRNAs Encoding of Leukocyte Transendothelial Migration KEGG Pathway Genes in TCMR and Normal Kidney Allograft Biopsies.

Table S5A. Leukocyte Migration Machinery Genes

| | Normal Allograft | | TCMR | | | | |
|---|---|---|---|---|---|---|---|
| Symbol | Median | IQR | Median | IQR | FC | PValue | FDR |
| PIK3R5 | 0.7 | 0.4 | 3.3 | 4.1 | 7.2 | 3.9E-20 | 2.1E-17 |
| ITGAL | 1.3 | 1.0 | 11.8 | 20.1 | 8.8 | 3.0E-18 | 6.7E-16 |
| ITGB2 | 7.6 | 5.0 | 32.3 | 61.5 | 6.6 | 6.1E-18 | 1.2E-15 |
| RAC2 | 3.3 | 2.6 | 18.4 | 26.8 | 8.4 | 3.9E-17 | 6.1E-15 |
| VAV1 | 1.1 | 0.8 | 5.4 | 6.7 | 6.5 | 5.3E-17 | 7.9E-15 |
| ITK | 0.0 | 0.0 | 1.5 | 5.3 | 9.3 | 2.0E-16 | 2.4E-14 |
| RHOH | 0.5 | 0.4 | 3.6 | 5.2 | 9.6 | 2.0E-16 | 2.4E-14 |
| RASSF5 | 1.6 | 1.0 | 7.4 | 7.8 | 5.2 | 4.0E-16 | 4.3E-14 |
| ITGAM | 1.2 | 0.7 | 4.2 | 4.2 | 3.6 | 2.7E-14 | 1.6E-12 |
| ITGA4 | 0.7 | 1.0 | 6.1 | 5.4 | 4.6 | 5.1E-14 | 2.9E-12 |
| ICAM1 | 5.3 | 2.5 | 23.3 | 22.5 | 3.1 | 7.3E-13 | 3.1E-11 |
| PRKCB | 0.4 | 0.3 | 3.3 | 4.7 | 6.4 | 8.6E-12 | 3.0E-10 |
| PIK3CG | 0.3 | 0.4 | 1.6 | 1.2 | 4.4 | 1.2E-10 | 3.2E-09 |
| MYLPF | 0.5 | 0.3 | 1.1 | 0.8 | 16.4 | 8.5E-08 | 1.3E-06 |
| PIK3CD | 0.0 | 0.0 | 8.0 | 6.5 | 2.0 | 1.3E-07 | 2.0E-06 |
| MYL2 | 0.1 | 0.2 | 0.2 | 0.5 | 19.3 | 3.2E-05 | 2.4E-04 |
| GNAI2 | 57.1 | 10.4 | 81.9 | 50.2 | 1.5 | 8.4E-04 | 3.8E-03 |
| MYL9 | 49.0 | 25.2 | 45.1 | 10.3 | -1.5 | 9.9E-04 | 4.3E-03 |
| GNAI3 | 12.9 | 2.6 | 18.0 | 3.3 | 1.2 | 1.4E-03 | 5.8E-03 |
| PTK2B | 10.6 | 2.0 | 15.3 | 7.9 | 1.4 | 1.8E-03 | 7.1E-03 |
| ITGB1 | 64.6 | 40.0 | 110.6 | 41.8 | 1.4 | 4.3E-03 | 1.5E-02 |
| ROCK1 | 3.5 | 2.6 | 5.8 | 2.4 | 1.3 | 4.7E-03 | 1.6E-02 |
| RAP1B | 31.6 | 5.7 | 43.6 | 11.4 | 1.2 | 1.5E-02 | 4.0E-02 |
| RAP1A | 27.1 | 3.5 | 46.8 | 16.7 | 1.2 | 2.5E-02 | NS |
| PIK3CA | 2.2 | 1.2 | 3.2 | 0.9 | 1.2 | 2.6E-02 | NS |
| MYL12A | 161.3 | 59.2 | 324.5 | 137.8 | 1.2 | 2.9E-02 | NS |
| RAPGEF4 | 1.9 | 0.7 | 1.8 | 0.7 | -1.2 | 5.6E-02 | NS |
| VAV3 | 23.5 | 29.8 | 34.8 | 30.7 | -1.3 | 7.7E-02 | NS |
| RHOA | 130.0 | 25.0 | 197.5 | 61.9 | 1.2 | 1.4E-01 | NS |
| PIK3CB | 10.7 | 2.4 | 10.8 | 2.7 | -1.1 | 2.2E-01 | NS |
| VAV2 | 13.2 | 2.2 | 13.0 | 2.9 | -1.1 | 2.8E-01 | NS |
| MYL12B | 214.9 | 37.8 | 262.7 | 41.0 | -1.1 | 3.4E-01 | NS |
| PIK3R3 | 5.7 | 1.5 | 7.1 | 3.3 | 1.1 | 4.1E-01 | NS |
| ROCK2 | 4.6 | 2.6 | 4.7 | 2.5 | 1.1 | 4.2E-01 | NS |
| PIK3R1 | 9.1 | 3.0 | 11.4 | 4.1 | 1.1 | 4.4E-01 | NS |
| GNAI1 | 8.9 | 19.9 | 15.7 | 21.7 | 1.1 | 6.6E-01 | NS |
| RAPGEF3 | 35.2 | 24.9 | 35.4 | 20.6 | 1.0 | 7.5E-01 | NS |
| CDC42 | 75.3 | 8.2 | 89.9 | 11.7 | 1.0 | 7.6E-01 | NS |

Table S5B. Genes of Endothelial Cell Changes to Allow Migration

| Symbol | Normal Allograft Median | Normal Allograft IQR | TCMR Median | TCMR IQR | FC | PValue | FDR |
|---|---|---|---|---|---|---|---|
| CYBB | 3.8 | 3.0 | 20.4 | 29.1 | 6.4 | 1.5E-18 | 3.8E-16 |
| NCF2 | 1.6 | 1.0 | 7.7 | 15.3 | 5.9 | 3.8E-16 | 4.1E-14 |
| NCF4 | 2.7 | 0.8 | 10.5 | 10.0 | 4.0 | 1.1E-13 | 5.7E-12 |
| CXCR4 | 3.4 | 2.2 | 13.7 | 17.2 | 4.0 | 1.5E-12 | 6.3E-11 |
| NCF1 | 1.6 | 0.7 | 7.2 | 8.6 | 5.4 | 2.0E-12 | 7.9E-11 |
| CLDN1 | 2.8 | 1.8 | 8.3 | 5.3 | 2.9 | 2.9E-12 | 1.1E-10 |
| SIPA1 | 13.1 | 7.7 | 13.4 | 14.6 | 2.0 | 1.4E-09 | 3.2E-08 |
| MMP9 | 0.8 | 0.7 | 7.1 | 9.9 | 7.7 | 1.8E-09 | 3.9E-08 |
| VASP | 12.2 | 3.3 | 25.6 | 12.0 | 1.7 | 5.1E-08 | 8.0E-07 |
| VCAM1 | 24.8 | 10.1 | 77.5 | 63.0 | 2.5 | 2.4E-07 | 3.3E-06 |
| TXK | 0.7 | 0.4 | 1.6 | 1.0 | 2.1 | 2.4E-05 | 1.8E-04 |
| ACTN2 | 0.5 | 0.3 | 0.4 | 0.2 | 4.7 | 5.5E-05 | 3.8E-04 |
| CLDN3 | 13.4 | 8.4 | 23.7 | 14.2 | 1.7 | 6.1E-05 | 4.1E-04 |
| CLDN4 | 23.5 | 10.6 | 48.4 | 33.1 | 1.6 | 1.9E-04 | 1.1E-03 |
| MAPK13 | 6.1 | 1.3 | 9.3 | 4.3 | 1.5 | 7.1E-04 | 3.3E-03 |
| ACTN1 | 14.9 | 7.0 | 31.8 | 13.2 | 1.5 | 8.6E-04 | 3.9E-03 |
| PLCG1 | 11.5 | 3.2 | 18.8 | 4.6 | 1.3 | 9.9E-04 | 4.3E-03 |
| MSN | 75.5 | 26.6 | 110.0 | 38.5 | 1.4 | 5.0E-03 | 1.6E-02 |
| MAPK14 | 10.5 | 3.2 | 14.9 | 2.5 | 1.2 | 1.6E-02 | 4.3E-02 |
| THY1 | 257.4 | 121.5 | 122.3 | 119.6 | -2.2 | 3.9E-05 | 2.8E-04 |
| CLDN22 | 1.6 | 2.2 | 0.0 | 0.0 | -1.5 | 1.1E-03 | 4.6E-03 |
| CLDN8 | 29.2 | 12.8 | 17.6 | 16.1 | -1.7 | 5.4E-03 | 1.7E-02 |
| CTNNA3 | 0.1 | 0.1 | 0.0 | 0.1 | -1.9 | 6.1E-03 | 1.9E-02 |
| CLDN10 | 88.4 | 27.5 | 81.5 | 73.9 | -1.4 | 1.9E-02 | 4.9E-02 |
| CLDN9 | 0.1 | 0.2 | 0.4 | 0.1 | 1.7 | 3.8E-02 | NS |
| PTPN11 | 13.7 | 6.8 | 19.3 | 4.9 | 1.2 | 3.8E-02 | NS |
| ACTB | 613.1 | 123.5 | 764.1 | 477.7 | 1.3 | 6.4E-02 | NS |
| PLCG2 | 14.6 | 3.9 | 14.0 | 4.3 | -1.2 | 6.9E-02 | NS |
| CTNNA1 | 97.4 | 21.7 | 95.4 | 12.2 | -1.2 | 7.8E-02 | NS |
| CD99 | 48.0 | 15.9 | 59.0 | 54.6 | 1.2 | 9.4E-02 | NS |
| ACTN4 | 171.1 | 48.1 | 183.9 | 54.6 | -1.2 | 1.2E-01 | NS |
| ACTG1 | 514.6 | 132.9 | 715.7 | 314.6 | 1.2 | 1.2E-01 | NS |
| CLDN2 | 116.0 | 42.3 | 77.6 | 67.0 | -1.5 | 1.4E-01 | NS |
| CLDN16 | 19.1 | 18.9 | 9.2 | 19.2 | -1.4 | 1.7E-01 | NS |
| MMP2 | 16.9 | 7.3 | 24.6 | 20.5 | 1.2 | 1.9E-01 | NS |
| ARHGAP35 | 11.4 | 3.4 | 11.6 | 3.4 | -1.1 | 1.9E-01 | NS |
| CLDN18 | 0.1 | 0.1 | 0.2 | 0.1 | -1.3 | 2.1E-01 | NS |
| PRKCA | 6.9 | 2.2 | 6.7 | 2.5 | -1.1 | 2.2E-01 | NS |
| CLDN5 | 5.6 | 3.5 | 7.1 | 4.3 | 1.2 | 2.4E-01 | NS |
| CLDN11 | 2.4 | 1.6 | 2.7 | 2.2 | 1.3 | 2.4E-01 | NS |
| VCL | 18.1 | 10.6 | 24.2 | 11.4 | 1.1 | 2.7E-01 | NS |
| CLDN23 | 1.0 | 1.5 | 2.1 | 0.7 | 1.3 | 2.8E-01 | NS |
| CTNNB1 | 52.8 | 19.5 | 67.7 | 11.3 | 1.1 | 2.8E-01 | NS |
| CXCL12 | 96.5 | 64.5 | 110.6 | 64.4 | -1.2 | 2.9E-01 | NS |
| CYBA | 198.0 | 76.8 | 279.4 | 258.0 | 1.2 | 3.1E-01 | NS |
| BCAR1 | 26.5 | 5.6 | 30.3 | 9.6 | -1.1 | 5.2E-01 | NS |
| CDH5 | 7.7 | 6.9 | 10.7 | 5.0 | 1.1 | 5.3E-01 | NS |
| PXN | 32.4 | 4.8 | 33.8 | 4.7 | -1.1 | 5.3E-01 | NS |
| MAPK11 | 6.4 | 1.1 | 8.0 | 2.8 | 1.1 | 5.4E-01 | NS |
| MLLT4 | 10.4 | 5.1 | 13.1 | 6.9 | 1.1 | 5.8E-01 | NS |
| CLDN19 | 12.2 | 37.3 | 14.4 | 30.6 | -1.2 | 5.9E-01 | NS |
| F11R | 20.7 | 8.1 | 23.9 | 10.0 | -1.0 | 5.9E-01 | NS |
| CLDN14 | 2.7 | 3.4 | 3.4 | 5.4 | 1.2 | 6.0E-01 | NS |
| PTK2 | 19.7 | 8.1 | 23.2 | 7.4 | -1.0 | 6.0E-01 | NS |
| JAM3 | 3.5 | 3.5 | 4.2 | 5.1 | -1.1 | 6.1E-01 | NS |
| ESAM | 24.4 | 24.0 | 25.4 | 27.9 | -1.1 | 6.2E-01 | NS |
| CTNND1 | 44.4 | 17.3 | 51.9 | 19.4 | 1.0 | 6.8E-01 | NS |
| ARHGAP5 | 12.1 | 6.0 | 14.8 | 8.1 | 1.0 | 7.1E-01 | NS |
| RAC1 | 66.7 | 5.1 | 77.7 | 18.3 | 1.0 | 7.3E-01 | NS |
| OCLN | 4.8 | 7.3 | 7.0 | 6.1 | -1.0 | 7.7E-01 | NS |
| EZR | 100.7 | 25.0 | 128.2 | 35.9 | -1.0 | 9.3E-01 | NS |
| CLDN15 | 8.6 | 3.4 | 11.0 | 6.5 | -1.0 | 1.0E+00 | NS |

Supplemental Table S6. Intragraft Expression of mRNAs Encoding Cell Cycle KEGG Pathway Genes in TCMR and Normal Kidney Allograft Biopsies.

Table S6A. Cyclins, CDKs, and Cycle Inhibitor Complex

| | Normal Allograft | | TCMR | | | | |
|---|---|---|---|---|---|---|---|
| Symbol | Median | IQR | Median | IQR | FC | PValue | FDR |
| ***Cyclins & CDKs*** | | | | | | | |
| CCNA2 | 1.059 | 0.404 | 2.23 | 1.903 | 1.945 | 1.00E-05 | 8.76E-05 |
| CCNB1 | 3.393 | 0.868 | 5.36 | 3.547 | 1.64 | 1.16E-03 | 4.90E-03 |
| CCNB2 | 0.461 | 0.34 | 1.33 | 1.236 | 3.117 | 2.95E-06 | 3.01E-05 |
| CCND2 | 1.561 | 1.017 | 5.08 | 2.698 | 2.689 | 9.99E-10 | 2.30E-08 |
| CCND3 | 17.599 | 4.399 | 26.6 | 11.116 | 1.338 | 2.82E-03 | 1.03E-02 |
| CCNE1 | 0.663 | 0.232 | 1.13 | 0.728 | 1.821 | 1.70E-03 | 6.79E-03 |
| CCNE2 | 0 | 0.616 | 1.12 | 1.872 | 1.43 | 2.44E-03 | 9.12E-03 |
| CDC7 | 0.911 | 0.623 | 2.43 | 2.472 | 2.405 | 3.51E-07 | 4.59E-06 |
| CDK1 | 1.779 | 0.943 | 2.81 | 4.143 | 1.828 | 6.76E-04 | 3.16E-03 |
| CDK2 | 4.418 | 2.53 | 9.79 | 11.008 | 1.438 | 1.64E-04 | 9.50E-04 |
| CDK4 | 34.052 | 13.757 | 32.44 | 28.958 | -1.303 | 2.60E-03 | 9.60E-03 |
| CCNA1 | 0.124 | 0.082 | 0.2 | 0.185 | 1.025 | 9.36E-01 | 9.59E-01 |
| CCNB3 | 0.198 | 0.108 | 0.23 | 0.188 | -1.019 | 9.30E-01 | 9.55E-01 |
| CCND1 | 26.944 | 8.851 | 24.86 | 8.517 | -1.196 | 6.43E-02 | 1.30E-01 |
| CCNH | 14.011 | 8.008 | 9.68 | 5.685 | -1.181 | 3.63E-02 | 8.22E-02 |
| CDK6 | 1.452 | 0.98 | 2.32 | 0.775 | 1.304 | 2.01E-02 | 5.08E-02 |
| CDK7 | 12.646 | 5.333 | 13.81 | 4.318 | -1.134 | 1.65E-01 | 2.70E-01 |
| ***Cycle Inhibitor Complex*** | | | | | | | |
| CDKN1A | 9.667 | 4.757 | 20.28 | 7.522 | 1.69 | 2.97E-04 | 1.58E-03 |
| CDKN2B | 0.251 | 0.57 | 0.74 | 0.533 | 2.524 | 5.05E-06 | 4.84E-05 |
| CUL1 | 11.545 | 2.322 | 16.62 | 5.189 | 1.248 | 1.82E-03 | 7.17E-03 |
| MYC | 2.28 | 1.486 | 5.71 | 6.206 | 2.834 | 2.77E-08 | 4.67E-07 |
| PKMYT1 | 0 | 0 | 0 | 0.066 | 2.72 | 7.10E-07 | 8.52E-06 |
| SFN | 0.372 | 0.47 | 1.9 | 1.13 | 5.582 | 2.26E-07 | 3.11E-06 |
| SKP2 | 2 | 0.801 | 2.89 | 0.671 | 1.237 | 1.69E-02 | 4.42E-02 |
| TGFB1 | 9.016 | 2.602 | 19.59 | 9.281 | 1.72 | 9.02E-07 | 1.05E-05 |
| TGFB3 | 2.677 | 1.818 | 4.36 | 1.559 | 1.508 | 1.23E-03 | 5.15E-03 |
| SKP1 | 317.706 | 30.841 | 365.42 | 85.54 | -1.332 | 2.45E-03 | 9.15E-03 |
| CDKN1B | 14.972 | 2.605 | 18.08 | 3.51 | 1.104 | 1.74E-01 | 2.81E-01 |
| CDKN1C | 23.308 | 18.281 | 20.6 | 10.821 | -1.412 | 5.26E-02 | 1.10E-01 |
| CDKN2C | 5.026 | 2.449 | 5.27 | 2.109 | 1.001 | 9.97E-01 | 9.98E-01 |
| CDKN2D | 3.839 | 1.773 | 5.38 | 2.411 | 1.169 | 2.40E-01 | 3.62E-01 |
| GADD45G | 6.081 | 4.342 | 6.56 | 2.254 | -1.382 | 4.39E-02 | 9.57E-02 |
| GSK3B | 4.178 | 1.408 | 5.5 | 1.676 | 1.182 | 2.53E-02 | 6.14E-02 |
| PCNA | 24.017 | 4.671 | 30.49 | 9.483 | 1.095 | 2.98E-01 | 4.25E-01 |
| SMAD2 | 10.786 | 3.46 | 17.8 | 8.891 | 1.171 | 2.55E-02 | 6.17E-02 |
| SMAD3 | 9.408 | 2.994 | 12.46 | 3.99 | 1.164 | 3.24E-02 | 7.49E-02 |
| SMAD4 | 7.644 | 4.554 | 9.38 | 3.873 | 1.078 | 2.80E-01 | 4.06E-01 |
| TGFB2 | 0.994 | 1.167 | 1.81 | 1.366 | 1.273 | 1.82E-01 | 2.91E-01 |
| ZBTB17 | 7.122 | 1.031 | 9.83 | 3.909 | 1.076 | 3.16E-01 | 4.44E-01 |

Table S6B. Transcription, Division Machinery and Damage Control Genes

| Symbol | Normal Allograft Median | Normal Allograft IQR | TCMR Median | TCMR IQR | FC | PValue | FDR |
|---|---|---|---|---|---|---|---|
| ***Transcription*** | | | | | | | |
| DBF4 | 0 | 0.98 | 1.96 | 3.399 | 1.575 | 3.12E-04 | 1.65E-03 |
| E2F1 | 0.583 | 0.209 | 0.94 | 0.899 | 2.106 | 2.45E-04 | 1.34E-03 |
| E2F2 | 0.104 | 0.045 | 0.25 | 0.32 | 4.261 | 2.18E-07 | 3.01E-06 |
| HDAC1 | 30.156 | 3.037 | 43.11 | 11.323 | 1.205 | 8.76E-03 | 2.59E-02 |
| RB1 | 5.719 | 3.109 | 9.66 | 4.853 | 1.562 | 5.84E-06 | 5.46E-05 |
| RBL1 | 0.636 | 0.495 | 1.28 | 0.692 | 1.575 | 4.40E-04 | 2.21E-03 |
| ABL1 | 10.947 | 3.091 | 12.18 | 3.639 | -1.012 | 8.83E-01 | 9.25E-01 |
| E2F3 | 2.847 | 1.048 | 4.31 | 1.962 | 1.233 | 5.60E-02 | 1.16E-01 |
| E2F4 | 22.253 | 2.521 | 25.33 | 4.722 | -1.033 | 6.46E-01 | 7.50E-01 |
| E2F5 | 1.665 | 0.814 | 2.04 | 0.746 | -1.12 | 1.26E-01 | 2.19E-01 |
| EP300 | 5.695 | 4.014 | 8.57 | 3.034 | 1.208 | 5.08E-02 | 1.07E-01 |
| HDAC2 | 22.529 | 1.58 | 27.2 | 4.368 | 1.049 | 4.19E-01 | 5.48E-01 |
| RBL2 | 14.711 | 8.307 | 22.39 | 7.19 | 1.147 | 1.00E-01 | 1.83E-01 |
| RBX1 | 29.984 | 9.094 | 31.85 | 6.573 | -1.011 | 9.16E-01 | 9.47E-01 |
| STAG2 | 8.026 | 6.141 | 10.25 | 5.377 | 1.153 | 1.45E-01 | 2.43E-01 |
| TFDP1 | 13.954 | 4.642 | 19.66 | 5.637 | 1.033 | 6.45E-01 | 7.49E-01 |
| TFDP2 | 4.348 | 2.249 | 5 | 2.216 | -1.069 | 4.29E-01 | 5.57E-01 |
| ***Division Machinary*** | | | | | | | |
| ANAPC4 | 8.219 | 3.296 | 12.61 | 2.983 | 1.234 | 1.86E-03 | 7.29E-03 |
| BUB1 | 0.282 | 0.104 | 0.63 | 1.146 | 4.867 | 4.83E-09 | 9.66E-08 |
| BUB1B | 0 | 0 | 0.8 | 1.302 | 3.321 | 2.62E-07 | 3.57E-06 |
| CDC25A | 0.36 | 0.152 | 0.62 | 0.409 | 1.506 | 7.40E-03 | 2.26E-02 |
| CDC25B | 7.98 | 3.999 | 16.07 | 10.782 | 1.912 | 6.49E-07 | 7.88E-06 |
| CDC27 | 4.409 | 1.653 | 6.79 | 2.108 | 1.296 | 2.32E-04 | 1.27E-03 |
| CDC45 | 0.377 | 0.205 | 0.64 | 1.64 | 4.24 | 1.52E-07 | 2.18E-06 |
| CDC6 | 0.328 | 0.343 | 0.64 | 0.787 | 2.658 | 2.01E-05 | 1.60E-04 |
| MCM2 | 2.737 | 1.1 | 3.62 | 1.641 | 1.485 | 1.55E-03 | 6.25E-03 |
| MCM5 | 6.257 | 1.326 | 9.97 | 4.358 | 1.55 | 1.12E-04 | 6.85E-04 |
| MCM6 | 2.777 | 0.416 | 5.67 | 4.446 | 1.715 | 1.19E-05 | 1.01E-04 |
| MDM2 | 4.238 | 2.078 | 7.06 | 4.516 | 1.364 | 2.19E-05 | 1.72E-04 |
| PLK1 | 1.087 | 0.216 | 1.45 | 0.876 | 1.784 | 5.75E-04 | 2.76E-03 |
| PTTG1 | 1.761 | 0.659 | 4.41 | 7.019 | 3.56 | 1.64E-07 | 2.33E-06 |
| SMC3 | 6.805 | 4.538 | 10.86 | 3.858 | 1.26 | 7.82E-03 | 2.37E-02 |
| TTK | 0.208 | 0.094 | 0.45 | 1.044 | 3.181 | 1.61E-06 | 1.75E-05 |
| YWHAH | 24.776 | 5.697 | 48.68 | 21.429 | 1.576 | 3.05E-07 | 4.08E-06 |
| YWHAZ | 68.065 | 12.6 | 105.98 | 30.099 | 1.274 | 4.84E-03 | 1.60E-02 |
| ANAPC11* | 0 | 0 | 0 | 0 | -1.484 | 3.76E-04 | 1.93E-03 |
| ANAPC13 | 55.023 | 16.538 | 47.07 | 7.612 | -1.35 | 1.32E-04 | 7.89E-04 |
| CDC14B | 8.538 | 1.446 | 9.34 | 2.117 | -1.254 | 3.76E-03 | 1.30E-02 |
| YWHAE | 169.641 | 29.179 | 171.65 | 19.99 | -1.244 | 1.51E-02 | 4.03E-02 |
| ANAPC2 | 19.025 | 2.292 | 20.42 | 3.245 | -1.108 | 1.94E-01 | 3.06E-01 |
| ANAPC5 | 46.269 | 6.909 | 48.42 | 7.054 | -1.045 | 5.06E-01 | 6.30E-01 |
| ANAPC7** | 0 | 0 | 0 | 0 | 1.101 | 1.50E-01 | 2.51E-01 |
| BUB3 | 22.013 | 6.049 | 30.97 | 8.092 | 1.154 | 3.24E-02 | 7.49E-02 |
| CDC14A | 2.358 | 0.83 | 2.25 | 0.587 | -1.12 | 3.07E-01 | 4.34E-01 |
| CDC16 | 22.407 | 3.397 | 26.45 | 4.086 | 1.019 | 7.60E-01 | 8.38E-01 |
| CDC20 | 1.63 | 1.375 | 2.01 | 4.686 | 1.816 | 2.02E-02 | 5.11E-02 |
| CDC23 | 9.579 | 2.124 | 12.44 | 3.389 | -1.045 | 4.93E-01 | 6.18E-01 |
| CDC26 | 22.758 | 9.452 | 23.77 | 4.493 | -1.221 | 8.68E-02 | 1.64E-01 |
| ESPL1 | 4.952 | 4.22 | 2.72 | 3.207 | -1.57 | 6.59E-02 | 1.32E-01 |
| FZR1 | 11.462 | 2.882 | 11.72 | 1.017 | -1.11 | 1.75E-01 | 2.82E-01 |
| MAD1L1 | 5.487 | 1.323 | 6.77 | 3.95 | 1.119 | 2.52E-01 | 3.75E-01 |
| MAD2L1 | 2.1 | 0.623 | 2.49 | 2.191 | 1.35 | 4.30E-02 | 9.42E-02 |
| MAD2L2 | 12.352 | 4.834 | 14.05 | 5.011 | -1.016 | 9.01E-01 | 9.37E-01 |
| MCM3 | 9.992 | 3.14 | 17.38 | 9.167 | 1.239 | 2.80E-02 | 6.65E-02 |
| MCM4 | 4.754 | 1.823 | 6.87 | 3.314 | 1.23 | 2.75E-02 | 6.56E-02 |
| MCM7 | 20.872 | 6.245 | 27.79 | 6.953 | 1.172 | 3.11E-02 | 7.25E-02 |
| ORC1 | 0.514 | 0.164 | 0.6 | 0.554 | 1.093 | 6.32E-01 | 7.38E-01 |
| ORC2 | 3.263 | 1.078 | 3.95 | 0.712 | 1.109 | 1.75E-01 | 2.82E-01 |
| ORC3 | 6.924 | 1.355 | 8.4 | 1.582 | -1.02 | 7.80E-01 | 8.52E-01 |
| ORC4 | 15.711 | 9.386 | 16.29 | 8.424 | -1.153 | 3.32E-02 | 7.64E-02 |
| ORC5 | 5.36 | 0.588 | 5.71 | 0.762 | -1.041 | 6.37E-01 | 7.42E-01 |
| ORC6 | 3.762 | 0.968 | 4.63 | 2.236 | 1.166 | 2.51E-01 | 3.74E-01 |
| RAD21 | 28.807 | 11.574 | 37.21 | 6.233 | 1.088 | 2.60E-01 | 3.84E-01 |
| SMC1A | 6.17 | 2.654 | 8.01 | 2.497 | 1.127 | 1.66E-01 | 2.71E-01 |
| STAG1 | 5.076 | 3.211 | 6.67 | 2.068 | 1.183 | 6.89E-02 | 1.37E-01 |
| WEE1 | 5.181 | 3.862 | 5.75 | 4.029 | 1.023 | 8.45E-01 | 8.99E-01 |
| YWHAB | 63.672 | 8.755 | 77.79 | 10.127 | 1.08 | 3.57E-01 | 4.86E-01 |
| YWHAG | 22.746 | 3.712 | 30.89 | 3.103 | 1.131 | 8.26E-02 | 1.58E-01 |
| YWHAQ | 62.71 | 18.625 | 77.15 | 30.768 | 1.034 | 6.45E-01 | 7.49E-01 |
| ***Damage Control*** | | | | | | | |
| ATM | 2.319 | 1.873 | 5.57 | 1.173 | 1.811 | 2.11E-08 | 3.68E-07 |
| ATR | 9.808 | 8.001 | 14.09 | 13.147 | 1.298 | 3.73E-04 | 1.92E-03 |
| PRKDC | 17.607 | 6.906 | 23.91 | 8.145 | 1.221 | 1.06E-02 | 3.02E-02 |
| TP53 | 7.653 | 1.938 | 13.26 | 5.066 | 1.401 | 7.56E-05 | 4.95E-04 |
| GADD45A | 32.842 | 22.036 | 23.2 | 9.874 | -1.547 | 3.33E-03 | 1.17E-02 |
| CHEK1 | 0 | 0 | 0 | 0 | 1.037 | 7.62E-01 | 8.39E-01 |
| CHEK2 | 0 | 1.233 | 2.5 | 1.945 | 1.232 | 6.74E-02 | 1.34E-01 |
| CREBBP | 5.306 | 4.899 | 8.6 | 3.493 | 1.171 | 1.00E-01 | 1.83E-01 |
| GADD45B | 16.055 | 10.278 | 26.06 | 13.465 | 1.217 | 9.14E-02 | 1.71E-01 |

Supplemental Table S7. Correlation Analysis of Cyclins.

Table S7A. D Cyclins Vs. Growth Factor CSF1, T-Cell activation pathways, CDKs, Pocket Protein Genes, E2F Expression Factors, and the Mini Chromosome Maintenance complex

| Gene | FC | FDR | CCND2 | CCND3 |
|---|---|---|---|---|
| CSF1 | 1.85 | 3.10E-08 | 0.821 | 0.872 |
| CSF1R | 4.11 | 9.78E-13 | 0.831 | 0.850 |
| IRAK4 | 1.54 | 6.94E-07 | 0.911 | 0.754 |
| PPP3R1 | 1.20 | 3.00E-02 | 0.899 | 0.770 |
| NFATC1 | 1.63 | 7.91E-04 | 0.815 | 0.833 |
| NFATC2 | 3.68 | 1.31E-11 | 0.898 | 0.861 |
| PIK3R5 | 7.24 | 2.12E-17 | 0.820 | 0.862 |
| TGFB1 | 1.72 | 1.05E-05 | 0.781 | 0.927 |
| TNFRSF10B | 1.51 | 9.68E-07 | 0.817 | 0.743 |
| CDK4 | -1.30 | 9.60E-03 | -0.035 | -0.069 |
| CDK6 | 1.30 | 5.08E-02 | 0.640 | 0.390 |
| RB1 | 1.56 | 5.46E-05 | 0.854 | 0.571 |
| RBL1 | 1.58 | 2.21E-03 | 0.873 | 0.709 |
| RBL2 | 1.15 | 1.83E-01 | 0.526 | 0.271 |
| E2F1 | 2.11 | 1.34E-03 | 0.706 | 0.752 |
| E2F2 | 4.26 | 3.01E-06 | 0.742 | 0.790 |
| MCM5 | 1.55 | 6.85E-04 | 0.810 | 0.911 |
| MCM6 | 1.72 | 1.01E-04 | 0.788 | 0.832 |
| EIF2AK3 | 1.61 | 8.64E-06 | 0.890 | 0.629 |

Table S7B. G1 Phase E Cyclins, and S Phase A and B Cyclins Vs. E2F Cell Cycle Transcription Factors

| Gene | FC | FDR | E2F1 | E2F2 |
|---|---|---|---|---|
| CCNE1 | 1.82 | 6.79E-03 | 0.896 | 0.908 |
| CCNE2 | 1.43 | 9.12E-03 | 0.743 | 0.734 |
| CCNA1 | 1.03 | 9.59E-01 | 0.520 | 0.932 |
| CCNA2 | 1.95 | 8.76E-05 | 0.936 | 0.942 |
| CCNB1 | 1.64 | 4.90E-03 | 0.510 | 0.932 |
| CCNB2 | 3.12 | 3.01E-05 | 0.507 | 0.942 |
| CCNB3 | -1.02 | 9.55E-01 | 0.251 | 0.097 |

Table S7C. G1 Phase E Cyclins, and S Phase A and B Cyclins Vs. E2F Cell Cycle Transcription Factors

| Gene | FC | FDR | MCM5 | MCM6 |
|---|---|---|---|---|
| CCNE1 | 1.82 | 6.79E-03 | 0.913 | 0.795 |
| CCNE2 | 1.43 | 9.12E-03 | 0.734 | 0.787 |
| CCNA2 | 1.95 | 8.76E-05 | 0.914 | 0.919 |
| CCNB1 | 1.64 | 4.90E-03 | 0.870 | 0.835 |
| CCNB2 | 3.12 | 3.01E-05 | 0.883 | 0.818 |
| CASP1 | 3.94 | 5.86E-15 | 0.903 | 0.946 |
| CASP3 | 1.94 | 2.26E-09 | 0.919 | 0.909 |
| CASP8 | 1.65 | 8.36E-07 | 0.821 | 0.921 |
| BCL2A1 | 9.43 | 4.97E-10 | 0.905 | 0.899 |
| BIRC3 | 5.74 | 5.74E-21 | 0.865 | 0.911 |
| BCL3 | 2.48 | 1.13E-10 | 0.875 | 0.911 |
| BTK | 6.97 | 9.11E-16 | 0.891 | 0.916 |
| **Gene** | **FC** | **FDR** | **CFLAR** | **PAK2** |
| ANAPC4 | 1.23 | 7.29E-03 | 0.919 | 0.926 |
| ATM | 1.81 | 3.68E-07 | 0.965 | 0.942 |
| CDC27 | 1.3 | 1.27E-03 | 0.913 | 0.960 |
| RBL1 | 1.58 | 2.21E-03 | 0.912 | 0.924 |

Supplemental Table S8. Intragraft Expression of mRNAs Encoding p53 KEGG Pathway Genes in TCMR and Normal Kidney Allograft Biopsies.

| Symbol | Normal Allograft Median | Normal Allograft IQR | TCMR Median | TCMR IQR | FC | PValue | FDR |
|---|---|---|---|---|---|---|---|
| ATM | 2.319 | 1.873 | 5.57 | 1.173 | 1.811 | 2.11E-08 | 3.68E-07 |
| ATR | 9.808 | 8.001 | 14.09 | 13.147 | 1.298 | 3.73E-04 | 1.92E-03 |
| CDKN1A | 9.667 | 4.757 | 20.28 | 7.522 | 1.69 | 2.97E-04 | 1.58E-03 |
| GORAB | 2.569 | 1.294 | 4.25 | 1.259 | 1.33 | 4.32E-03 | 1.46E-02 |
| MDM2 | 4.238 | 2.078 | 7.06 | 4.516 | 1.364 | 2.19E-05 | 1.72E-04 |
| MDM4 | 4.541 | 3.256 | 8.29 | 2.629 | 1.416 | 3.21E-04 | 1.69E-03 |
| TP53 | 7.653 | 1.938 | 13.26 | 5.066 | 1.401 | 7.56E-05 | 4.95E-04 |
| APAF1 | 1.651 | 0.933 | 3.29 | 2.033 | 1.72 | 2.82E-07 | 3.81E-06 |
| BBC3 | 1.889 | 0.945 | 3.98 | 1.508 | 1.867 | 1.23E-05 | 1.05E-04 |
| CASP3 | 4.287 | 1.107 | 9 | 4.449 | 1.943 | 7.99E-11 | 2.26E-09 |
| CASP8 | 6.403 | 1.768 | 12.16 | 6.979 | 1.645 | 5.30E-08 | 8.36E-07 |
| CCNB1 | 3.393 | 0.868 | 5.36 | 3.547 | 1.64 | 1.16E-03 | 4.90E-03 |
| CCNB2 | 0.461 | 0.34 | 1.33 | 1.236 | 3.117 | 2.95E-06 | 3.01E-05 |
| CCND2 | 1.561 | 1.017 | 5.08 | 2.698 | 2.689 | 9.99E-10 | 2.30E-08 |
| CCND3 | 17.599 | 4.399 | 26.6 | 11.116 | 1.338 | 2.82E-03 | 1.03E-02 |
| CCNE1 | 0.663 | 0.232 | 1.13 | 0.728 | 1.821 | 1.70E-03 | 6.79E-03 |
| CCNE2 | 0 | 0.616 | 1.12 | 1.872 | 1.43 | 2.44E-03 | 9.12E-03 |
| CDK1 | 1.779 | 0.943 | 2.81 | 4.143 | 1.828 | 6.76E-04 | 3.16E-03 |
| CDK2 | 4.418 | 2.53 | 9.79 | 11.008 | 1.438 | 1.64E-04 | 9.50E-04 |
| DDB2 | 4.904 | 1.614 | 8.54 | 3.718 | 1.496 | 1.58E-04 | 9.20E-04 |
| FAS | 0 | 0 | 0 | 6.611 | 1.814 | 8.48E-07 | 1.00E-05 |
| GTSE1 | 0.365 | 0.182 | 0.73 | 1.23 | 3.398 | 5.02E-06 | 4.81E-05 |
| PMAIP1 | 0.282 | 0.425 | 1.23 | 1.31 | 3.821 | 4.45E-09 | 8.94E-08 |
| RRM2 | 0.824 | 0.182 | 2.29 | 6.599 | 3.553 | 2.15E-07 | 2.98E-06 |
| SERPINE1 | 1.108 | 1.385 | 4.81 | 4.551 | 7.372 | 1.30E-10 | 3.53E-09 |
| SESN3 | 4.363 | 1.505 | 6.15 | 3.003 | 1.317 | 1.39E-02 | 3.77E-02 |
| SFN | 0.372 | 0.47 | 1.9 | 1.13 | 5.582 | 2.26E-07 | 3.11E-06 |
| THBS1 | 23.284 | 8.944 | 32.77 | 19.351 | 1.514 | 1.46E-03 | 5.97E-03 |
| TNFRSF10B | 10.647 | 2.759 | 18.09 | 3.416 | 1.518 | 6.22E-08 | 9.68E-07 |
| TP73 | 0 | 0 | 0 | 0 | 1.549 | 1.65E-02 | 4.34E-02 |
| ZMAT3 | 1.929 | 0.766 | 3.13 | 1.003 | 1.357 | 1.79E-03 | 7.07E-03 |
| CASP9 | 12.664 | 2.724 | 10.83 | 3.369 | -1.453 | 6.65E-06 | 6.14E-05 |
| CCNG1 | 51.311 | 29.475 | 55.39 | 33.969 | -1.275 | 1.65E-02 | 4.33E-02 |
| CDK4 | 34.052 | 13.757 | 32.44 | 28.958 | -1.303 | 2.60E-03 | 9.60E-03 |
| CYCS | 65.582 | 18.776 | 66.06 | 25.758 | -1.308 | 3.17E-03 | 1.13E-02 |
| GADD45A | 32.842 | 22.036 | 23.2 | 9.874 | -1.547 | 3.33E-03 | 1.17E-02 |
| IGFBP3 | 55.935 | 22.456 | 47.4 | 24.215 | -1.538 | 7.31E-04 | 3.37E-03 |
| SESN2 | 17.951 | 9.427 | 11.69 | 5.691 | -1.817 | 4.32E-04 | 2.18E-03 |
| TP53I3 | 37.663 | 13.749 | 41.12 | 7.866 | -1.264 | 4.80E-03 | 1.59E-02 |
| CHEK2 | 0 | 1.233 | 2.5 | 1.945 | 1.232 | 6.74E-02 | 1.34E-01 |
| ADGRB1 | 0.338 | 0.346 | 0.59 | 0.492 | | 9.40E-02 | 1.75E-01 |
| BAX | 49.046 | 23.828 | 92.49 | 138.04 | 1.035 | 7.64E-01 | 8.41E-01 |
| CCND1 | 26.944 | 8.851 | 24.86 | 8.517 | -1.196 | 6.43E-02 | 1.30E-01 |
| CCNG2 | 20.984 | 7.415 | 26.12 | 6.356 | -1.094 | 2.20E-01 | 3.37E-01 |
| CDK6 | 1.452 | 0.98 | 2.32 | 0.775 | 1.304 | 2.01E-02 | 5.08E-02 |
| COP1 | 10.425 | 1.413 | 12.67 | 2.191 | | 2.67E-01 | 3.92E-01 |
| EI24 | 31.196 | 3.856 | 33.07 | 6.049 | -1.166 | 2.04E-02 | 5.14E-02 |
| GADD45B | 16.055 | 10.278 | 26.06 | 13.465 | 1.217 | 9.14E-02 | 1.71E-01 |
| GADD45G | 6.081 | 4.342 | 6.56 | 2.254 | -1.382 | 4.39E-02 | 9.57E-02 |
| IGF1 | 0.622 | 0.606 | 1.07 | 1.62 | 1.804 | 2.46E-02 | 6.00E-02 |
| PERP | 7.027 | 4.441 | 8.73 | 9.205 | 1.155 | 3.48E-01 | 4.77E-01 |
| PPM1D | 2.728 | 1.076 | 4.05 | 1.635 | 1.202 | 5.65E-02 | 1.17E-01 |
| PTEN | 7.28 | 2.066 | 10.28 | 1.512 | 1.156 | 2.19E-02 | 5.45E-02 |
| RCHY1 | 9.936 | 2.309 | 10.7 | 2.876 | -1.044 | 5.30E-01 | 6.51E-01 |
| RPRM | 0.442 | 1.011 | 0.35 | 0.795 | -1.444 | 2.89E-01 | 4.16E-01 |
| RRM2B | 6.864 | 2.777 | 7.87 | 3.225 | 1.083 | 4.46E-01 | 5.74E-01 |
| SESN1 | 8.059 | 1.482 | 10.57 | 2.83 | 1.112 | 1.40E-01 | 2.38E-01 |
| SHISA5 | 46.537 | 17.076 | 59.58 | 19.846 | 1.148 | 2.02E-01 | 3.15E-01 |
| SIAH1 | 10.699 | 3.372 | 13.39 | 2.079 | -1.008 | 9.07E-01 | 9.41E-01 |
| STEAP3 | 2.023 | 0.715 | 2.06 | 1.461 | -1.073 | 6.46E-01 | 7.50E-01 |
| TP53AIP1 | 0.34 | 0.256 | 0.24 | 0.284 | 1.04 | 8.53E-01 | 9.04E-01 |
| TSC2 | 26.451 | 13.534 | 15.89 | 8.758 | -1.12 | 2.18E-01 | 3.35E-01 |

Supplemental Table S9. Intragraft expression of mRNAs encoding of Inhibitory Receptors and Ligands and T cell Exhaustion in TCMR and Normal Kidney Allograft Biopsies.

Table S9A. Inhibitory Receptors and Ligands

| | Normal Allograft | | TCMR | | | | |
|---|---|---|---|---|---|---|---|
| Symbol | Median | IQR | Median | IQR | FC | PValue | FDR |
| CD274 | 0.6 | 0.3 | 2.2 | 2.3 | 3.9 | 2.6E-12 | 1.0E-10 |
| CD276 | 11.5 | 2.6 | 18.4 | 6.1 | 1.3 | 8.2E-03 | 2.4E-02 |
| CD28 | 0.4 | 0.3 | 1.9 | 1.4 | 5.5 | 6.4E-14 | 3.5E-12 |
| CD80 | 0.0 | 0.0 | 0.0 | 0.0 | 7.2 | 4.6E-08 | 7.3E-07 |
| CD86 | 0.9 | 0.5 | 5.4 | 7.0 | 7.7 | 3.9E-19 | 1.2E-16 |
| CTLA4 | 0.2 | 0.2 | 2.0 | 2.5 | 13.1 | 1.6E-12 | 6.4E-11 |
| ICOS | 0.1 | 0.1 | 1.6 | 2.0 | 15.2 | 1.1E-15 | 9.8E-14 |
| ICOSLG | 5.8 | 2.1 | 7.7 | 2.6 | 1.1 | 6.2E-01 | NS |
| LAG3 | 0.7 | 0.4 | 2.5 | 3.6 | 10.5 | 3.8E-13 | 1.7E-11 |
| PDCD1 | 0.3 | 0.2 | 2.0 | 2.1 | 15.3 | 2.1E-14 | 1.3E-12 |
| PDCD1LG2 | 0.6 | 0.4 | 2.3 | 2.8 | 4.5 | 1.9E-11 | 6.2E-10 |
| TREML2 | 0.1 | 0.1 | 0.2 | 0.4 | 5.5 | 3.0E-07 | 4.0E-06 |
| VTCN1 | 7.0 | 1.9 | 8.9 | 3.4 | 1.3 | 7.7E-02 | NS |

Table S9B. T cell Exhaustion

| Symbol | Normal Allograft Median | Normal Allograft IQR | TCMR Median | TCMR IQR | FC | PValue | FDR |
|---|---|---|---|---|---|---|---|
| CD244 | 0.2 | 0.2 | 1.1 | 1.7 | 9.3 | 8.3E-15 | 5.9E-13 |
| CD27 | 0.0 | 0.0 | 0.0 | 23.0 | 2.7 | 6.5E-09 | 1.3E-07 |
| CD44 | 7.1 | 4.4 | 21.0 | 17.9 | 2.6 | 3.3E-10 | 8.3E-09 |
| CD69 | 0.5 | 0.4 | 2.8 | 5.5 | 6.5 | 6.7E-11 | 1.9E-09 |
| CXCR3 | 0.5 | 0.4 | 3.6 | 7.0 | 8.8 | 4.8E-14 | 2.8E-12 |
| EOMES | 0.2 | 0.2 | 1.9 | 2.0 | 11.2 | 5.1E-16 | 5.2E-14 |
| FOXP3 | 0.0 | 0.0 | 0.0 | 1.2 | 2.4 | 1.9E-05 | 1.5E-04 |
| GZMB | 0.7 | 1.2 | 5.8 | 15.6 | 10.0 | 3.7E-08 | 6.0E-07 |
| IL15 | 4.0 | 1.8 | 7.1 | 4.7 | 1.6 | 2.7E-05 | 2.1E-04 |
| IL15RA | 2.9 | 1.2 | 7.6 | 7.4 | 2.9 | 4.4E-10 | 1.1E-08 |
| IL2RA | 0.4 | 0.2 | 1.2 | 2.4 | 6.1 | 1.1E-11 | 3.6E-10 |
| IL2RB | 0.5 | 0.3 | 5.4 | 8.3 | 12.0 | 1.0E-18 | 2.6E-16 |
| IL4R | 7.2 | 2.0 | 14.4 | 7.3 | 2.0 | 2.7E-11 | 8.5E-10 |
| IL7 | 0.7 | 0.8 | 1.4 | 0.7 | 1.5 | 6.7E-04 | 3.2E-03 |
| IL7R | 0.5 | 0.9 | 7.1 | 7.0 | 7.6 | 3.0E-12 | 1.2E-10 |
| KLRG1 | 1.4 | 0.6 | 3.8 | 2.9 | 2.6 | 2.9E-07 | 3.9E-06 |
| PRDM1 | 0.4 | 0.2 | 1.7 | 1.4 | 5.1 | 5.2E-14 | 3.0E-12 |
| SELL | 1.8 | 1.2 | 8.2 | 12.0 | 6.4 | 3.3E-15 | 2.7E-13 |
| SPN | 0.0 | 0.0 | 1.5 | 6.9 | 7.8 | 4.7E-17 | 7.2E-15 |
| LY6G5C | 1.8 | 0.7 | 1.9 | 1.0 | -1.1 | 5.1E-01 | NS |

Supplemental Table S10. Intragraft Expression of mRNAs Encoding Apoptosis KEGG Pathway Genes.

| Pathway | Symbol | Normal Allograft Median | Normal Allograft IQR | TCMR Median | TCMR IQR | FC | PValue | FDR |
|---|---|---|---|---|---|---|---|---|
| Extrinsic | FASLG | 0.1 | 0.1 | 1.5 | 2.0 | 14.5 | 4.4E-17 | 6.7E-15 |
| Extrinsic | TNF | 0.4 | 0.3 | 2.4 | 2.0 | 5.3 | 2.3E-14 | 1.4E-12 |
| Extrinsic | BCL2A1 | 0.5 | 0.2 | 4.2 | 9.4 | 9.4 | 1.5E-11 | 5.0E-10 |
| Extrinsic | TRAF1 | 3.4 | 0.7 | 9.9 | 7.4 | 2.4 | 5.3E-11 | 1.6E-09 |
| Extrinsic | JUN | 7.4 | 4.2 | 16.4 | 3.6 | 1.9 | 1.5E-09 | 3.2E-08 |
| Extrinsic | ATM | 2.3 | 1.9 | 5.6 | 1.2 | 1.8 | 2.1E-08 | 3.7E-07 |
| Extrinsic | CASP8 | 6.4 | 1.8 | 12.2 | 7.0 | 1.6 | 5.3E-08 | 8.4E-07 |
| Extrinsic | TNFRSF10B | 10.6 | 2.8 | 18.1 | 3.4 | 1.5 | 6.2E-08 | 9.7E-07 |
| Extrinsic | MAP3K14 | 6.1 | 2.1 | 8.9 | 3.9 | 1.6 | 6.1E-07 | 7.5E-06 |
| Extrinsic | NFKB1 | 8.3 | 3.7 | 15.3 | 4.6 | 1.5 | 7.5E-07 | 9.0E-06 |
| Extrinsic | CASP10 | 3.4 | 1.6 | 6.2 | 3.2 | 1.6 | 8.4E-07 | 9.9E-06 |
| Extrinsic | FAS | 0.0 | 0.0 | 0.0 | 6.6 | 1.8 | 8.5E-07 | 1.0E-05 |
| Extrinsic | IKBKB | 9.5 | 4.4 | 16.7 | 4.0 | 1.5 | 2.9E-06 | 3.0E-05 |
| Extrinsic | ITPR3 | 2.3 | 3.7 | 8.0 | 7.1 | 2.0 | 5.2E-06 | 5.0E-05 |
| Extrinsic | ERN1 | 1.9 | 1.0 | 3.3 | 0.6 | 1.5 | 1.7E-05 | 1.4E-04 |
| Extrinsic | TNFRSF10A | 2.0 | 0.8 | 4.1 | 1.6 | 1.7 | 2.4E-05 | 1.8E-04 |
| Extrinsic | CTSF | 46.1 | 10.2 | 35.0 | 16.3 | -1.6 | 3.1E-05 | 2.3E-04 |
| Extrinsic | NFKBIA | 27.7 | 11.1 | 54.8 | 19.9 | 1.5 | 4.2E-05 | 3.0E-04 |
| Extrinsic | TP53 | 7.7 | 1.9 | 13.3 | 5.1 | 1.4 | 7.6E-05 | 4.9E-04 |
| Extrinsic | TNFRSF10C | 1.3 | 1.2 | 2.7 | 1.4 | 1.6 | 8.5E-04 | 3.8E-03 |
| Extrinsic | CFLAR | 9.3 | 5.1 | 14.5 | 3.0 | 1.3 | 1.7E-03 | 6.7E-03 |
| Extrinsic | TNFSF10 | 79.4 | 27.5 | 144.1 | 76.6 | 1.6 | 1.9E-03 | 7.3E-03 |
| Extrinsic | MAP3K5 | 5.2 | 1.5 | 8.1 | 2.3 | 1.3 | 1.9E-03 | 7.5E-03 |
| Extrinsic | CASP7 | 8.1 | 2.8 | 11.7 | 5.2 | 1.3 | 5.0E-03 | 1.6E-02 |
| Extrinsic | TRAF2 | 5.5 | 1.6 | 7.3 | 4.8 | 1.4 | 5.4E-03 | 1.7E-02 |
| Extrinsic | DFFA | 9.4 | 6.0 | 7.1 | 5.3 | -1.2 | 6.1E-03 | 1.9E-02 |
| Extrinsic | RIPK1 | 9.1 | 1.5 | 12.3 | 2.9 | 1.2 | 1.4E-02 | 3.8E-02 |
| Extrinsic | BAK1 | 6.5 | 1.5 | 8.9 | 2.9 | 1.3 | 2.1E-02 | NS |
| Extrinsic | TNFRSF1A | 39.0 | 18.6 | 41.2 | 48.4 | 1.3 | 2.3E-02 | NS |
| Extrinsic | DAXX | 11.7 | 1.1 | 17.8 | 5.4 | 1.2 | 2.9E-02 | NS |
| Extrinsic | HRK | 0.1 | 0.1 | 0.1 | 0.1 | 1.5 | 1.2E-01 | NS |
| Extrinsic | CHUK | 8.6 | 1.1 | 7.8 | 2.1 | -1.1 | 2.9E-01 | NS |
| Extrinsic | RELA | 18.1 | 3.1 | 23.1 | 6.8 | 1.1 | 3.0E-01 | NS |
| Extrinsic | CASP6 | 10.3 | 3.4 | 11.3 | 4.3 | 1.1 | 5.9E-01 | NS |
| Extrinsic | PIDD1 | 0.0 | 0.0 | 0.0 | 0.0 | -1.0 | 6.5E-01 | NS |
| Extrinsic | ITPR2 | 3.2 | 2.1 | 4.0 | 1.8 | 1.0 | 6.5E-01 | NS |
| Extrinsic | IKBKG | 7.3 | 2.9 | 9.1 | 4.5 | 1.0 | 6.7E-01 | NS |
| Extrinsic | DAB2IP | 14.4 | 2.7 | 14.8 | 3.3 | -1.0 | 7.6E-01 | NS |
| Extrinsic | BAX | 49.0 | 23.8 | 92.5 | 138.0 | 1.0 | 7.6E-01 | NS |
| Extrinsic | TP53AIP1 | 0.3 | 0.3 | 0.2 | 0.3 | 1.0 | 8.5E-01 | NS |
| Extrinsic | TNFRSF10D | 4.1 | 0.9 | 4.6 | 1.7 | 1.0 | 8.6E-01 | NS |
| Extrinsic | ITPR1 | 9.1 | 5.8 | 9.8 | 4.8 | 1.0 | 8.8E-01 | NS |
| Extrinsic | DFFB | 3.0 | 0.5 | 3.6 | 0.4 | -1.0 | 9.4E-01 | NS |
| Extrinsic | FADD | 2.0 | 0.6 | 3.4 | 1.2 | -1.0 | 9.4E-01 | NS |
| Extrinsic | TRADD | 24.8 | 7.1 | 26.9 | 7.6 | 1.0 | 9.9E-01 | NS |
| Intrinsic | CTSS | 9.6 | 4.9 | 85.8 | 126.7 | 8.3 | 1.3E-22 | 5.5E-19 |
| Intrinsic | CTSW | 0.0 | 0.0 | 12.3 | 24.6 | 7.1 | 1.7E-10 | 4.5E-09 |
| Intrinsic | PMAIP1 | 0.3 | 0.4 | 1.2 | 1.3 | 3.8 | 4.4E-09 | 8.9E-08 |
| Intrinsic | MCL1 | 30.0 | 9.3 | 57.3 | 23.3 | 1.8 | 2.5E-08 | 4.3E-07 |
| Intrinsic | APAF1 | 1.7 | 0.9 | 3.3 | 2.0 | 1.7 | 2.8E-07 | 3.8E-06 |
| Intrinsic | CASP9 | 12.7 | 2.7 | 10.8 | 3.4 | -1.5 | 6.6E-06 | 6.1E-05 |
| Intrinsic | BBC3 | 1.9 | 0.9 | 4.0 | 1.5 | 1.9 | 1.2E-05 | 1.0E-04 |
| Intrinsic | CTSK | 6.5 | 3.6 | 15.0 | 9.6 | 1.9 | 2.0E-05 | 1.6E-04 |
| Intrinsic | ENDOG | 6.2 | 7.6 | 5.7 | 7.5 | -1.5 | 4.0E-04 | 2.0E-03 |
| Intrinsic | BCL2L11 | 4.0 | 2.1 | 6.5 | 1.0 | 1.3 | 4.8E-04 | 2.4E-03 |
| Intrinsic | CTSZ | 75.8 | 28.4 | 120.8 | 79.2 | 1.7 | 5.5E-04 | 2.7E-03 |
| Intrinsic | CTSV | 13.8 | 11.7 | 5.0 | 8.3 | -2.4 | 1.9E-03 | 7.4E-03 |
| Intrinsic | AIFM1 | 128.3 | 63.4 | 86.2 | 49.9 | -1.7 | 1.9E-03 | 7.5E-03 |
| Intrinsic | CTSO | 13.8 | 7.3 | 23.0 | 3.8 | 1.3 | 3.1E-03 | 1.1E-02 |
| Intrinsic | CYCS | 65.6 | 18.8 | 66.1 | 25.8 | -1.3 | 3.2E-03 | 1.1E-02 |
| Intrinsic | MAPK10 | 11.9 | 3.4 | 12.0 | 4.0 | -1.3 | 6.8E-03 | 2.1E-02 |
| Intrinsic | DDIT3 | 28.7 | 6.0 | 0.0 | 24.9 | -1.3 | 1.9E-02 | 4.8E-02 |
| Intrinsic | CTSH | 464.1 | 227.9 | 449.6 | 146.4 | -1.4 | 2.7E-02 | NS |
| Intrinsic | HTRA2 | 0.0 | 0.0 | 0.0 | 0.0 | -1.2 | 3.4E-02 | NS |
| Intrinsic | CTSB | 689.6 | 432.1 | 497.7 | 584.7 | -1.5 | 5.1E-02 | NS |
| Intrinsic | BID | 5.5 | 1.8 | 8.3 | 8.8 | 1.2 | 5.2E-02 | NS |
| Intrinsic | CTSL | 137.5 | 32.4 | 133.8 | 72.9 | -1.3 | 5.6E-02 | NS |
| Intrinsic | MAPK8 | 5.5 | 1.4 | 6.1 | 1.6 | -1.1 | 1.4E-01 | NS |
| Intrinsic | CTSC | 132.2 | 24.2 | 154.2 | 82.3 | 1.2 | 2.4E-01 | NS |
| Intrinsic | CTSD | 430.0 | 217.6 | 511.9 | 243.0 | -1.1 | 4.5E-01 | NS |
| Intrinsic | SEPT4 | 19.2 | 5.6 | 21.6 | 7.4 | -1.1 | 5.4E-01 | NS |
| Intrinsic | EIF2S1 | 6.5 | 1.2 | 8.0 | 1.7 | 1.0 | 6.3E-01 | NS |
| Intrinsic | ATF4 | 101.7 | 25.6 | 116.5 | 35.0 | -1.0 | 6.8E-01 | NS |
| Intrinsic | DIABLO | 41.7 | 12.3 | 34.1 | 14.4 | 1.0 | 8.5E-01 | NS |
| Intrinsic | MAPK9 | 7.3 | 2.1 | 9.3 | 5.5 | 1.0 | 8.5E-01 | NS |

| Function | Symbol | Normal Allograft Median | IQR | TCMR Median | IQR | FC | PValue | FDR |
|---|---|---|---|---|---|---|---|---|
| Survival Factors | BIRC3 | 1.7 | 0.6 | 10.3 | 12.2 | 5.7 | 3.5E-25 | 5.7E-21 |
| Survival Factors | PIK3R5 | 0.7 | 0.4 | 3.3 | 4.1 | 7.2 | 3.9E-20 | 2.1E-17 |
| Survival Factors | CSF2RB | 0.8 | 0.7 | 4.4 | 6.6 | 6.4 | 8.1E-16 | 7.7E-14 |
| Survival Factors | PIK3CG | 0.3 | 0.4 | 1.6 | 1.2 | 4.4 | 1.2E-10 | 3.2E-09 |
| Survival Factors | PIK3CD | 0.0 | 0.0 | 8.0 | 6.5 | 2.0 | 1.3E-07 | 2.0E-06 |
| Survival Factors | NRAS | 7.3 | 1.2 | 10.9 | 2.6 | 1.3 | 6.1E-04 | 2.9E-03 |
| Survival Factors | BCL2L1 | 53.1 | 8.8 | 46.6 | 8.1 | -1.3 | 8.0E-04 | 3.6E-03 |
| Survival Factors | GADD45A | 32.8 | 22.0 | 23.2 | 9.9 | -1.5 | 3.3E-03 | 1.2E-02 |
| Survival Factors | IL3RA | 3.2 | 1.3 | 5.6 | 5.1 | 1.4 | 8.4E-03 | 2.5E-02 |
| Survival Factors | BAD | 25.1 | 11.7 | 25.8 | 5.5 | -1.4 | 8.9E-03 | 2.6E-02 |
| Survival Factors | BCL2 | 3.0 | 2.3 | 4.8 | 1.8 | 1.3 | 1.1E-02 | 3.1E-02 |
| Survival Factors | FOS | 2.2 | 1.9 | 6.0 | 3.7 | 1.7 | 1.4E-02 | 3.8E-02 |
| Survival Factors | HRAS | 26.8 | 5.5 | 25.6 | 7.1 | -1.3 | 1.6E-02 | 4.3E-02 |
| Survival Factors | KRAS | 4.8 | 2.0 | 6.9 | 2.4 | 1.2 | 1.7E-02 | 4.5E-02 |
| Survival Factors | PIK3CA | 2.2 | 1.2 | 3.2 | 0.9 | 1.2 | 2.6E-02 | NS |
| Survival Factors | MAPK3 | 34.1 | 13.3 | 37.8 | 14.8 | -1.2 | 3.6E-02 | NS |
| Survival Factors | GADD45G | 6.1 | 4.3 | 6.6 | 2.3 | -1.4 | 4.4E-02 | NS |
| Survival Factors | XIAP | 6.3 | 1.6 | 7.9 | 2.0 | 1.1 | 5.1E-02 | NS |
| Survival Factors | BIRC2 | 15.8 | 1.2 | 19.4 | 3.5 | 1.1 | 6.3E-02 | NS |
| Survival Factors | AKT3 | 4.0 | 2.9 | 6.2 | 3.5 | 1.2 | 9.1E-02 | NS |
| Survival Factors | GADD45B | 16.1 | 10.3 | 26.1 | 13.5 | 1.2 | 9.1E-02 | NS |
| Survival Factors | NTRK1 | 0.9 | 1.4 | 0.7 | 1.1 | -1.5 | 1.2E-01 | NS |
| Survival Factors | RAF1 | 23.6 | 5.6 | 26.2 | 4.1 | -1.1 | 1.8E-01 | NS |
| Survival Factors | PIK3CB | 10.7 | 2.4 | 10.8 | 2.7 | -1.1 | 2.2E-01 | NS |
| Survival Factors | AKT1 | 33.7 | 5.5 | 43.9 | 12.4 | 1.1 | 2.3E-01 | NS |
| Survival Factors | MAPK1 | 7.7 | 3.2 | 10.3 | 2.7 | 1.1 | 2.6E-01 | NS |
| Survival Factors | NGF | 1.2 | 0.7 | 1.3 | 0.7 | -1.2 | 3.4E-01 | NS |
| Survival Factors | MAP2K2 | 41.0 | 14.1 | 43.5 | 12.7 | -1.1 | 3.5E-01 | NS |
| Survival Factors | PIK3R3 | 5.7 | 1.5 | 7.1 | 3.3 | 1.1 | 4.1E-01 | NS |
| Survival Factors | MAP2K1 | 19.3 | 10.1 | 29.3 | 9.0 | 1.0 | 4.3E-01 | NS |
| Survival Factors | PIK3R1 | 9.1 | 3.0 | 11.4 | 4.1 | 1.1 | 4.4E-01 | NS |
| Survival Factors | PDPK1 | 7.8 | 2.4 | 9.3 | 2.0 | 1.0 | 5.9E-01 | NS |
| Survival Factors | AKT2 | 35.1 | 15.7 | 39.0 | 12.1 | -1.0 | 6.7E-01 | NS |
| Survival Factors | PTPN13 | 6.1 | 6.5 | 8.2 | 6.3 | 1.1 | 7.2E-01 | NS |
| Cytotoxic granules | PRF1 | 0.8 | 0.5 | 4.6 | 9.9 | 9.7 | 2.6E-13 | 1.2E-11 |
| Cytotoxic granules | LMNB1 | 1.4 | 0.6 | 5.0 | 5.0 | 4.4 | 8.2E-13 | 3.5E-11 |
| Cytotoxic granules | GZMB | 0.7 | 1.2 | 5.8 | 15.6 | 10.0 | 3.7E-08 | 6.0E-07 |
| Cytotoxic granules | TUBA1A | 41.0 | 7.4 | 84.8 | 46.3 | 1.7 | 1.7E-05 | 1.4E-04 |
| Cytotoxic granules | BIRC5 | 0.8 | 0.5 | 1.5 | 1.7 | 2.5 | 9.4E-05 | 5.9E-04 |
| Cytotoxic granules | LMNB2 | 4.2 | 0.8 | 5.9 | 2.4 | 1.4 | 2.9E-04 | 1.6E-03 |
| Cytotoxic granules | PARP1 | 15.8 | 2.7 | 21.4 | 4.2 | 1.3 | 2.3E-03 | 8.7E-03 |
| Cytotoxic granules | PARP4 | 12.8 | 4.4 | 19.4 | 3.6 | 1.3 | 2.5E-03 | 9.2E-03 |
| Cytotoxic granules | SPTA1 | 0.1 | 0.2 | 0.2 | 0.1 | 2.6 | 3.5E-03 | 1.2E-02 |
| Cytotoxic granules | TUBA4A | 43.5 | 11.8 | 43.2 | 7.4 | -1.4 | 3.7E-03 | 1.3E-02 |
| Cytotoxic granules | TUBA1B | 121.7 | 20.9 | 173.1 | 81.0 | 1.4 | 1.8E-02 | 4.6E-02 |
| Cytotoxic granules | TUBAL3 | 6.8 | 4.9 | 3.6 | 4.2 | -2.3 | 1.9E-02 | 4.9E-02 |
| Cytotoxic granules | PARP2 | 9.7 | 2.2 | 10.5 | 4.2 | -1.2 | 5.3E-02 | NS |
| Cytotoxic granules | ACTB | 613.1 | 123.5 | 764.1 | 477.7 | 1.3 | 6.4E-02 | NS |
| Cytotoxic granules | ACTG1 | 514.6 | 132.9 | 715.7 | 314.6 | 1.2 | 1.2E-01 | NS |
| Cytotoxic granules | TUBA1C | 51.0 | 11.9 | 70.1 | 44.3 | -1.1 | 2.8E-01 | NS |
| Cytotoxic granules | PARP3 | 17.2 | 2.8 | 18.5 | 6.1 | -1.1 | 4.5E-01 | NS |
| Cytotoxic granules | LMNA | 56.2 | 13.5 | 67.7 | 28.6 | 1.1 | 5.1E-01 | NS |
| Cytotoxic granules | SPTAN1 | 32.0 | 5.6 | 38.2 | 11.4 | -1.0 | 1.0E+00 | NS |
| Ca induced Cell Death | CASP3 | 4.3 | 1.1 | 9.0 | 4.4 | 1.9 | 8.0E-11 | 2.3E-09 |
| Ca induced Cell Death | EIF2AK3 | 2.5 | 1.4 | 5.2 | 1.6 | 1.6 | 7.2E-07 | 8.6E-06 |
| Ca induced Cell Death | CASP2 | 3.5 | 2.6 | 7.3 | 3.9 | 1.5 | 1.6E-04 | 9.2E-04 |
| Ca induced Cell Death | CAPN1 | 35.2 | 7.1 | 46.0 | 17.7 | 1.1 | 2.6E-01 | NS |
| Ca induced Cell Death | CAPN2 | 59.6 | 16.4 | 66.0 | 12.0 | -1.0 | 6.4E-01 | NS |
| Ca induced Cell Death | CASP12 | 0.3 | 0.5 | 0.3 | 0.4 | -1.1 | 7.6E-01 | NS |

Supplemental Table S11. Intragraft Expression of mRNAs Encoding Necroptosis KEGG Pathway Genes.

| Symbol | Normal Allograft Median | Normal Allograft IQR | TCMR Median | TCMR IQR | FC | PValue | FDR |
|---|---|---|---|---|---|---|---|
| ALOX15 | 0.07 | 0.063 | 0.26 | 0.309 | 4.478 | 4.05E-05 | 2.92E-04 |
| BCL2 | 3.035 | 2.275 | 4.79 | 1.815 | 1.272 | 1.11E-02 | 3.15E-02 |
| BIRC3 | 1.654 | 0.609 | 10.29 | 12.231 | 5.744 | 3.50E-25 | 5.74E-21 |
| CAMK2D | 6.086 | 2.562 | 9.58 | 2.72 | 1.292 | 1.59E-04 | 9.23E-04 |
| CASP1 | 4.933 | 0.921 | 21.45 | 16.697 | 3.94 | 3.72E-17 | 5.86E-15 |
| CASP8 | 6.403 | 1.768 | 12.16 | 6.979 | 1.645 | 5.30E-08 | 8.36E-07 |
| CFLAR | 9.31 | 5.085 | 14.49 | 3.03 | 1.322 | 1.68E-03 | 6.72E-03 |
| CYBB | 3.844 | 2.953 | 20.43 | 29.109 | 6.413 | 1.50E-18 | 3.75E-16 |
| CYLD | 3.138 | 0.931 | 7.72 | 5.13 | 1.904 | 1.96E-13 | 9.65E-12 |
| EIF2AK2 | 3.794 | 1.257 | 5.98 | 2.053 | 1.347 | 4.41E-04 | 2.21E-03 |
| FAS | 0 | 0 | 0 | 6.611 | 1.814 | 8.48E-07 | 1.00E-05 |
| FASLG | 0.146 | 0.067 | 1.45 | 1.958 | 14.474 | 4.38E-17 | 6.71E-15 |
| GLUL | 55.791 | 41.354 | 103.15 | 47.847 | 1.594 | 7.23E-05 | 4.78E-04 |
| H2AFX | 4.454 | 1.605 | 6.76 | 2.83 | 1.588 | 7.04E-04 | 3.27E-03 |
| H2AFY | 38.669 | 5.721 | 51.86 | 15.294 | 1.36 | 1.28E-04 | 7.67E-04 |
| HIST1H2AG | 0.11 | 0.102 | 0.32 | 0.218 | 2.572 | 5.13E-04 | 2.51E-03 |
| IFNAR1 | 8.635 | 7.29 | 13.37 | 6.758 | 1.252 | 3.32E-03 | 1.17E-02 |
| IFNAR2 | 1.192 | 3.462 | 4 | 6.906 | 2.739 | 2.70E-18 | 6.19E-16 |
| IFNG | 0.146 | 0.158 | 1.16 | 1.825 | 8.12 | 2.20E-08 | 3.84E-07 |
| IFNGR1 | 17.104 | 5.537 | 36.31 | 17.98 | 1.766 | 2.43E-11 | 7.64E-10 |
| IFNGR2 | 21.258 | 4.636 | 33.48 | 14.595 | 1.335 | 7.86E-04 | 3.58E-03 |
| IL1B | 0.318 | 0.227 | 1.84 | 2.683 | 5.078 | 1.49E-09 | 3.28E-08 |
| IRF9 | 19.191 | 30.376 | 25.29 | 99.942 | 1.977 | 1.90E-08 | 3.36E-07 |
| JAK1 | 18.401 | 9.895 | 36.75 | 15.367 | 1.593 | 4.57E-07 | 5.78E-06 |
| JAK2 | 2.079 | 1.564 | 5.5 | 5.409 | 2.145 | 2.73E-09 | 5.73E-08 |
| JAK3 | 0.917 | 0.525 | 6.64 | 7.831 | 8.212 | 2.76E-23 | 1.50E-19 |
| MLKL | 2.573 | 1.356 | 6.1 | 4.031 | 2.535 | 1.88E-10 | 4.91E-09 |
| NLRP3 | 0.368 | 0.234 | 1.29 | 1.325 | 3.824 | 1.35E-10 | 3.65E-09 |
| PARP1 | 15.819 | 2.666 | 21.41 | 4.189 | 1.269 | 2.30E-03 | 8.70E-03 |
| PARP4 | 12.774 | 4.432 | 19.43 | 3.569 | 1.278 | 2.47E-03 | 9.22E-03 |
| PLA2G4A | 2.076 | 1.238 | 5.25 | 4.293 | 1.742 | 5.67E-03 | 1.82E-02 |
| PLA2G4C | 8.861 | 3.268 | 16.03 | 9.819 | 1.442 | 1.15E-02 | 3.24E-02 |
| PYCARD | 3.376 | 1.313 | 11.21 | 12.632 | 4.103 | 9.64E-12 | 3.30E-10 |
| PYGL | 4.486 | 1.365 | 7.12 | 3.959 | 1.605 | 3.46E-04 | 1.80E-03 |
| RIPK1 | 9.095 | 1.477 | 12.27 | 2.916 | 1.176 | 1.42E-02 | 3.84E-02 |
| RIPK3 | 0 | 0 | 5.9 | 6.968 | 2.494 | 9.56E-10 | 2.21E-08 |
| RNF31 | 11.484 | 17.105 | 0 | 0 | 1.258 | 2.64E-03 | 9.70E-03 |
| STAT1 | 29.2 | 9.22 | 95.01 | 91.771 | 3.083 | 2.74E-11 | 8.46E-10 |
| STAT2 | 23.628 | 11.453 | 52.44 | 25.337 | 1.749 | 5.80E-07 | 7.13E-06 |
| STAT3 | 24.51 | 8.243 | 40.37 | 14.269 | 1.499 | 9.20E-06 | 8.13E-05 |
| STAT4 | 0.874 | 0.486 | 3.69 | 3.641 | 3.935 | 1.21E-13 | 6.21E-12 |
| STAT5A | 8.171 | 2.282 | 12.83 | 8.852 | 1.625 | 1.02E-04 | 6.37E-04 |
| STAT6 | 37.96 | 12.356 | 51.93 | 16.143 | 1.3 | 3.31E-03 | 1.17E-02 |
| TICAM1 | 3.881 | 0.907 | 5.94 | 2.046 | 1.345 | 4.54E-03 | 1.52E-02 |
| TICAM2 | 0 | 0 | 0 | 0 | 3.224 | 3.28E-05 | 2.43E-04 |
| TLR3 | 2.775 | 3.085 | 5.34 | 3.784 | 1.34 | 4.54E-03 | 1.52E-02 |
| TLR4 | 2.018 | 1.352 | 6.27 | 4.08 | 2.478 | 5.55E-10 | 1.35E-08 |
| TNF | 0.423 | 0.254 | 2.36 | 2.044 | 5.26 | 2.33E-14 | 1.43E-12 |
| TNFAIP3 | 2.171 | 0.844 | 8.63 | 8.172 | 3.827 | 4.45E-16 | 4.64E-14 |
| TNFRSF10A | 1.956 | 0.751 | 4.09 | 1.607 | 1.718 | 2.36E-05 | 1.83E-04 |
| TNFRSF10B | 10.647 | 2.759 | 18.09 | 3.416 | 1.518 | 6.22E-08 | 9.68E-07 |
| TNFRSF10C | 1.303 | 1.217 | 2.69 | 1.393 | 1.638 | 8.50E-04 | 3.82E-03 |
| TNFSF10 | 79.448 | 27.547 | 144.11 | 76.569 | 1.622 | 1.87E-03 | 7.32E-03 |
| TRAF2 | 5.474 | 1.62 | 7.31 | 4.763 | 1.376 | 5.37E-03 | 1.74E-02 |
| TRAF5 | 3.298 | 2.612 | 8.62 | 4.231 | 2.004 | 2.33E-09 | 4.94E-08 |
| VPS4B | 11.316 | 2.398 | 15.19 | 2.12 | 1.177 | 1.25E-02 | 3.46E-02 |
| ZBP1 | 0.318 | 0.139 | 2.34 | 2.849 | 11.185 | 3.19E-15 | 2.61E-13 |
| AIFM1 | 128.253 | 63.364 | 86.17 | 49.909 | -1.721 | 1.93E-03 | 7.52E-03 |
| CAMK2A | 0.554 | 0.229 | 0.35 | 0.359 | -1.742 | 9.20E-03 | 2.69E-02 |
| CAMK2G | 19.261 | 4.362 | 14.64 | 4.305 | -1.509 | 3.10E-05 | 2.32E-04 |
| CHMP2A | 114.522 | 46.315 | 99.14 | 17.49 | -1.476 | 1.12E-03 | 4.77E-03 |
| FTH1 | 1,485.91 | 1,032.03 | 1,391.04 | 755.067 | -1.636 | 1.56E-02 | 4.13E-02 |
| FTL | 6,769.91 | 5,906.51 | 3,986.05 | 3,103.41 | -2.205 | 3.03E-03 | 1.09E-02 |
| GLUD1 | 138.367 | 44.075 | 94.05 | 59.751 | -1.595 | 2.23E-03 | 8.47E-03 |
| GLUD2 | 4.607 | 2.138 | 4.38 | 4.327 | -1.714 | 1.79E-03 | 7.08E-03 |
| MAPK10 | 11.907 | 3.393 | 12.04 | 4.024 | -1.255 | 6.80E-03 | 2.11E-02 |
| PLA2G4F | 10.273 | 3.866 | 6.36 | 3.352 | -1.58 | 3.62E-04 | 1.87E-03 |
| PPID | 22.692 | 19.716 | 21.53 | 3.014 | -1.323 | 4.77E-04 | 2.36E-03 |
| SHARPIN | 22.017 | 4.194 | 19.87 | 3.542 | -1.32 | 2.21E-03 | 8.41E-03 |
| SLC25A4 | 48.948 | 8.677 | 38.01 | 16.203 | -1.595 | 2.26E-06 | 2.37E-05 |
| SLC25A5 | 371.224 | 104.916 | 246.6 | 80.171 | -1.781 | 8.84E-06 | 7.85E-05 |
| SLC25A6 | 195.406 | 45.156 | 184.79 | 30.888 | -1.397 | 4.36E-03 | 1.47E-02 |
| SMPD1 | 47.992 | 24.943 | 32.14 | 9.742 | -1.567 | 2.72E-04 | 1.46E-03 |
| VDAC1 | 103.702 | 16.948 | 97 | 25.768 | -1.373 | 2.37E-03 | 8.90E-03 |
| VDAC2 | 82.439 | 12.011 | 77.47 | 12.444 | -1.384 | 2.79E-05 | 2.12E-04 |
| VDAC3 | 57.685 | 7.626 | 58.18 | 8.54 | -1.188 | 1.80E-02 | 4.65E-02 |
| VPS4A | 63.209 | 13.543 | 56.67 | 13.969 | -1.325 | 2.59E-03 | 9.56E-03 |

| Symbol | Normal Allograft Median | IQR | TCMR Median | IQR | FC | PValue | FDR |
|---|---|---|---|---|---|---|---|
| BAX | 49.046 | 23.828 | 92.49 | 138.04 | 1.035 | 7.64E-01 | 8.41E-01 |
| BID | 5.503 | 1.777 | 8.3 | 8.795 | 1.243 | 5.23E-02 | 1.10E-01 |
| BIRC2 | 15.763 | 1.209 | 19.36 | 3.505 | 1.128 | 6.33E-02 | 1.28E-01 |
| CAMK2B | 0.962 | 0.342 | 0.78 | 0.527 | -1.549 | 2.25E-02 | 5.58E-02 |
| CAPN1 | 35.171 | 7.053 | 46.02 | 17.672 | 1.108 | 2.60E-01 | 3.84E-01 |
| CAPN2 | 59.558 | 16.432 | 66.02 | 12.036 | -1.043 | 6.42E-01 | 7.47E-01 |
| CHMP1A | 37.012 | 6.664 | 38.91 | 5.33 | -1.097 | 3.34E-01 | 4.63E-01 |
| CHMP1B | 13.902 | 2.209 | 15.6 | 2.591 | -1.023 | 7.25E-01 | 8.14E-01 |
| CHMP2B | 12.397 | 1.978 | 14.07 | 4.619 | -1.09 | 1.82E-01 | 2.91E-01 |
| CHMP3 | 22.89 | 4.71 | 25.74 | 8.457 | 1.016 | 8.05E-01 | 8.70E-01 |
| CHMP4B | 47.243 | 10.795 | 50.62 | 9.673 | -1.04 | 6.45E-01 | 7.49E-01 |
| CHMP4C | 5.273 | 1.105 | 7.18 | 2.874 | -1.002 | 9.86E-01 | 9.90E-01 |
| CHMP5 | 45.623 | 6.454 | 49.54 | 8.265 | -1.16 | 3.69E-02 | 8.32E-02 |
| CHMP6 | 9.979 | 2.198 | 10.82 | 3.172 | -1.204 | 5.01E-02 | 1.06E-01 |
| CHMP7 | 12.267 | 1.279 | 14.83 | 2.846 | 1.049 | 4.83E-01 | 6.09E-01 |
| DNM1L | 10.741 | 5.764 | 13.24 | 5.929 | 1.076 | 3.58E-01 | 4.87E-01 |
| FADD | 2.012 | 0.571 | 3.39 | 1.195 | -1.008 | 9.38E-01 | 9.59E-01 |
| FAF1 | 13.778 | 2.354 | 14.45 | 3.048 | -1.094 | 1.79E-01 | 2.88E-01 |
| H2AFJ | 23.487 | 12.103 | 28.24 | 5.017 | -1.072 | 5.05E-01 | 6.29E-01 |
| H2AFV | 45.833 | 5.492 | 50.24 | 16.493 | -1.144 | 4.75E-02 | 1.02E-01 |
| H2AFY2 | 7.106 | 1.717 | 8.22 | 2.965 | 1.027 | 7.73E-01 | 8.47E-01 |
| H2AFZ | 66.695 | 14.228 | 87.1 | 33.014 | 1.011 | 9.07E-01 | 9.41E-01 |
| HIST1H2AA | 0.553 | 0.548 | 0.32 | 0.73 | -1.878 | 7.62E-02 | 1.48E-01 |
| HIST1H2AC | 19.267 | 9.07 | 21.55 | 8.947 | -1.107 | 3.07E-01 | 4.34E-01 |
| HIST3H2A | 0.552 | 0.271 | 0.89 | 0.755 | 1.612 | 4.08E-02 | 9.04E-02 |
| HMGB1 | 84.561 | 15.898 | 124.52 | 47.187 | 1.014 | 8.59E-01 | 9.09E-01 |
| HSP90AA1 | 201.393 | 41.343 | 211.47 | 31.105 | -1.081 | 5.05E-01 | 6.30E-01 |
| HSP90AB1 | 200.887 | 27.098 | 224.47 | 30.98 | -1.106 | 3.57E-01 | 4.86E-01 |
| IL33 | 3.058 | 1.263 | 5.24 | 2.908 | 1.305 | 1.61E-01 | 2.64E-01 |
| JMJD7-PLA2G | 4.796 | 2.351 | 5.89 | 2.547 | 1.251 | 1.36E-01 | 2.32E-01 |
| MAPK8 | 5.502 | 1.404 | 6.1 | 1.566 | -1.112 | 1.43E-01 | 2.41E-01 |
| MAPK9 | 7.312 | 2.103 | 9.27 | 5.55 | 1.012 | 8.52E-01 | 9.03E-01 |
| PARP2 | 9.71 | 2.15 | 10.48 | 4.212 | -1.156 | 5.28E-02 | 1.11E-01 |
| PARP3 | 17.246 | 2.755 | 18.5 | 6.055 | -1.062 | 4.52E-01 | 5.80E-01 |
| PGAM5 | 9.477 | 1.657 | 9.12 | 0.892 | -1.187 | 5.38E-02 | 1.12E-01 |
| PLA2G4B | 0 | 4.418 | 0 | 1.106 | 1.318 | 1.95E-01 | 3.06E-01 |
| PPIA | 730.253 | 183.715 | 867.97 | 128.977 | -1.158 | 2.23E-01 | 3.41E-01 |
| PYGB | 23.443 | 5.497 | 27.04 | 6.713 | -1.009 | 9.14E-01 | 9.46E-01 |
| PYGM | 1.38 | 0.569 | 1.48 | 0.35 | 1.217 | 3.39E-01 | 4.68E-01 |
| RBCK1 | 29.842 | 7.831 | 38.47 | 15.353 | 1.144 | 1.20E-01 | 2.11E-01 |
| SPATA2 | 3.615 | 0.74 | 3.95 | 1.01 | -1.119 | 1.51E-01 | 2.52E-01 |
| SQSTM1 | 119.236 | 30.081 | 150.93 | 62.891 | -1.04 | 7.45E-01 | 8.28E-01 |
| STAT5B | 15.131 | 2.162 | 17.05 | 2.938 | 1.041 | 5.56E-01 | 6.74E-01 |
| TNFRSF10D | 4.075 | 0.882 | 4.58 | 1.664 | 1.017 | 8.62E-01 | 9.10E-01 |
| TNFRSF1A | 38.985 | 18.555 | 41.17 | 48.357 | 1.255 | 2.29E-02 | 5.64E-02 |
| TRADD | 24.781 | 7.056 | 26.89 | 7.645 | 1.001 | 9.89E-01 | 9.93E-01 |
| TRPM7 | 6.201 | 3.756 | 9.94 | 5.167 | -1.038 | 6.93E-01 | 7.88E-01 |
| TYK2 | 26.556 | 31.949 | 29.6 | 20.508 | 1.152 | 8.02E-02 | 1.54E-01 |
| USP21 | 14.509 | 3.148 | 15.72 | 3.219 | -1.104 | 1.30E-01 | 2.25E-01 |
| XIAP | 6.279 | 1.613 | 7.87 | 2.009 | 1.148 | 5.13E-02 | 1.08E-01 |

Supplemental Table S12. Expression of Core Matrisome Genes in TCMR and Normal Kidney Allograft Biopsies.

Table S12A. Collagens

| Symbol | Normal Allograft Median | Normal Allograft IQR | TCMR Median | TCMR IQR | FC | PValue | FDR |
|---|---|---|---|---|---|---|---|
| COL8A2 | 0.318 | 0.203 | 1.251 | 0.731 | 3.901 | 4.23E-13 | 1.90E-11 |
| **COL7A1** | **2.883** | **1.344** | **10.027** | **9.185** | **2.656** | **3.34E-07** | **4.41E-06** |
| COL6A3 | 8.748 | 6.572 | 21.495 | 13.720 | 2.466 | 3.41E-07 | 4.49E-06 |
| COL11A1 | 0.221 | 0.287 | 0.209 | 0.327 | 5.847 | 1.61E-06 | 1.75E-05 |
| **COL4A1** | **21.617** | **15.731** | **50.183** | **16.900** | **1.957** | **2.22E-06** | **2.34E-05** |
| COL17A1 | 0.094 | 0.058 | 0.196 | 0.184 | 2.681 | 7.30E-06 | 6.66E-05 |
| COL8A1 | 1.720 | 1.369 | 4.766 | 5.691 | 2.699 | 1.03E-05 | 9.00E-05 |
| COL5A2 | 3.210 | 3.427 | 7.781 | 6.925 | 2.001 | 7.54E-05 | 4.94E-04 |
| **COL3A1** | **58.603** | **38.986** | **126.735** | **144.793** | **2.419** | **1.26E-04** | **7.59E-04** |
| **COL1A1** | **23.169** | **19.658** | **40.230** | **75.681** | **2.869** | **1.36E-04** | **8.07E-04** |
| COL15A1 | 4.889 | 2.604 | 8.985 | 5.107 | 1.634 | 2.47E-04 | 1.35E-03 |
| **COL4A2** | **37.034** | **19.922** | **68.099** | **18.808** | **1.595** | **3.40E-04** | **1.77E-03** |
| COL5A1 | 3.565 | 4.419 | 9.721 | 8.426 | 2.029 | 3.54E-04 | 1.84E-03 |
| COL10A1 | 0.142 | 0.101 | 0.271 | 0.619 | 2.535 | 4.19E-03 | 1.43E-02 |
| **COL1A2** | **60.082** | **26.101** | **98.108** | **129.351** | **1.850** | **7.05E-03** | **2.18E-02** |
| COL16A1 | 7.219 | 4.632 | 13.255 | 10.556 | 1.492 | 1.25E-02 | 3.46E-02 |
| COL11A2 | 0.857 | 0.273 | 0.546 | 0.219 | -1.756 | 7.81E-05 | 5.09E-04 |
| COL19A1 | 0.611 | 0.445 | 0.387 | 0.534 | -1.895 | 1.27E-02 | 3.51E-02 |
| COL9A1 | 0.389 | 0.245 | 0.215 | 0.230 | -1.750 | 2.40E-02 | 5.87E-02 |
| COL4A4 | 4.815 | 4.506 | 7.831 | 4.133 | 1.254 | 5.75E-02 | 1.18E-01 |
| COL14A1 | 7.136 | 4.896 | 12.094 | 13.092 | 1.521 | 7.84E-02 | 1.51E-01 |
| COL6A5 | 0.027 | 0.019 | 0.071 | 0.035 | 1.567 | 1.01E-01 | 1.85E-01 |
| COL25A1 | 0.279 | 0.570 | 0.258 | 1.067 | 1.697 | 1.17E-01 | 2.08E-01 |
| COL6A2 | 35.491 | 16.190 | 48.469 | 17.892 | 1.207 | 1.71E-01 | 2.77E-01 |
| COL6A1 | 58.790 | 16.823 | 53.910 | 24.725 | -1.162 | 2.22E-01 | 3.39E-01 |
| COL5A3 | 1.109 | 1.466 | 1.901 | 1.446 | 1.259 | 2.23E-01 | 3.41E-01 |
| COL12A1 | 9.269 | 6.625 | 12.821 | 11.078 | 1.230 | 2.41E-01 | 3.62E-01 |
| COL13A1 | 1.131 | 0.748 | 1.485 | 0.567 | 1.177 | 2.84E-01 | 4.11E-01 |
| COL2A1 | 0.090 | 0.133 | 0.099 | 0.154 | 1.425 | 2.84E-01 | 4.11E-01 |
| COL27A1 | 14.276 | 12.609 | 22.488 | 10.121 | 1.138 | 3.08E-01 | 4.35E-01 |
| COL21A1 | 0.731 | 0.593 | 1.004 | 0.903 | 1.175 | 4.25E-01 | 5.54E-01 |
| COL22A1 | 0.652 | 0.841 | 0.872 | 0.779 | 1.157 | 5.21E-01 | 6.43E-01 |
| COL18A1 | 54.965 | 17.846 | 73.653 | 22.345 | 1.072 | 5.30E-01 | 6.51E-01 |
| COL28A1 | 2.201 | 2.446 | 2.890 | 2.491 | 1.099 | 6.10E-01 | 7.20E-01 |
| COL4A3 | 9.584 | 8.363 | 13.557 | 9.325 | 1.061 | 6.33E-01 | 7.39E-01 |
| COL9A2 | 6.076 | 3.901 | 7.439 | 9.007 | 1.075 | 6.44E-01 | 7.48E-01 |
| COL4A6 | 1.036 | 0.953 | 0.826 | 1.294 | 1.060 | 8.00E-01 | 8.67E-01 |
| COL6A6 | 0.075 | 0.082 | 0.071 | 0.082 | -1.040 | 8.59E-01 | 9.09E-01 |
| COL9A3 | 1.137 | 3.395 | 1.752 | 4.813 | -1.039 | 8.74E-01 | 9.19E-01 |
| COL24A1 | 0.226 | 0.351 | 0.357 | 0.170 | 1.030 | 9.10E-01 | 9.43E-01 |
| COL4A5 | 4.541 | 3.344 | 3.709 | 5.245 | 1.002 | 9.90E-01 | 9.94E-01 |
| COL23A1 | 1.563 | 0.637 | 1.851 | 0.762 | 1.000 | 1.00E+00 | 1.00E+00 |

Table S12B. Glycoproteins

| Symbol | Normal Allograft Median | Normal Allograft IQR | TCMR Median | TCMR IQR | FC | PValue | FDR |
|---|---|---|---|---|---|---|---|
| EMILIN2 | 1.566 | 0.555 | 5.576 | 3.792 | 4.498 | 4.22E-15 | 3.29E-13 |
| COMP | 0.688 | 0.446 | 1.627 | 5.034 | 22.593 | 1.53E-10 | 4.11E-09 |
| ZPLD1 | 0.099 | 0.071 | 0.387 | 0.364 | 5.755 | 2.53E-10 | 6.49E-09 |
| NELL2 | 0.543 | 0.350 | 2.012 | 3.760 | 5.290 | 1.52E-09 | 3.34E-08 |
| LRG1 | 0.000 | 0.000 | 2.485 | 1.670 | 3.207 | 6.59E-09 | 1.28E-07 |
| **TGFBI** | **18.575** | **13.653** | **58.767** | **45.536** | **2.686** | **1.40E-08** | **2.55E-07** |
| DDX26B | 2.010 | 0.882 | 4.767 | 1.829 | 1.772 | 2.35E-08 | 4.07E-07 |
| FGL2 | 19.707 | 8.389 | 39.567 | 23.608 | 1.965 | 2.97E-08 | 4.98E-07 |
| LAMC2 | 1.488 | 1.631 | 6.780 | 5.689 | 2.932 | 1.11E-07 | 1.63E-06 |
| CTHRC1 | 1.156 | 1.059 | 4.280 | 5.448 | 4.000 | 2.43E-07 | 3.32E-06 |
| THBS2 | 3.732 | 3.298 | 9.749 | 4.427 | 2.214 | 7.37E-07 | 8.81E-06 |
| **SPON2** | **8.524** | **4.406** | **23.657** | **28.538** | **2.474** | **9.18E-07** | **1.07E-05** |
| THBS4 | 0.930 | 0.679 | 0.898 | 1.084 | 5.619 | 1.01E-06 | 1.16E-05 |
| **FBN1** | **4.852** | **4.804** | **12.562** | **10.230** | **2.124** | **1.23E-06** | **1.38E-05** |
| **FBN2** | **0.071** | **0.058** | **0.328** | **0.275** | **3.523** | **1.71E-06** | **1.84E-05** |
| LAMA3 | 0.743 | 0.667 | 2.153 | 1.940 | 2.122 | 2.15E-06 | 2.27E-05 |
| LAMB3 | 2.427 | 1.184 | 3.939 | 2.917 | 1.892 | 9.00E-06 | 7.98E-05 |
| PXDN | 3.454 | 2.418 | 6.332 | 1.908 | 1.643 | 1.65E-05 | 1.35E-04 |
| NTNG2 | 0.157 | 0.110 | 0.402 | 0.245 | 2.623 | 2.51E-05 | 1.93E-04 |
| CILP2 | 0.099 | 0.121 | 0.086 | 0.248 | 6.801 | 4.79E-05 | 3.36E-04 |
| TNFAIP6 | 0.127 | 0.127 | 0.361 | 0.779 | 3.633 | 6.70E-05 | 4.49E-04 |
| VWF | 3.707 | 1.933 | 7.555 | 8.869 | 2.067 | 2.19E-04 | 1.21E-03 |
| MATN3 | 0.560 | 0.292 | 1.086 | 0.362 | 1.903 | 3.06E-04 | 1.62E-03 |
| LAMA4 | 2.143 | 1.460 | 4.350 | 1.654 | 1.507 | 4.50E-04 | 2.25E-03 |
| **FN1** | **49.994** | **27.544** | **80.029** | **75.700** | **1.871** | **6.27E-04** | **2.97E-03** |
| GLDN | 0.234 | 0.186 | 0.362 | 0.528 | 2.407 | 6.34E-04 | 2.99E-03 |
| CILP | 0.753 | 0.456 | 1.467 | 1.562 | 2.319 | 6.60E-04 | 3.10E-03 |
| LAMC1 | 8.579 | 6.143 | 14.454 | 6.040 | 1.442 | 7.62E-04 | 3.49E-03 |
| VWA5A | 7.456 | 1.864 | 11.856 | 2.526 | 1.298 | 1.28E-03 | 5.35E-03 |
| THBS1 | 23.284 | 8.944 | 32.766 | 19.351 | 1.514 | 1.46E-03 | 5.97E-03 |
| EMILIN1 | 13.260 | 4.355 | 20.356 | 13.309 | 1.548 | 1.93E-03 | 7.53E-03 |
| FGG | 0.140 | 0.274 | 0.314 | 0.876 | 4.770 | 2.07E-03 | 7.95E-03 |
| SPON1 | 3.189 | 5.992 | 9.640 | 8.073 | 1.767 | 3.20E-03 | 1.14E-02 |
| **EFEMP1** | **32.030** | **14.892** | **62.146** | **30.018** | **1.377** | **3.92E-03** | **1.35E-02** |
| CYR61 | 10.178 | 6.859 | 17.534 | 16.784 | 1.569 | 4.08E-03 | 1.40E-02 |
| WISP1 | 2.334 | 1.771 | 3.669 | 1.385 | 1.497 | 7.04E-03 | 2.17E-02 |
| FGA | 0.120 | 0.463 | 0.308 | 1.473 | 3.038 | 8.17E-03 | 2.45E-02 |
| VWDE | 0.077 | 0.086 | 0.158 | 0.283 | 1.966 | 8.50E-03 | 2.53E-02 |
| SVEP1 | 4.207 | 1.765 | 8.179 | 5.810 | 1.550 | 9.47E-03 | 2.75E-02 |
| AGRN | 29.514 | 6.838 | 43.421 | 22.583 | 1.407 | 9.69E-03 | 2.80E-02 |
| MFAP3 | 4.691 | 3.364 | 7.424 | 3.071 | 1.261 | 1.00E-02 | 2.88E-02 |
| CRISPLD2 | 4.722 | 2.899 | 7.383 | 10.684 | 1.613 | 1.34E-02 | 3.68E-02 |
| SRPX2 | 1.074 | 0.791 | 3.317 | 2.943 | 1.800 | 1.65E-02 | 4.34E-02 |

| Symbol | Normal Allograft Median | IQR | TCMR Median | IQR | FC | PValue | FDR |
|---|---|---|---|---|---|---|---|
| IGFBP4 | 339.368 | 178.861 | 197.059 | 98.102 | -1.862 | 3.38E-04 | 1.76E-03 |
| TNN | 0.297 | 0.138 | 0.160 | 0.157 | -2.309 | 6.44E-04 | 3.04E-03 |
| IGFBP3 | 55.935 | 22.456 | 47.402 | 24.215 | -1.538 | 7.31E-04 | 3.37E-03 |
| ZP2 | 0.141 | 0.137 | 0.082 | 0.089 | -3.009 | 1.26E-03 | 5.27E-03 |
| HMCN2 | 3.417 | 1.615 | 2.975 | 2.884 | -1.777 | 1.93E-03 | 7.52E-03 |
| VWA7 | 7.116 | 2.204 | 4.951 | 3.267 | -1.421 | 3.18E-03 | 1.13E-02 |
| EMILIN3 | 3.220 | 0.773 | 3.143 | 1.317 | -1.420 | 7.05E-03 | 2.18E-02 |
| SLIT2 | 5.121 | 2.870 | 3.315 | 2.563 | -1.801 | 8.41E-03 | 2.50E-02 |
| EGFLAM | 3.249 | 4.496 | 3.059 | 1.861 | -1.813 | 1.00E-02 | 2.88E-02 |
| NELL1 | 6.223 | 7.192 | 2.290 | 4.542 | -1.917 | 1.32E-02 | 3.62E-02 |
| ECM2 | 9.242 | 4.616 | 9.136 | 4.377 | -1.356 | 1.52E-02 | 4.06E-02 |
| FBN3 | 0.710 | 0.516 | 0.496 | 0.408 | -1.765 | 1.58E-02 | 4.18E-02 |
| POMZP3 | 6.442 | 3.731 | 5.778 | 3.433 | -1.417 | 1.65E-02 | 4.33E-02 |
| GAS6 | 48.201 | 10.643 | 45.009 | 16.010 | -1.232 | 2.76E-02 | 6.58E-02 |
| FBLN5 | 93.150 | 52.038 | 84.266 | 44.590 | -1.354 | 2.82E-02 | 6.69E-02 |
| NTNG1 | 1.633 | 1.505 | 0.669 | 1.191 | -1.934 | 2.83E-02 | 6.71E-02 |
| PCOLCE2 | 13.364 | 12.480 | 2.592 | 11.518 | -2.322 | 2.86E-02 | 6.77E-02 |
| SPP1 | 958.241 | 481.048 | 1822.545 | 1419.708 | 1.518 | 3.28E-02 | 7.56E-02 |
| NDNF | 3.510 | 2.539 | 1.179 | 3.291 | -2.024 | 3.41E-02 | 7.81E-02 |
| NID1 | 10.835 | 6.827 | 14.185 | 9.434 | 1.288 | 3.56E-02 | 8.10E-02 |
| TINAGL1 | 48.360 | 16.739 | 67.270 | 24.905 | 1.229 | 3.89E-02 | 8.70E-02 |
| POSTN | 7.653 | 8.038 | 8.706 | 27.968 | 1.916 | 4.11E-02 | 9.07E-02 |
| CRELD1 | 21.869 | 5.399 | 21.364 | 3.524 | -1.162 | 4.24E-02 | 9.31E-02 |
| RELN | 0.233 | 0.259 | 0.467 | 0.404 | 1.616 | 4.94E-02 | 1.05E-01 |
| RSPO4 | 0.380 | 0.379 | 0.335 | 0.479 | -1.646 | 4.96E-02 | 1.05E-01 |
| OIT3 | 1.358 | 0.606 | 0.678 | 0.973 | -1.874 | 5.37E-02 | 0.112033 |
| HMCN1 | 0.867 | 1.727 | 3.167 | 2.455 | 1.460 | 5.40E-02 | 0.112579 |
| NOV | 2.113 | 3.618 | 3.804 | 6.807 | 1.588 | 5.57E-02 | 0.115448 |
| TINAG | 64.326 | 25.622 | 36.004 | 32.913 | -1.791 | 5.63E-02 | 0.116324 |
| EMID1 | 7.414 | 3.577 | 6.182 | 3.757 | -1.343 | 6.19E-02 | 0.12572 |
| MXRA5 | 2.208 | 2.156 | 4.010 | 4.586 | 1.387 | 6.35E-02 | 0.128264 |
| NPNT | 27.646 | 9.513 | 20.025 | 10.311 | -1.215 | 6.63E-02 | 0.132658 |
| LAMB2 | 146.276 | 34.802 | 118.538 | 51.332 | -1.229 | 6.90E-02 | 0.136943 |
| MFAP2 | 3.561 | 3.257 | 7.553 | 4.683 | 1.389 | 7.44E-02 | 0.145233 |
| LAMA2 | 1.495 | 0.968 | 2.605 | 0.747 | 1.267 | 9.31E-02 | 0.173556 |
| CRELD2 | 11.455 | 3.431 | 15.541 | 7.394 | 1.162 | 9.32E-02 | 0.173744 |
| VWA9 | 12.845 | 1.364 | 13.641 | 1.907 | -1.115 | 9.72E-02 | 0.17953 |
| FBLN2 | 3.562 | 2.312 | 6.427 | 4.537 | 1.356 | 9.81E-02 | 0.180766 |
| ADIPOQ | 0.041 | 0.037 | 0.059 | 0.213 | 2.906 | 9.83E-02 | 0.181017 |
| NTN4 | 39.553 | 17.364 | 49.189 | 34.156 | 1.251 | 1.02E-01 | 0.186966 |
| NID2 | 5.525 | 4.139 | 8.995 | 4.487 | 1.215 | 0.106214 | 0.192169 |
| EDIL3 | 1.597 | 4.380 | 3.072 | 6.753 | 1.480 | 0.114053 | 0.202788 |
| LAMB4 | 0.537 | 0.262 | 0.552 | 0.171 | 1.437 | 0.123767 | 0.216046 |
| MMRN1 | 0.521 | 0.314 | 0.916 | 0.675 | 1.601 | 0.126641 | 0.219874 |
| VWA2 | 0.141 | 0.472 | 0.444 | 0.947 | 1.573 | 0.127083 | 0.2205 |
| PAPLN | 24.433 | 9.732 | 33.291 | 19.723 | 1.238 | 0.143322 | 0.241638 |
| SBSPON | 4.244 | 1.874 | 6.443 | 5.217 | 1.206 | 0.145995 | 0.245287 |
| VWA3A | 0.203 | 0.173 | 0.435 | 0.359 | 1.407 | 0.146451 | 0.245951 |
| MGP | 316.015 | 206.469 | 444.569 | 466.647 | 1.289 | 0.149304 | 0.24972 |
| SPARC | 174.705 | 56.776 | 344.513 | 326.367 | 1.251 | 0.163621 | 0.268301 |

| Symbol | Normal Allograft Median | Normal Allograft IQR | TCMR Median | TCMR IQR | FC | PValue | FDR |
|---|---|---|---|---|---|---|---|
| IGSF10 | 0.370 | 0.453 | 0.619 | 0.415 | 1.349 | 0.166996 | 0.272141 |
| LGI4 | 1.518 | 0.883 | 1.531 | 1.209 | -1.243 | 0.171674 | 0.278132 |
| ABI3BP | 6.910 | 4.193 | 11.708 | 5.989 | 1.198 | 0.179626 | 0.28808 |
| DPT | 7.586 | 6.577 | 12.002 | 9.589 | 1.374 | 0.182019 | 0.291092 |
| RSPO1 | 4.818 | 5.582 | 2.324 | 4.470 | 1.512 | 0.183884 | 0.29333 |
| FNDC1 | 0.603 | 0.654 | 0.569 | 1.603 | 1.465 | 0.186642 | 0.296746 |
| VWCE | 1.106 | 0.500 | 1.816 | 1.382 | 1.279 | 0.187693 | 0.298086 |
| LTBP4 | 29.694 | 12.815 | 24.336 | 20.212 | -1.219 | 0.190408 | 0.30104 |
| RSPO3 | 0.907 | 0.763 | 1.219 | 1.009 | 1.423 | 0.19978 | 0.312599 |
| PCOLCE | 25.114 | 8.822 | 35.623 | 6.742 | 1.186 | 0.202687 | 0.31597 |
| NTN1 | 0.675 | 0.290 | 0.755 | 0.596 | -1.288 | 0.20642 | 0.32063 |
| MFAP5 | 0.828 | 2.218 | 0.993 | 4.635 | 1.747 | 0.21652 | 0.33282 |
| TNXB | 3.028 | 6.260 | 3.393 | 5.441 | 1.200 | 0.224037 | 0.341964 |
| TSKU | 27.619 | 19.146 | 13.274 | 13.466 | -1.425 | 0.233945 | 0.353889 |
| VWA3B | 0.127 | 0.158 | 0.223 | 0.165 | 1.427 | 0.237513 | 0.358557 |
| MFAP4 | 36.869 | 34.142 | 55.360 | 21.058 | -1.235 | 0.243053 | 0.364869 |
| LGI2 | 0.789 | 0.714 | 0.985 | 2.834 | 1.406 | 0.243308 | 0.365085 |
| PXDNL | 0.774 | 1.587 | 0.447 | 1.134 | -1.516 | 0.267579 | 0.392233 |
| AEBP1 | 47.105 | 20.774 | 61.039 | 33.726 | 1.177 | 0.278762 | 0.404464 |
| SMOC2 | 4.109 | 2.741 | 6.274 | 4.617 | 1.231 | 0.280099 | 0.406152 |
| SMOC1 | 0.494 | 1.022 | 0.666 | 0.618 | -1.336 | 0.285264 | 0.41169 |
| LAMA5 | 18.571 | 6.703 | 19.628 | 13.829 | 1.122 | 0.297351 | 0.424221 |
| VWA5B1 | 0.717 | 0.449 | 0.575 | 0.575 | -1.264 | 0.304279 | 0.431027 |
| COCH | 0.000 | 1.798 | 0.000 | 0.000 | -1.277 | 0.336346 | 0.46507 |
| FRAS1 | 7.791 | 3.178 | 7.453 | 5.644 | -1.124 | 0.348974 | 0.478012 |
| TECTA | 0.286 | 0.050 | 0.295 | 0.108 | -1.120 | 0.374485 | 0.504436 |
| MFGE8 | 31.035 | 20.180 | 41.332 | 12.115 | -1.101 | 0.394639 | 0.524294 |
| FBLN7 | 1.073 | 0.587 | 1.566 | 0.608 | 1.136 | 0.446862 | 0.575109 |
| THSD4 | 2.587 | 3.527 | 2.693 | 3.197 | 1.149 | 0.469995 | 0.597052 |
| LAMA1 | 3.439 | 4.375 | 4.237 | 4.966 | 1.146 | 0.47468 | 0.60086 |
| LGI3 | 0.190 | 0.113 | 0.165 | 0.130 | -1.259 | 0.478215 | 0.604359 |
| SRPX | 2.125 | 1.434 | 2.438 | 2.818 | 1.152 | 0.48355 | 0.609779 |
| IGFBP5 | 140.591 | 70.929 | 146.346 | 79.650 | -1.123 | 0.48463 | 0.610906 |
| IGFBP1 | 0.234 | 0.254 | 0.111 | 0.230 | -1.316 | 0.507293 | 0.631515 |
| MMRN2 | 11.137 | 9.663 | 14.375 | 9.702 | 1.087 | 0.516849 | 0.639416 |
| IGFBP2 | 47.656 | 26.466 | 41.313 | 50.283 | -1.124 | 0.527484 | 0.648849 |
| OTOG | 0.072 | 0.063 | 0.109 | 0.091 | -1.169 | 0.54028 | 0.659832 |
| ELN | 14.349 | 7.662 | 14.032 | 6.176 | 1.118 | 0.546118 | 0.664436 |
| FBLN1 | 32.930 | 20.958 | 40.950 | 22.986 | -1.101 | 0.556467 | 0.673873 |
| ZP3 | 3.374 | 3.349 | 1.292 | 2.639 | -1.097 | 0.556832 | 0.674266 |
| COLQ | 0.000 | 2.482 | 0.000 | 0.000 | 1.070 | 0.560433 | 0.677825 |
| LTBP1 | 6.494 | 11.103 | 9.407 | 15.596 | 1.182 | 0.567623 | 0.683844 |
| CRIM1 | 23.441 | 15.441 | 25.955 | 10.849 | -1.047 | 0.692058 | 0.787155 |
| CRISPLD1 | 0.796 | 0.558 | 0.840 | 1.030 | -1.060 | 0.716238 | 0.80626 |
| IGFBP7 | 848.833 | 123.831 | 960.944 | 237.182 | -1.045 | 0.727632 | 0.815188 |
| LTBP2 | 3.692 | 1.663 | 4.025 | 1.892 | 1.037 | 0.771648 | 0.845962 |
| VWA1 | 27.078 | 6.733 | 29.649 | 9.585 | -1.027 | 0.779773 | 0.851792 |
| LTBP3 | 29.503 | 16.677 | 38.752 | 14.634 | 1.032 | 0.783757 | 0.854889 |
| LAMC3 | 13.081 | 6.134 | 11.935 | 11.174 | -1.049 | 0.798005 | 0.865333 |
| FGB | 2.365 | 4.851 | 3.553 | 12.923 | 1.113 | 0.836939 | 0.893094 |
| IGFBP6 | 21.982 | 12.687 | 25.606 | 18.604 | 1.037 | 0.846064 | 0.899433 |
| MATN2 | 15.377 | 9.176 | 21.458 | 20.613 | -1.030 | 0.85834 | 0.908182 |
| LAMB1 | 56.066 | 49.409 | 79.510 | 33.667 | 1.023 | 0.860694 | 0.909497 |
| ECM1 | 6.049 | 1.977 | 7.034 | 4.069 | 1.021 | 0.886688 | 0.927242 |
| SNED1 | 2.187 | 1.752 | 3.925 | 3.111 | 1.011 | 0.915252 | 0.946631 |
| SLIT3 | 8.830 | 5.651 | 14.004 | 6.184 | 1.016 | 0.915783 | 0.946881 |
| SPARCL1 | 34.058 | 14.681 | 36.972 | 32.038 | 1.015 | 0.926462 | 0.952691 |
| CTGF | 26.284 | 27.674 | 36.804 | 23.396 | 1.014 | 0.927468 | 0.953247 |
| THBS3 | 12.657 | 6.333 | 16.728 | 7.150 | -1.007 | 0.933413 | 0.956717 |
| WISP2 | 0.348 | 1.042 | 0.568 | 3.020 | 1.028 | 0.945201 | 0.964454 |
| VIT | 0.089 | 0.129 | 0.189 | 0.256 | 1.018 | 0.958716 | 0.973092 |
| MFAP1 | 13.531 | 1.842 | 16.140 | 2.422 | -1.001 | 0.982743 | 0.988597 |
| BMPER | 0.250 | 0.291 | 0.251 | 0.220 | 1.001 | 0.994597 | 0.99697 |

Table S12C. Proteoglycans

| Symbol | Normal Allograft Median | Normal Allograft IQR | TCMR Median | TCMR IQR | FC | PValue | FDR |
|---|---|---|---|---|---|---|---|
| **SRGN** | **21.264** | **6.983** | **75.969** | **74.474** | **3.604** | **1.80E-15** | **1.53E-13** |
| VCAN | 1.729 | 1.644 | 13.158 | 14.912 | 4.844 | 5.84E-13 | 2.55E-11 |
| **HAPLN3** | **1.482** | **1.182** | **7.840** | **11.195** | **6.391** | **1.67E-12** | **6.72E-11** |
| EPYC | 0.023 | 0.039 | 0.058 | 0.190 | 16.740 | 2.04E-05 | 1.63E-04 |
| LUM | 134.858 | 41.241 | 247.852 | 162.656 | 1.637 | 2.38E-03 | 8.95E-03 |
| HAPLN1 | 0.052 | 0.134 | 0.216 | 0.456 | 3.485 | 9.93E-03 | 0.02863 |
| BCAN | 0.104 | 0.098 | 0.075 | 0.158 | 2.474 | 1.05E-02 | 3.00E-02 |
| ESM1 | 4.216 | 3.909 | 3.397 | 2.287 | -1.762 | 2.13E-03 | 8.17E-03 |
| NYX | 0.356 | 0.193 | 0.255 | 0.286 | -2.057 | 4.16E-03 | 1.42E-02 |
| FMOD | 4.754 | 4.560 | 10.255 | 5.832 | 1.483 | 3.85E-02 | 0.086126 |
| IMPG2 | 0.059 | 0.133 | 0.115 | 0.138 | 1.592 | 8.78E-02 | 0.165627 |
| OGN | 3.021 | 4.149 | 3.441 | 4.288 | -1.547 | 0.189739 | 0.300388 |
| ACAN | 1.303 | 0.861 | 1.151 | 1.165 | -1.286 | 0.285386 | 0.411777 |
| PODN | 2.224 | 1.757 | 3.916 | 2.537 | 1.279 | 0.305987 | 0.432848 |
| HSPG2 | 22.092 | 8.347 | 28.977 | 11.410 | 1.163 | 0.315359 | 0.44295 |
| SPOCK1 | 4.383 | 1.277 | 4.256 | 2.666 | -1.066 | 0.661113 | 0.762819 |
| SPOCK3 | 0.118 | 0.113 | 0.151 | 0.207 | 1.098 | 0.794393 | 0.862242 |
| ASPN | 5.429 | 3.465 | 5.874 | 3.900 | 1.027 | 0.890563 | 0.929536 |
| BGN | 84.638 | 39.210 | 106.894 | 65.963 | -1.021 | 0.897229 | 0.934362 |
| SPOCK2 | 16.351 | 9.130 | 15.955 | 10.670 | -1.023 | 0.897921 | 0.934965 |
| DCN | 184.460 | 94.813 | 265.780 | 263.341 | 1.017 | 0.921 | 0.949877 |
| PRELP | 5.752 | 6.903 | 7.311 | 6.566 | -1.021 | 0.922008 | 0.950544 |
| IMPG1 | 0.418 | 0.473 | 0.487 | 0.487 | 1.017 | 0.927388 | 0.953225 |
| OMD | 0.635 | 0.394 | 0.948 | 0.826 | -1.020 | 0.929872 | 0.954819 |

#### Supplemental Table S13. 18 ECM Highly Connected Center Nodes Genes

| Type | Symbol | description | FC_TCMR | FDR | Gene Significance | Connectivity | Tissue/Blood Cells |
|---|---|---|---|---|---|---|---|
| Collagens | COL1A1 | collagen type I alpha 1 chain | 2.87 | 8.07E-04 | 0.456 | 16.65 | ECM connective tissue; Naïve CD8 T Cells |
| Collagens | COL17A1 | collagen type XVII alpha 1 chain | 2.68 | 6.66E-05 | 0.336 | 0.45 | Neutrophils, Monocytes |
| Collagens | COL3A1 | collagen type III alpha 1 chain | 2.42 | 7.59E-04 | 0.507 | 20.35 | ECM, Not detected in Immune Cells |
| Collagens | COL1A2 | collagen type I alpha 2 chain | 1.85 | 2.18E-02 | 0.542 | 25.50 | ECM connective tissue;not detected in Immune ce |
| Glycoproteins | FGG | fibrinogen gamma chain | 4.77 | 7.95E-03 | 0.315 | 0.73 | blood-borne glycoprotein (Liver) |
| Glycoproteins | FGA | fibrinogen alpha chain | 3.04 | 2.45E-02 | 0.437 | 1.08 | blood-borne glycoprotein (Liver) |
| Coagulation & P | ITGB3 | integrin subunit beta 3 | 2.91 | 7.04E-10 | 0.620 | 14.80 | cell-surface proteins; Baso and Neutro |
| Glycoproteins | VWF | von Willebrand factor | 2.07 | 1.21E-03 | 0.488 | 1.42 | low levels kidney; Eos, Neutro, Monocytes |
| Glycoproteins | FN1 | fibronectin 1 | 1.87 | 2.97E-03 | 0.541 | 12.87 | fibroblasts, epithelial and other cell types |
| Regulators | CTSS | cathepsin S | 8.34 | 5.52E-19 | 0.693 | 45.89 | High Tubules & Glomeruli medium; Monocytes, D |
| Regulators | MMP9 | matrix metallopeptidase 9 | 7.71 | 3.88E-08 | 0.686 | 10.37 | ECM, Neutrophils |
| Regulators | CTSG | cathepsin G | 4.44 | 4.90E-05 | 0.446 | 4.03 | Moncytes, DCs, NK Cell, Memory CD8 and naïve T C |
| Regulators | MMP25 | matrix metallopeptidase 25 | 2.88 | 1.69E-08 | 0.693 | 32.74 | ECM; Neutrophils, inactivates SERPINA1 |
| Regulators | TIMP1 | TIMP metallopeptidase inhibitor 1 | 2.79 | 2.92E-07 | 0.736 | 30.30 | ECM most tissues, Immune Cells |
| Regulators | ADAM8 | ADAM metallopeptidase domain 8 | 2.72 | 1.09E-09 | 0.733 | 36.63 | Neutrophils and Monocytes |
| Regulators | CTSK | cathepsin K | 1.94 | 1.60E-04 | 0.615 | 5.64 | Moncytes, DCs, NK Cells; renal tissue |
| Regulators | ADAM17 | ADAM metallopeptidase domain 17 | 1.47 | 1.67E-04 | 0.666 | 16.17 | Most tissues and immune cells |
| Secreted | TGFB1 | transforming growth factor beta 1 | 1.72 | 1.05E-05 | 0.714 | 32.68 | ECM most tissue, Immune cells |
